# Mouse peritoneal macrophages undergo female-specific remodeling with aging driven by both hormone-dependent and -independent mechanisms

**DOI:** 10.1101/2025.06.11.659200

**Authors:** Ryan J. Lu, Shiqi Chen, Minhoo Kim, Nirmal K. Sampathkumar, Evelyn H. Lee, Amy Christensen, Eric E. Wang, Isabella Y. Lau, Shivay Parihar, Charan K. Ravikumar, Jane Jung, Shelby Brown, Alan Xu, Julio L. Alvarenga, Kristen Mehalko, Changhan D. Lee, Helen S. Goodridge, Bérénice A. Benayoun

## Abstract

Aging is a complex process characterized by a progressive decline in physiological functions. Immune function is strongly influenced by biological sex, affecting both innate and adaptive responses. Here, we investigated the effects of age and sex on mouse peritoneal immune cells and identified macrophages as the top affected cell type. Macrophages, as central components of the innate immune system, play critical roles in homeostasis maintenance and infection response. We found that aging induces sex-specific remodeling of mouse peritoneal macrophage ‘omic’ landscapes, which is accompanied by female-specific age-related functional remodeling (*i.e*. decreased phagocysis, increased glycolysis). We show that age-related changes in circulating estrogen levels likely drive aspects of female-specific macrophage age-related changes, specifically age-related phagocytosis decline. By leveraging our multi-omic dataset, we identify transcription factors whose female-specific age-regulation may drive female-specific age-related changes in peritoneal macrophage phenotypes. Interestingly, *Irf2* downregulation was sufficient to recapitulate aspects of female peritoneal macrophage aging phenotypes, including increased glycolysis. Mechanistically, decreased Irf2 expression leads to decreased binding to its target genes, including *Hk3*, which encodes hexokinase (the rate-limiting enzyme of glycolysis). Our findings support the notion that aging leads to female-specific remodeling of mouse peritoneal macrophages through hormone-dependent and -independent mechanisms.

## Introduction

Aging is a complex process characterized by a progressive decline in physiological functions, with notable differences between sexes^1^. This decline is driven by a combination of biological factors and extrinsic factors^2^. Life expectancy of females is, on average, longer than that of males^3^. However, despite increased longevity, females tend to experience higher levels of comorbidities and disability than males (*i.e*. the mortality-morbidity paradox)^4–6^. Additionally, females face an elevated risk of developing age-related chronic conditions (*e.g*. coronary heart disease, osteoarthritis, Type 2 diabetes, neurodegenerative disorders)^7^. Chronic sterile inflammation, a mechanism dubbed “inflamm-aging”, is a hallmark of aging and is associated with more general immune dysfunction or “immunosenescence”^8,9^. Despite well-documented sex-differences in aging and disease susceptibility, mechanisms driving sex-based aging disparities are poorly understood.

Immune function is heavily shaped by biological sex, influencing both innate and adaptive immune cells^10,11^, including macrophages^12^. Females generally exhibit a more robust immune response compared to males (*e.g*. quicker pathogen clearance, higher antibody response, higher cytokine secretion, and less infection-related mortality)^10,11,13–15^. However, females are also more susceptible to autoimmune diseases^16^. During aging, hematopoietic stem cells (HSC) skew production toward myeloid cells (*e.g*. monocytes, neutrophils) at the expense of lymphoid cells (*e.g.* B- and T-cells), a shift more pronounced in males than females, thereby impacting leukemogenesis and hematopoiesis^17,18^. Importantly, lifelong sex differences in immune function can be shaped by differential ‘omic’ regulation in immune cell types^19–24^. Despite the potential significance of such sex-differences in immune aging, how biological sex shapes the impact of age on ‘omic’ and functional landscapes of innate immune cells is largely unexplored.

Macrophages are key components of the innate immune system, functioning as both regulatory and effector cells in maintaining tissue homeostasis and responding to infections. Tissue-resident macrophages (*e.g*. large peritoneal macrophages), which originate primarily from embryonic precursors, exhibit specialized, organ-specific functions shaped by the local microenvironment. In contrast, bone marrow-derived macrophages differentiate from circulating monocytes and typically adopt more pro-inflammatory phenotypes during immune responses^25,26^. Macrophages have a repertoire of functions: phagocytosis of pathogens and other foreign materials, clearance of dead/dying and senescent cells, chemotaxis and migration, and cytokine production^27^. During aging, macrophages have been reported to exhibit altered phagocytosis, metabolic changes, and cytokine secretion, leading to poor inflammation control^28–36^. In addition to age-related differences, sex-differences in young macrophages have also been reported^10^, where young females exhibited greater phagocytic activity and more efficient antigen presentation^37,38^. Interestingly, microglia, the brain’s resident macrophages, undergo a female-specific shift in glycolytic output with age dependent on C3a signaling^39^. While pioneering studies have started to unearth sex-differences in immune aging^40^, systematic characterization of such sex-dimorphic phenotypes in aging macrophages and the molecular mechanisms driving them are still missing.

In this study, we leverage a functional genomics approach to identify sex-independent and sex-specific phenotypes of primary peritoneal macrophages in female *vs*. male mice with aging, as well as understand potential mechanistic drivers of sex-specific aging phenotypes.

## Results

### Macrophages are the immune cell type most impacted by aging in the peritoneal cavity niche

The peritoneal cavity hosts a rich and accessible immune niche, housing B-cells, T-cells, macrophages, neutrophils, and monocytes^41^. This immune niche is easily accessible through post-mortem peritoneal lavage^42,43^, making it tractable for studies without the need for mechanical tissue dissociation (which compromises both cell viability and phenotypes^44^). We used this niche to examine how biological sex may impact the mice’s aging immune system. Specifically, we obtained peritoneal lavage cells from young adult (4-5 months) and old (20-21 months) female and male C57BL/6JNia and C57BL/6J mice (**Figure 1a**). Isolated cells were used for unbiased profiling by single-cell RNA sequencing (scRNA-seq) using the 10xGenomics v2 and v3.1 3’ platforms in 3 independent sets of aging mice (see **Methods**; **Figure 1a**).

**Figure 1.**
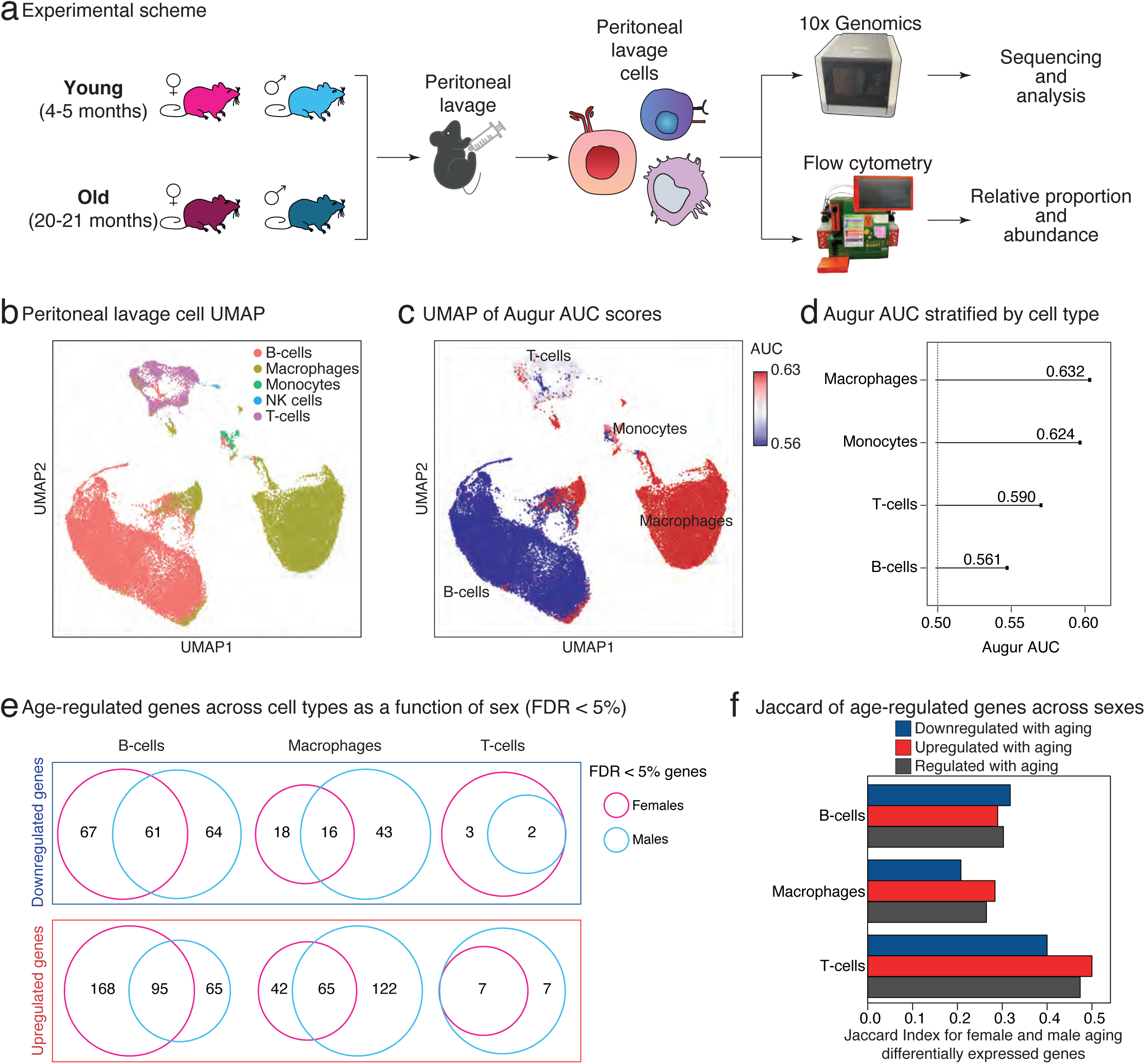
Peritoneal macrophages are the most impacted cell type in the peritoneal cavity niche as a function of age and biological sex. (**a**) Experimental scheme illustrating age-based cell profiling in mouse peritoneal lavage. (**b**) Uniform Manifold Approximation and Projection (UMAP) visualization of unsupervised clustering of peritoneal lavage cells. (**c**) UMAP representation of AUGUR area under the curve (AUC) scores for peritoneal lavage immune cell populations. (**d**) Lollipop plot of Augur AUC results stratified by immune cell types. (**e**) Venn diagrams for robust age-related genes in female vs. male B-cells, macrophages and T-cells (FDR < 5%). In each independent scRNA-seq dataset, data was pseudobulked by biological replicate and differential gene expression analysis was conducted. Robustly regulated genes were obtained using metaRNAseq (see Methods). Note limited overlap between age-regulated female vs. male genes. (**f**) Jaccard index analysis of overlapping robust age-related genes in female vs. male B-cells, macrophages and T-cells (FDR < 5%). A value of 1 indicates perfect overlap, a value of 0 indicates no overlap.

We robustly identified expected cell types within this niche in our dataset: B-cells, macrophages, monocytes, natural killer (NK) cells, and T-cells (**Figure 1b**; 18,805 cells, 10,758 cells, and 14,595 cells for the NIA 10x v2, NIA 10x v3 and JAX datasets, respectively). The scRNA-seq can be used to infer relative cell type abundances, especially in the absence of potential dissociation artifacts. Thus, we asked if there were any age- or sex-related changes in the relative abundances of peritoneal cell types, using both sample-specific determination and the scRNA-seq specific algorithm scProportionTest^45^ (**Extended Data Figure 1a-d**). Interestingly, both methods revealed a relative increase in the proportion of B-cells concomitant with a relative decrease in the proportion of macrophages with aging, regardless of sex (**Extended Data Figure 1e**), consistent with previous studies^46,47^. To note, we also observed a relative increase in the proportion of T-cells in the peritoneal niche only in aging males (**Extended Data Figure 1e**). To validate these results independently *in vivo*, we used flow cytometry staining of peritoneal cells using cell surface markers CD19^+^ (B-cells), F4/80^+^ (macrophages), and CD3^+^ (T-cells) (**Extended Data Figure 2a**). We confirmed the relative B-cells increase, and relative macrophages decrease with age in both sexes, but not the male-specific T-cells increase (**Extended Data Figure 2b**). Next, we accessed whether these relative proportional changes were accompanied by alterations in absolute abundances. Result revealed that the absolute numbers of all three cell types increased in the peritoneal cavity with aging, with B-cells showing the most pronounced increase, particularly in females (**Extended Data Figure 2c**).

We next investigated whether the transcriptional landscapes any of these cell types were significantly influenced by age and/or sex. For this purpose, we leveraged Augur^48^, an algorithm that helps prioritize cell types based on their transcriptional response to biological conditions using machine-learning (**Figure 1c,d**; **Extended Data Figure 3a-f**). Using Augur-derived area under the curve (AUC) scores, we observed that, among peritoneal cell types, macrophage transcriptomes were most impacted in both the combined (AUC = 0.632; **Figure 1d**) and individual datasets (AUC ≥ 0.567; **Extended Data Figure 3d, e, f**). Furthermore, we evaluated how similar the transcriptomic response to aging was in female *vs*. male peritoneal B-cells, macrophages, and T-cells (**Figure 1e-f**), using pseudobulked differential gene expression analysis across the 3 independent scRNA-seq datasets (see **Methods; Extended Data Table 2A-F**). Interestingly, there was only a modest overlap between robust age-regulated genes in females vs. males across the 3 cell types (**Figure 1e**), with Jaccard indeces < 0.5 indicating divergence in aging transcriptomes as a function of sex (**Figure 1f**). Consistent with our Augur results, age-related transcriptional changes in macrophages showed the lowest overlap between sexes of the 3 peritoneal cell types (**Figure 1f**), suggesting their aging may be most differentially impacted by sex. Thus, we hereafter decided to focus on the peritoneal macrophage compartment to understand potential sex-dimorphic molecular and functional effects with aging.

### Peritoneal macrophages show sex-dimorphic ‘omic’ phenotypes with aging

To better understand the impact of aging and sex on aging peritoneal macrophages, we performed a bulk multi-omic profiling approach using MACS-purified macrophages from young adults (4-5 months) and old (20-21 months) female and male C57BL/6JNia mice. We opted to use a bulk approach due to its greater sensitivity and robustness compared to single-cell profiling, particularly for well-defined purified cells. For this study, we profiled transcriptional and epigenomic features, specifically (i) transcriptome profiling by bulk sequencing of ribosomal-RNA depleted RNA (RNA-seq), (ii) chromatin accessibility landscape through the Assay for Transposon-Accessible Chromatin followed by sequencing (ATAC-seq)^49,50^, and (iii) promoter activity/priming using CUT&RUN profiling for the H3K4me3 histone post-translational modification^51^ (**Figure 2a**). These ’omic’ assays were each performed on an independent cohort of C57BL/6JNia animals (n = 5 animals per group; **Extended Data Table 1A**). After accounting for premature death before the experimental day (**Extended Data Table 1A**) and retaining only samples with > 90% macrophage purity (**Extended Data Figure 4a, b**), final sample numbers ranged from 3-5 per group and ‘omic’ layer.

**Figure 2.**
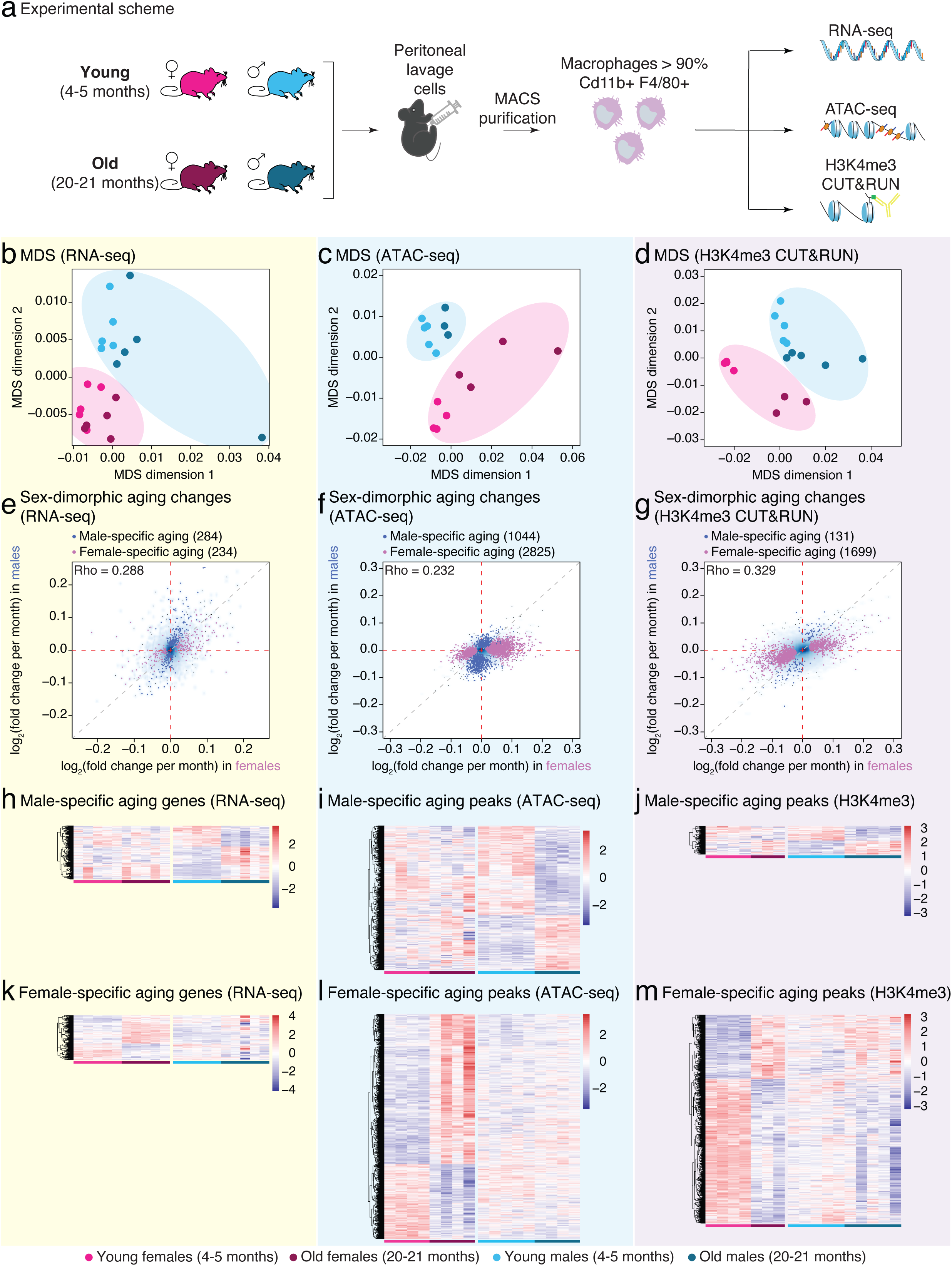
The peritoneal macrophage transcriptome and epigenome shows sex-dimorphic remodeling with aging. (**a**) Experimental scheme of murine peritoneal macrophage ‘omics’ profiling. (**b**) Multidimensional scaling (MDS) plot of transcriptomic profiles, indicating distinct transcriptomic aging patterns between sexes. (**c**) MDS plot of chromatin accessibility profiles by ATAC-seq. (**d**) MDS plot of H3K4me3 CUT&RUN profiles. (**e**) Scatterplot of age-related changes in peritoneal macrophage transcriptomes in female vs. males, reported as DESeq2 log_2_ fold change in gene expression per month. 284 Male-specific (blue) and 234 female-specific (pink) aging-regulated genes (Transcriptome-wide Spearman Rho = 0.288). (**f**) Scatterplot of age-related changes in peritoneal macrophage ATAC-seq accessible regions in female vs. males, reported as DESeq2 log_2_ fold change in gene expression per month. 1044 Male-specific (blue) and 2825 female-specific (pink) regions (Genome-wide Spearman Rho = 0.232). (**g**) Scatterplot of age-related changes in peritoneal macrophage H3K4me3 CUT&RUN signal in female vs. males, reported as DESeq2 log_2_ fold change in gene expression per month. 131 male-specific (blue) and 1699 female-specific (pink) marks (Genome-wide Spearman Rho = 0.329). (**h-j**) Heatmap of <u>male</u>-specific aging changes in gene expression by RNA-seq (**h**), in chromatin accessibility by ATAC-seq (**i**) H3K4me3 peak intensity by CUT&RUN (**j**). (**k-m**) Heatmap of <u>female</u>-specific aging changes in gene expression by RNA-seq (**k**), in chromatin accessibility by ATAC-seq (**l**) H3K4me3 peak intensity by CUT&RUN (**m**).

The peritoneal macrophage population could be comprised of both large peritoneal macrophages (LPM; with high surface levels of F4/80) and small peritoneal macrophages (SPM; with lower surface levels of F4/80)^52^. LPMs are the tissue-resident macrophages of the peritoneal cavity derived from the fetal yolk sac, and constitute the overwhelming majority of peritoneal macrophages in naïve mice^53–55^. In contrast, SPMs are constantly derived throughout life from blood monocytes, which are recruited to the peritoneal cavity in response to inflammatory signals, and represent a minority of peritoneal macrophages in naïve mice^55,56^. To rule out the possibility that age-related changes in purified peritoneal macrophages could be trivial byproducts of shifts in LPM/SPM relative abundances with aging, we analyzed F4/80 surface protein levels in MACS-purified peritoneal macrophages from C57BL/6JNia mice (**Extended Data Figure 4c**). The result indicates that LPMs remained the major macrophage type in the aging peritoneal cavity, regardless of sex. There were no significant changes in the relative proportions of LPMs *vs*. SPMs in aging C57BL/6JNia mice, regardless of sex (**Extended Data Figure 4d-e**). These analyses gave us confidence that any observed changes in bulk macrophage ‘omic’ profiles with aging would not be the mere result of a depletion of the tissue-resident LPM population in the aging peritoneal cavity.

We then leveraged Multi-Dimensional Scaling [MDS] to evaluate the main axes of separation of our samples across the 3 bulk ‘omic’ layers (**Figure 2b-d**). Interestingly, MDS revealed clear separation of samples by sex, regardless of age (**Figure 2b-d**). The MDS analysis supports the existence of strong sex-dimorphism in the molecular aging landscape of primary peritoneal macrophages. We also observed a secondary separation of samples by age of the mice, albeit weaker than that by sex (**Figure 2b-d**). Thus, as suggested by previous scRNA-seq analyses (**Figure 1e-f**), we asked whether aging influences the ‘omic’ landscapes of peritoneal macrophages in a sex-dimorphic manner. For every detected gene/peak in aging female *vs*. male peritoneal macrophages, we compared DESeq2 log_2_ fold changes in gene expression (**Figure 2e, Extended Data Figure 5a**), chromatin accessibility (**Figure 2f, Extended Data Figure 5b**) and H3K4me3 CUT&RUN signal (**Figure 2g, Extended Data Figure 5c**). We reasoned that high rank correlation values would correspond to largely colinear effects of aging on ‘omic’ landscapes regardless of sex, and divergent aging trajectories would result in weaker values. Using this method, we observed only relatively weak rank correlations between ‘omic’ aging trajectories of female and male peritoneal macrophages (Genome-wide Spearman Rho = 0.232-0.329; **Figure 2e-g, Extended Data Figure 5a-c**).

To decipher these divergent patterns, we identified significant sex-specific ‘omic’ changes. Specifically, we labeled genes with FDR < 5% in one sex, but FDR > 10% in the other sex, as significant in one sex only (sex-divergent regulation) and genes with FDR < 5% in both sexes as consistently regulated (sex-independent regulation; **Extended Data Table 2G-I**). Importantly, although a core set of genes and peaks significantly changed similarly with aging in both sexes (respectively 61, 420 and 244; **Extended Data Figure 5a-c**), there were many more genes and/or peaks with significant aging changes in only one sex (respectively 284/234, 1044/2825 and 131/1699 [male-specific/female-specific]; **Figure 2e-m**). Together, our analyses are consistent with the notion of broadly sex-divergent aging ‘omic’ trajectories in peritoneal macrophages.

In general, sex differences can result from (i) downstream effects of sex hormone signaling (*e.g*. lifelong exposure to an estrogen- or androgen-rich milieu, or (ii) differential impact of sex chromosome karyotypes (*e.g*. XX vs. XY), including through X chromosome inactivation [XCI] escape^40^. Genes and/or peaks with sex-dimorphic aging regulation were distributed across the genome, indicating that autosomal gene dysregulation remains a major feature of sex-dimorphism in the aging ‘omic’ landscapes of macrophages (**Extended Data Figure 5d-f**).

Intriguingly, *Xist*, which encodes a non-coding RNA critical to XCI initiation and maintenance^57,58^, was one of the top female-specific age-regulated genes in our bulk RNA-seq dataset (FDR = 5.4x10^-5^; **Extended Data Figure 6a**). Consistently, increased *Xist* expression was also observed in macrophages from our 10xGenomics v3.1 scRNA-seq datasets (but not v2 dataset; **Extended Data Figure 6b**). Upregulation of *Xist* has been reported across cell types in aging female brains, including in brain macrophages/microglia^59^. Moreover, increased XCI escape was recently reported across several female tissues with mouse aging^60,61^, which was proposed to underlie the unique female risk for autoimmunity^62^. Since we observed upregulation of *Xist* in aging female macrophages, we next asked whether there was any trend of regulation of X-linked genes and/or known XCI-escapees among female age-regulated genes in peritoneal macrophages. Gene set enrichment analysis [GSEA] revealed a slight, but significant, trend for downregulation of both X-linked genes (FDR = 1.4 x10^-2^) and known XCI escapees (FDR = 3.6x10^-3^) in aged female macrophages (**Extended Data Figure 6c**). This downregulation is consistent with dysregulation of XCI in aging female macrophages, albeit skewing towards potentially increased repression – in contrast with prior observations of increased escape in the aging brain^59,60^.

Together, our data reveals strong sex differences in the ‘omic’ landscapes of primary peritoneal macrophages throughout life and suggest that aging may differentially impact the function of these cells in female *vs.* male mice.

### Functional enrichment analysis reveals sex-dimorphic regulation of key immune pathways with aging

We next investigated whether sex-specific, age-related changes in peritoneal macrophage gene expression regulation, at the transcriptomic and epigenomic levels, are expected to impact downstream functions, using protein-protein interaction network construction and functional enrichment analysis (**Figure 3; Extended Data Table 3**).

**Figure 3.**
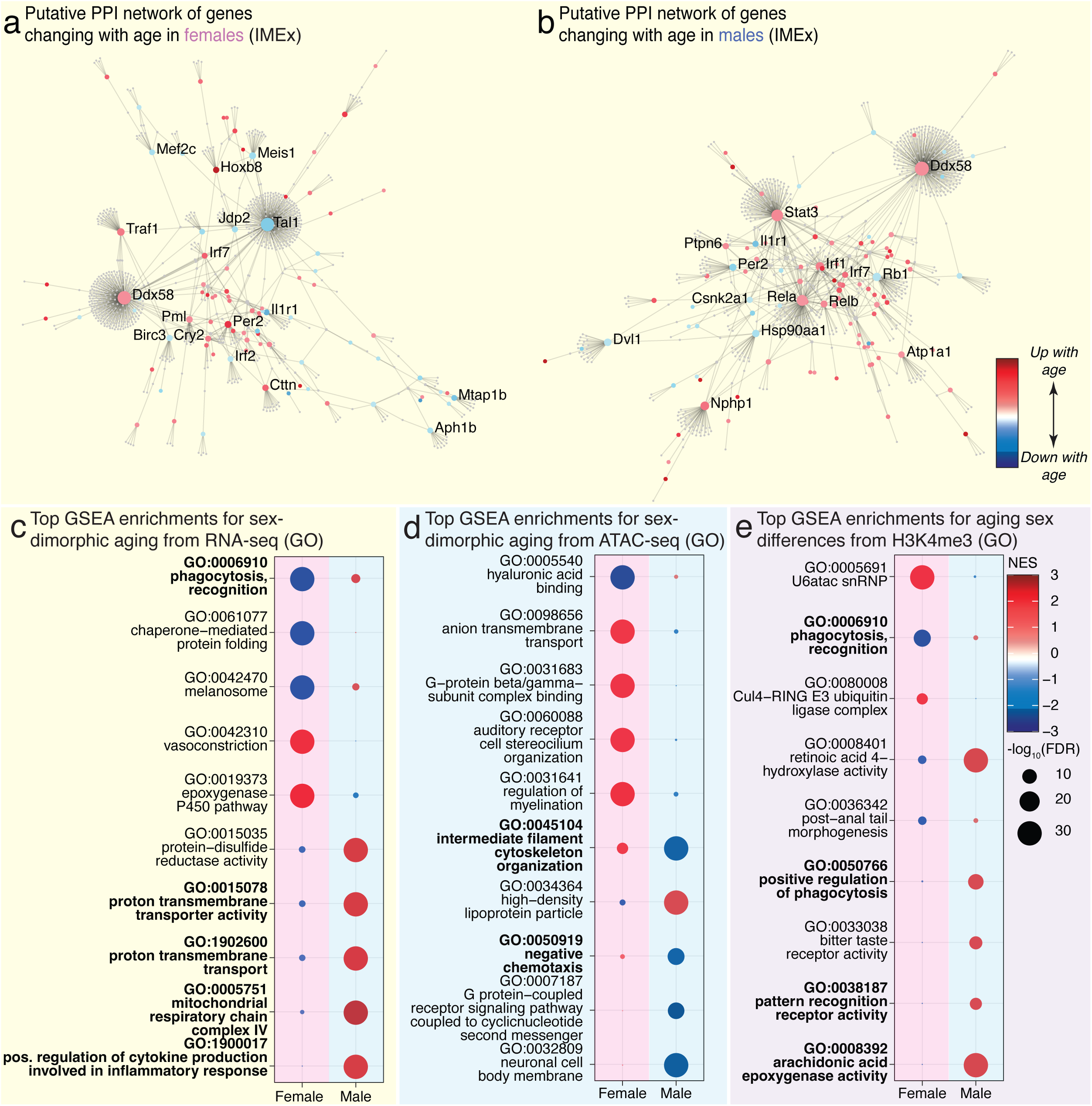
Peritoneal macrophages show sex-dimorphic functional enrichments at the transcriptomic and epigenomic levels. (**a**, **b**) Putative PPI network of genes with significantly changed expression (DESeq2 FDR <5%) with age in female peritoneal macrophages (**a**) or male (**b**) peritoneal macrophages, using NetworkAnalyst and IMEx interaction information. (**c**, **d**, **e**) Top sex-dimorphic GSEA enrichments for Gene Ontology (GO) terms with aging in each sex for RNA-seq (**c**), ATAC-seq (**d**), and H3K4me3 CUT&RUN (**e**). NES: Normalized Enrichment Score. FDR: False Discovery Rate.

First, we generated predicted networks for female and male age-related changes using NetworkAnalyst 3.0^63^ and protein-protein interaction [PPI] information from IMEx/InnateDB data^64^, a knowledge base geared towards innate immunity (**Figure 3a-b**). Our analysis revealed that genes regulated with aging in female and male macrophages are part of clear, interconnected PPI networks (**Figure 3a, b**). Interestingly, although some predicted nodes were identified in both sexes (Irf7, Il1r1), there was a majority of sex-specific nodes (*e.g*. Irf2 and Mef2c in females; Irf1 and Rela in males; **Figure 3a, b**). The tight gene interconnectivity of each predicted PPI network supports the notion that macrophage transcriptional changes with aging are coordinated within each sex, likely leading to differential aging-related functional remodeling.

To gain deeper insights into the downstream functional effects of sex-specific ‘omic’ remodeling, we performed GSEA analysis using gene sets from Gene Ontology (GO; all domains) and Reactome to evaluate differentially regulated processes in aging peritoneal macrophages at the levels of transcription, chromatin accessibility and H3K4me3 signal, both in a sex-independent (**Extended Data Figure 7a-f; Extended Data Table 3A-F**) and sex-dimorphic manner (**Figure 3c-e, Extended Data Figure 7g-i; Extended Data Table 3A-F**). The top gene sets that were regulated in a sex-independent manner were consistent with the expected immune activation with aging^36^ (*e.g*. Gene ontology “innate immune response”; Reactome “complement pathway” and “neutrophil degranulation”; **Extended Data Figure 7a-f; Extended Data Table 3A-F**). These observations are consistent with widespread observations of age-related immune remodeling or “inflamm-aging”^36^, regardless of sex. In addition, we observed strong sex-dimorphic regulation of GO and Reactome genesets in aging peritoneal macrophages (**Figure 3c-e, Extended Data Figure 7g-i; Extended Data Table 3A-F**). Most notably, top significant sex-dimorphic aging gene sets revealed several phagocytosis-related gene sets (*e.g*. GO “phagocytosis, recognition”, “endosomes”; Reactome “cross-presentation of soluble exogenous antigens endosomes”), metabolism-related gene sets (*e.g*. GO “proton transmembrane transport”, “mitochondrial respiratory chain complex IV”; Reactome “respiratory electron transport ATP synthesis by chemiosmotic coupling and heat production by uncoupling proteins”) and inflammation-related gene sets (*e.g.* GO “positive regulation of cytokine production involved in inflammatory response”, “arachidonic acid epoxygenase activity”; Reactome “interleukin 2 family signaling”, “defensins”; **Figure 3c-e, Extended Data Figure 7g-i; Extended Data Table 3A-F**).

Together, our functional enrichment analyses suggest that several processes key to macrophage function may be regulated in a sex-specific manner: (i) phagocytosis capacity, which is required for pathogen and/or dead cell clearance^65,66^, (ii) metabolic preference, which can reflects macrophage inflammatory status — with greater reliance on glycolysis being associated to more pro-inflammatory macrophages^67^, and (iii) macrophage polarization trends, whereby distinct pro- and anti-inflammatory programs are activated^68^.

### Peritoneal macrophages undergo female-specific age-related functional decline

Based on our ‘omic’ analyses, we hypothesized that peritoneal macrophages would undergo sex-specific functional remodeling in phagocytosis capacity, metabolic preferences, and inflammatory state with age. Thus, we carefully selected *ex-vivo* assays to evaluate these phenotypes using primary peritoneal macrophages from young and old, female and male C57BL/6JNia and C57BL/6J mice (**Figure 4a**).

**Figure 4.**
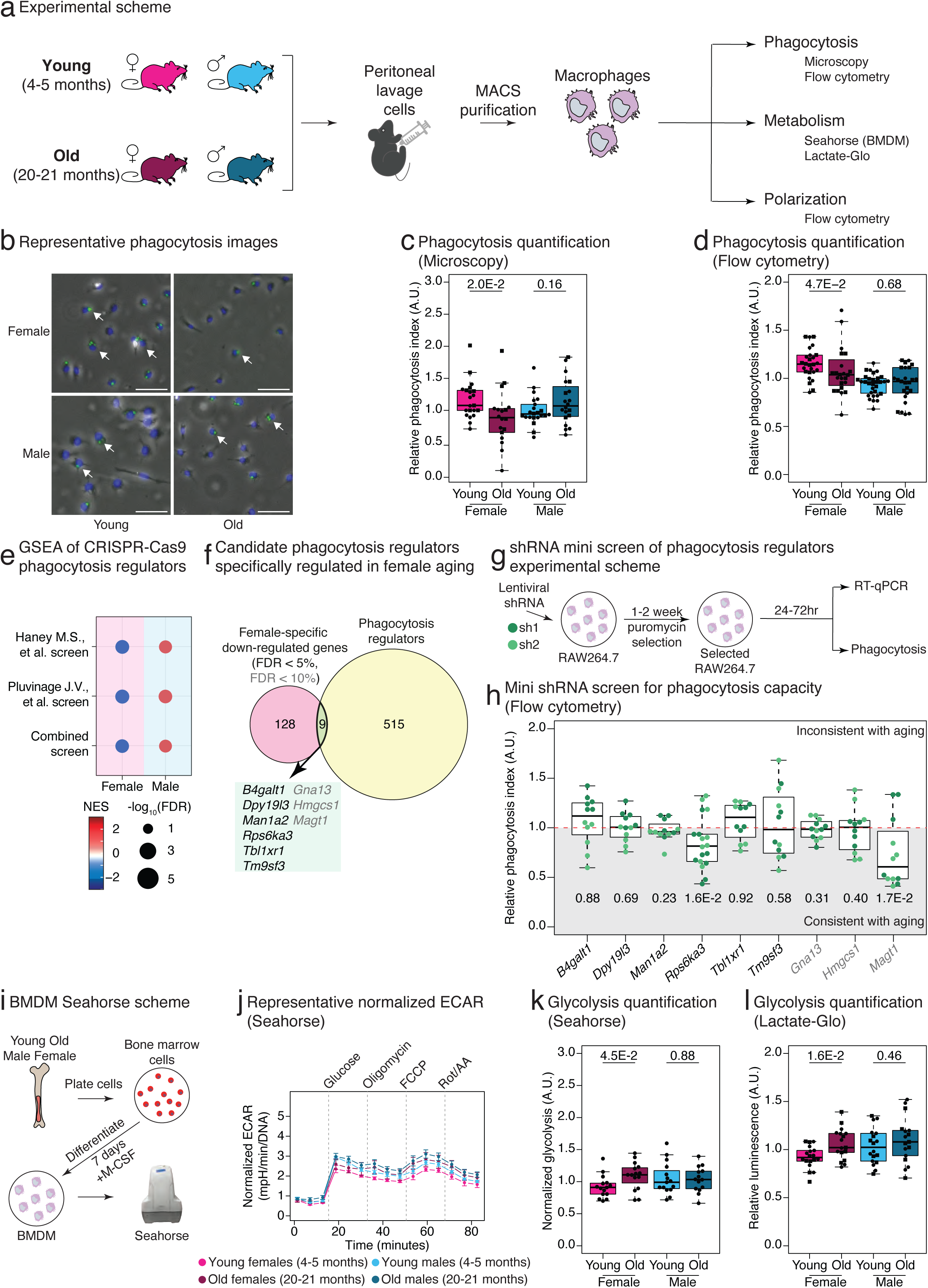
Peritoneal macrophages show female-specific remodeling of phagocytosis capacity and metabolic preferences with aging. (**a**) Experimental scheme for functional profiling of aging murine peritoneal macrophages. (**b**) Representative microscopy image showing phagocytosis events by MACS-purified peritoneal macrophages engulfing Zymosan bioparticles. Blue = DAPI, Green = Zymosan bioparticles, Scale bar: 200μm. (**c**) Quantification of microscopy-based Zymosan phagocytosis assay from young vs. old, female vs. male peritoneal macrophages. The phagocytosis index is calculated by dividing phagocytosing cells by the total number of imaged cells. Due to cohort-to-cohort batch effects, values are normalized to the median value of each independent experiment. Data obtained from animals from 5 independent cohorts (3 from NIA, 2 from JAX; Circles/squares represent NIA/JAX mice, respectively). Young females n = 21, old females n = 18, young males n = 22, and old males n = 20 (variation in group sizes are due to animal deaths prior to experiment). (**d**) Flow cytometry-based Zymosan phagocytosis quantification of peritoneal macrophages. Animals from 6 independent cohorts (3 from NIA, 3 from JAX; Circles/squares represent NIA/JAX mice, respectively). Samples with < 5000 F4/80^+^ events or < 75% F4/80^+^ purity were excluded from the analysis. For (**c**, **d**), significance in non-parametric two-sided Wilcoxon rank-sum. (**e**) GSEA of CRISPR-Cas9 phagocytosis regulators from screens, vs. our foundational aging female and male peritoneal macrophage transcriptome dataset. (**f**) Venn diagram for CRISPR-Cas9 phagocytosis regulators and female-specific down-regulated genes (FDR <10%) according to DESeq2. Genes in the intersection are colored based on DESeq2 significance, with black: FDR < 5% and grey: FDR < 10%. (**g**) Experimental scheme of mini lentiviral shRNA knock-down screen for phagocytosis regulators in RAW264.7 macrophages. (**h**) Flow cytometry Zymosan phagocytosis of RAW264.7 macrophages treated with candidate gene shRNA lentiviruses screen. For visual convenience, the data is normalized to the median of non-targeting and luciferase shRNA hairpins. The horizontal red line in the panel shows the phagocytosis index expected value for normalized data (1; no change upon gene knock-down). Dark green/light green for sh1/sh2 samples. Significance in non-parametric one-sided Wilcoxon rank-sum tests were reported (Hypothesis: consistent regulation with aging). (**i**) Experimental scheme of derivation and metabolic characterization by Agilent Seahorse of aging bone marrow-derived macrophages (BMDMs). (**j**) Representative normalized extracellular acidification rate (ECAR) mpH/min/DNA for BMDMs of one cohort of animals. (**k**) Glycolysis quantification using Agilent Seahorse (n = 15 for young old females and males; animals from 3 independent NIA cohorts). Significance in non-parametric two-sided Wilcoxon rank-sum tests. (**l**) Glycolysis quantification using Lactate-Glo (n = 18 for young males, n = 18 for young females, n = 17 for old males, and n = 17 for old females; animals from 5 independent cohorts, 3 from NIA, 2 from JAX). Significance in non-parametric two-sided Wilcoxon rank-sum tests. For boxplots in panels (**c, d**, **k, l**), circles/squares represent NIA/JAX mice, respectively. For all boxplots (**c, d**, **h, k, l**), the center line of the box plots represents the sample median, the box limits consist of the 25^th^ and 75^th^ percentiles, the whiskers span 1.5x the interquartile range.

To evaluate potential sex-differences in macrophages’ phagocytic ability, we used Alexa Fluor 488-Zymosan bioparticles as cargo to evaluate phagocytosis capacity as a function of age and sex. We evaluated the “phagocytosis index”, defined as the proportion of macrophages internalizing the labelled cargo in a set amount of time, using 2 complementary approaches based on (i) microscopy, after a day of *ex vivo* culture^69^, and (ii) flow cytometry, on freshly isolated macrophages (**Extended Data Figure 8a**; see **Methods**). In line with computational predictions, we observed a significant reduction in phagocytosis capacity specifically in aged female peritoneal macrophages, as demonstrated by both microscopy (p-value = 2.0x10^-2^; **Figure 4b, c**) and flow cytometry (p-value = 4.7x10^-2^; **Figure 4d**). We also observed a consistent suggestive trend of decreased phagocytosis in aging female bone marrow-derived macrophages (BMDMs), a distinct primary macrophage population (p-value = 0.07; **Extended Data Figure 8b**), suggesting that female-specific phagocytic remodeling with aging may not be niche-specific. Because MACS-purified peritoneal macrophages contain a small number of SPMs and aged female JAX (but not NIA) mice exhibit age-related proportional shifts (**Extended Data Figure 4d, e**), we sought to exclude the possibility that compositional properties influenced our data. Accordingly, we reanalyzed phagocytosis using our flow cytometry paradigm to assess the phagocytic capacity of LPMs (F4/80 high) vs. SPMs (F4/80 low) separately (**Extended Data Figure 8c, d**). Notably, the phagocytic capacity of the most abundant tissue-resident LPMs significantly decreased with aging in females (regardless of mouse provider or C57BL/6J substrain; **Extended Data Figure 8d**). Together, these findings indicate that tissue resident LPMs are the primary driver of observed signal, and excluded shifts in macrophage subtypes as a trivial explanation for female-specific age-related remodeling.

In addition, we examined whether differential expression of pattern recognition receptors involved in Zymosan recognition may explain decreased phagocytosis in aging females. Zymosan is recognized by both Tlr2 and Dectin-1 (encoded by *Clec7a*)^70^; there was no evidence of differential regulation of Zymosan receptors *Tlr2* and *Clec7a* in our RNA-seq datasets (**Extended Data Figure 8e**). Surface protein quantification by flow cytometry shows no change in Tlr2 protein expression with aging or sex (**Extended Data Figure 8f, g**). In contrast, we observed decreased surface Dectin-1 signal in aging female macrophages (**Extended Data Figure 8f, g**). However, Dectin-1 levels in aged females matched those of males regardless of age (**Extended Data Figure 8f, g**), despite comparable phagocytosis levels in young animals (**Figure 4c**), suggesting that the observed age-related decrease is not a mere consequence of decreased surface Dectin-1 protein.

We wanted to identify potential drivers of the female-specific age-related decline in phagocytic capacity of peritoneal macrophages. To focus on *bona fide* regulators, we curated a list of phagocytosis regulator genes derived from phagocytosis CRISPR/Cas9 screens^71,72^ (**Extended Data Table 4A**). Consistent with our unbiased GSEA analyses (**Figure 3**), we observed female-specific downregulation of CRISPR/Cas9 phagocytosis regulators (**Figure 4e; Extended Data Table 3G**). We next intersected female-specific aging downregulated genes identified by RNA-seq with CRISPR/Cas9 phagocytosis regulators, revealing nine candidate genes (6 at FDR < 5%: *B4galt1*, *Dpy19l3*, *Man1a2*, *Rps6ka3*, *Tbl1xr1*, and *Tm9sf3;* 3 at FDR < 10%: *Gna13*, *Hmgcs1*, *Magt1*; **Figure 4f; Extended Data Table 4A**). To validate whether candidate regulators could indeed impact Zymosan phagocytosis in murine macrophages, we performed a mini lentiviral short hairpin RNA (shRNA)-mediated knockdown screen in the RAW264.7 immortalized macrophage cell line (**Figure 4g, h**; **Extended Data Figure 9a**). Among our eight candidates, only knockdown of *Rps6ka3* and *Magt1* resulted in a significant reduction in Zymosan phagocytosis capability (**Figure 4h**), consistent with a potential mechanistic role in female-specific age-related decline in phagocytosis.

Next, we evaluated the metabolic preference of primary macrophages with aging and sex. Macrophage function in both homeostasis and disease conditions relies on metabolic reprogramming^73^, whereby increased reliance on glycolysis is characteristic of pro-inflammatory states, and increased preference for mitochondrial respiration is associated with tissue remodeling^67^. We utilized the Agilent Seahorse platform to perform metabolic phenotyping of primary macrophages^74^. Due to required cell yields for this method, we leveraged BMDMs to measure oxygen consumption rate [OCR] and extracellular acidification rate [ECAR], using a protocol designed to evaluate glycolytic and mitochondrial activity^74^ (**Figure 4i-k, Extended Data Figure 10a, b**). Contrary to expectations from ‘omic’ analysis, we did not observe any significant metabolic differences in aging male BMDMs (**Figure 4j, k, Extended Data Figure 10a, b**). However, we observed a significant increase in glycolytic activity specifically in aging female BMDMs (**Figure 4j, k**). To independently validate this phenotype in primary peritoneal macrophages, we used a luminescence-based assay with low cell number requirements, which confirmed a female-specific glycolytic shift with aging (**Figure 4l**).

Finally, we evaluated inflammatory signaling through polarization phenotype with aging and sex. For this purpose, we first curated an unbiased list of genes involved in M1-like and M2-like polarization based on a publicly available BMDM RNA-seq dataset^75^. Interestingly, both female and male macrophages showed significant upregulation of M1-enriched genes, whereas only male macrophages showed concomitant upregulation of M2-enriched genes as well (**Extended Data Figure 11a; Extended Data Table 3I; Extended Data Table 4B**). To test the functional relevance of our ‘omic’ findings of sex-dimorphic activation of macrophage polarization, we leveraged validated flow cytometry quantification of cell surface markers known to be associated to M1-like (CD80, CD86) and M2-like (CD163, CD206) macrophage phenotypes^76–78^ (**Extended Data Figure 11b**). Surprisingly, we found that M1-like cell surface markers CD80 and CD86 increased with age in both sexes (**Extended Data Figure 11c**), indicating a general shift toward a more inflammatory phenotype with aging regardless of sex, consistent with previous studies in the field. In contrast, M2-like markers CD163 and CD206 showed inconsistent age-related changes; notably, CD206—but not CD163—significantly increased with age specifically in female macrophages (**Extended Data Figure 11d**). These observations suggest that shifts in polarization phenotypes are not strongly sex-dimorphic with age.

Together, our functional validation analyses revealed pronounced female-specific age-related functional remodeling of primary peritoneal macrophages particularly in phagocytosis and metabolic programming.

### Impact of ovarian hormone signaling on macrophage phenotypes with female-specific age-related remodeling

Our functional analyses revealed two phenotypes with strong female-specific age-related remodeling: (i) phagocytosis and (ii) metabolic rewiring towards a more glycolytic state. With age-related cessation of reproduction, female mammals undergo drastic age-related decreases in ovarian hormones (*e.g*. estradiol, progesterone)^79^. Interestingly, emerging evidence suggests that ovarian hormones can impact macrophage function, including phagocytosis^80,81^ and metabolism^82^. Consistent with the ability to sense and respond to changes in circulating levels of sex hormones, we confirmed that primary peritoneal macrophages express sex hormone receptors, with *Esr1* (encoding estrogen receptor a) being most robustly expressed (**Extended Data Figure 12a-b**). Hence, we asked whether the known age-related decline in circulating ovarian hormones in female mice^79^, specifically 17b-estradiol [E2] (the predominant form of estrogen in females), may drive female-specific age-related remodeling of peritoneal macrophages.

For this purpose, we analyzed macrophages from young females across the estrus cycle, which experience endogenous, physiological changes in E2; **Extended Data Figure 12c-g**). Using vaginal cytology^83^, actively cycling animals were identified within cohorts of 4 month old C57BL/6NTac female mice to obtain peritoneal macrophages exposed to varying endogenous levels of E2^84^ (proestrus/estrus: high E2, metestrus/diestrus: low E2; **Extended Data Figure 12C; Extended Data Table 1B, C**). Consistent with aging data, peritoneal macrophages isolated from females in low E2 phases of the estrus cycle showed significantly lower phagocytosis *vs*. those in high E2 phases (p-value = 1.3x10^-2^; **Extended Data Figure 12d**). However, metabolic profiling of BMDMs from naturally cycling mice did not reveal E2-dependent differences in metabolic preference (**Extended Data Figure 12e-g**). These experiments suggested that fluctuations of sex-hormones can recapitulate changes in phagocytosis observed with aging.

We next leveraged cohorts of young C57BL/6J female mice subjected to ovariectomy [OVX] vs. sham surgery at 3 months of age, followed by a one-month recovery period prior to euthanasia (**Figure 5a-e; Extended Data Table 1D**). OVX mice experience an immediate and abrupt loss of circulating gonadal hormones, including E2, providing an independent model to evaluate the impact of gonadal hormone level on aging-relevant macrophage phenotypes (**Figure 5a**). In these mice, we then evaluated 3 phenotypes: (i) peritoneal macrophage transcriptomes, (ii) peritoneal macrophage phagocytosis, and (iii) peritoneal macrophage glycolytic activity (**Figure 5a**). The transcriptional landscape of peritoneal macrophages was strongly remodeled in OVX vs. sham animals (**Figure 5b**; samples with > 90% purity; **Extended Data Figure 12h; Extended Data Table 5A**). To determine whether OVX impacts the peritoneal macrophage transcriptome in a manner similar to aging, we assessed how genes differentially regulated upon OVX (FDR < 5%) are modulated in the aging female peritoneal macrophage transcriptome using GSEA (**Figure 5c**). Importantly, genes upregulated/downregulated upon OVX were similarly upregulated/downregulated in aging female macrophages (**Figure 5c**), indicating that OVX recapitulated the transcriptional effects of aging in macrophages. We next evaluated the functional impact of OVX on peritoneal macrophages. Peritoneal macrophages from OVX mice exhibit significantly reduced phagocytic capacity compared to sham-operated mice (p-value = 2.2x10^-3^; **Figure 5d**), similar to aging. However, lactate secretion (and, by proxy, glycolysis) was unaffected by OVX (**Figure 5e**). Together, our findings support the notion that lower E2 levels are sufficient to recapitulate aspects of female macrophage aging (*i.e*. transcriptome, phagocytosis), but not others (*i.e*. metabolic preference).

**Figure 5.**
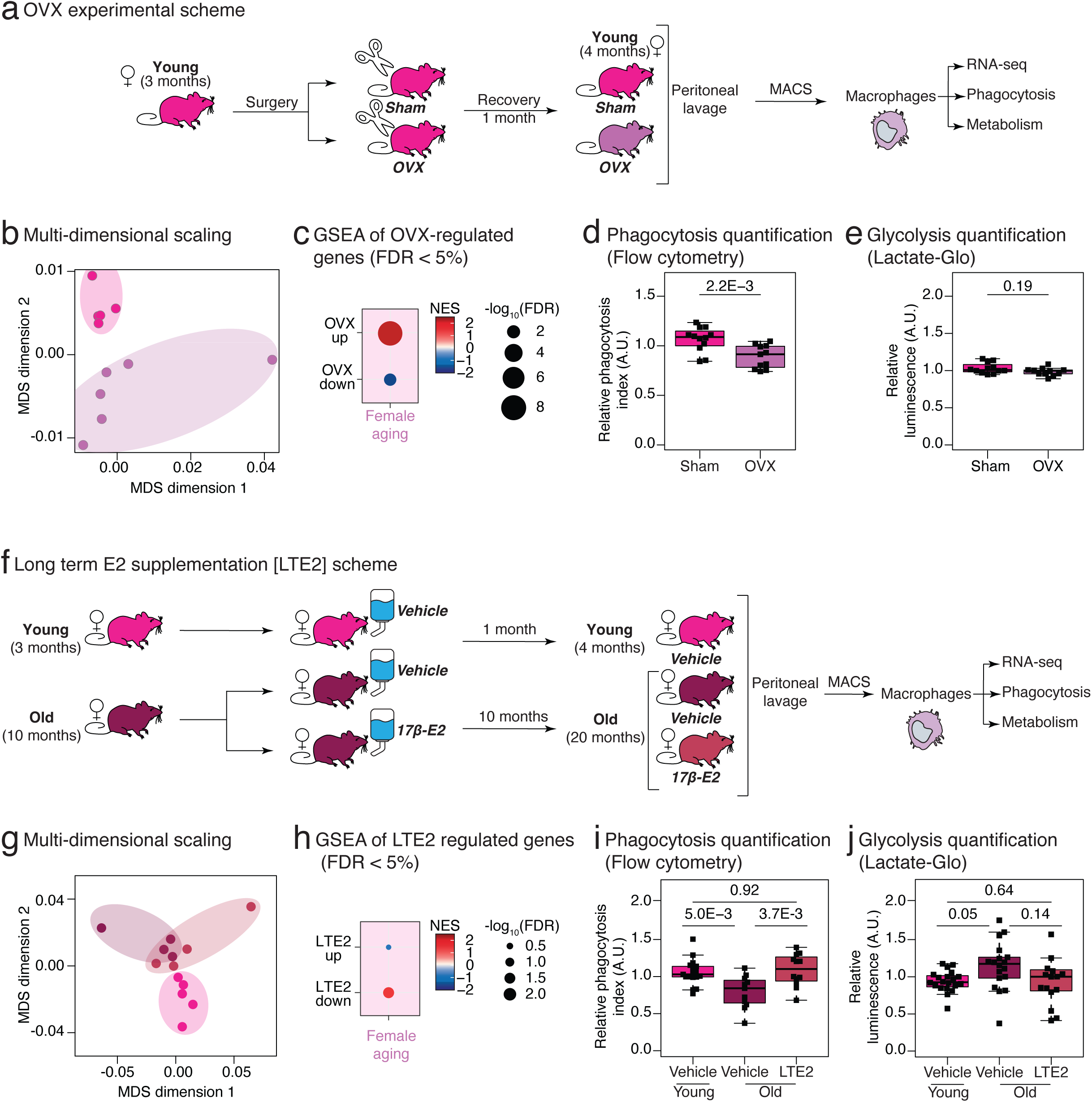
Sustained estrogen signaling is critical to maintain youthful peritoneal macrophage transcriptomes and phagocytic capacity, but not metabolic wiring. (**a**) Experimental scheme outlining the analysis of in female C57BL/6J mice receiving OVX or sham surgery. (**b**) Multi-dimensional scaling (MDS) analysis of gene expression profiles from Sham and OVX mice. (**c**) GSEA of OVX-regulated, *vs*. the foundational aging female peritoneal macrophage transcriptomic dataset. (**d**) Boxplot of phagocytosis quantification of peritoneal macrophages in Sham and OVX mice using flow cytometry (n = 11 for Sham, n = 12 for OVX; animals from 3 independent cohorts). (**e**) Boxplot of glycolysis quantification of peritoneal macrophages in Sham and OVX mice (n = 12 for Sham, n = 11 for OVX; 2 independent cohorts). For (**d**, **e**), significance in non-parametric two-sided Wilcoxon rank-sum. (**f**) Experimental scheme describing late-life long-term 17β-estradiol supplementation (LTE2) in female C57BL/6J mice. (**g**) MDS analysis of gene expression profiles in young vehicle females, old vehicle females, and old LTE2 females; n = 5 for young vehicle females, n = 4 for old vehicle females, n = 4 for LTE2 old females. (**h**) GSEA of LTE2-regulated genes in old female, *vs*. the foundational aging female peritoneal macrophage transcriptomic dataset. (**i**) Boxplot of phagocytosis of peritoneal macrophages in young females, old females, and LTE2-treated old females (n = 17 for young females, n = 11 for old females, n = 12 for E2-treated old females; animals from 4 independent cohorts). Significance in ANOVA/Turkey was reported. (**j**) Boxplot of glycolysis quantification of peritoneal macrophages in young females, old females, and LTE2-treated old females (n = 20 for young females, n = 18 for old females, and n = 16 for E2-treated old females; animals from 4 independent cohorts). Significance in Kruskal-Dunn test is reported. For all boxplots, the center line represents the sample median, the box limits consist of the 25^th^ and 75^th^ percentiles, and the whiskers span 1.5x the interquartile range.

We reasoned that, since decreased E2 conditions phenocopied aspects of female macrophage aging, E2 supplementation in post-estropausal mice may rescue aspects of aging. For this purpose, we first performed short-term late-life E2 supplementation in 19 months old female C57BL/6J mice with E2 (440ng/mL) or vehicle (*i.e*. ethanol 0.1%) in their drinking water^85,86^ for one month, together with young controls exposed to vehicle water (see **Methods**; **Extended Data Figure 13a; Extended Data Table 1E**). We purified peritoneal macrophages as above. Interestingly, the transcriptional landscape of peritoneal macrophages was strongly remodeled in response to aging and short-term late-life E2 supplementation [STE2] (samples with >90% purity; **Extended Data Figure 13 b-c**; **Extended Data Table 5B, C**). The transcriptomes of macrophages from STE2 mice clustered in between those of young vehicle-treated mice and old vehicle-treated mice (**Extended Data Figure 13c**). DESeq2 log_2_(fold change in gene expression) with aging in vehicle-treated animals was significantly anti-correlated with log_2_(fold change in gene expression) in response to STE2 in old animals (Spearman Rho = -0.393; **Extended Data Figure 13d**), and STE2 regulated genes (FDR < 5%) showed opposite regulation to that observed during normal female aging by GSEA (**Extended Data Figure 13e**). Thus, our transcriptomic data is consistent with partial transcriptional rejuvenation in response to STE2. We next evaluated whether STE2 could also functionally reprogram female macrophages toward a more youthful state. While we observed an expected age-related decrease in phagocytosis with vehicle-treated animals (p-value = 3.0x10^−2^), there was no significant rescue upon STE2 (p-value = 0.75; **Extended Data Figure 13f**). Glycolytic output, as measured by lactate production, was increased with aging as expected (p-value = 2.3x10^−2^), but did not show significant changes upon STE2 (**Extended Data Figure 13g**). Therefore, post-estropausal *in vivo* STE2, while showing significant rescue of transcriptional aging markers, did not rescue functional phenotypes.

Post-menopausal hormonal therapy is known to have a limited window of efficacy in humans, with best efficacy if initiated in the perimenopausal interval^87^. Therefore, since C57BL/6 mice undergo estropause between 12-15 months of age^88^, it is likely that short-term post-estropausal exposure cannot fully rescue the effects of age-related E2 loss. We asked whether we could observe stronger rescue of female age-related macrophage phenotypes by initiating E2 supplementation in the peri-estropausal window (10 months of age) and maintaining that supplementation until old age (Long Term E2 supplementation [LTE2]; **Figure 5f; Extended Data Table 1F**). We leveraged the same drinking water supplementation paradigm as for STE2, as it is minimally invasive and inherently well-suited for long-term supplementation (see **Methods**). We phenotyped purified peritoneal macrophages as before, evaluating potential rescue of female aging phenotypes at both the transcriptomic and functional levels. As expected from our STE2 findings, the transcriptional landscape of peritoneal macrophages was strongly remodeled in response to aging and long-term peri-estropausal E2 supplementation [LTE2] (**Figure 5g**; samples with >90% purity; **Extended Data Figure 13h; Extended Data Table 5D, E**). Macrophage transcriptomes from LTE2 mice were situated between those of young vehicle-treated mice and old vehicle-treated mice (**Figure 5g**). DESeq2 log_2_(fold change in gene expression) with aging in vehicle-treated animals was significantly anti-correlated with log_2_(fold change in gene expression) in response to LTE2 in old animals (Spearman Rho = -0.429; **Extended Data Figure 13i**) and LTE2 regulated genes (FDR < 5%) showed opposite regulation to that observed during normal female aging by GSEA (**Figure 5h**). Thus, transcriptomic profiling of LTE2 macrophages is consistent with partial transcriptional rejuvenation, similar to that observed with STE2. Next, we evaluated whether LTE2 could functionally reprogram female macrophages, in contrast with STE2. Indeed, in contrast to what we observed with STE2, phagocytosis capacity was fully restored to the level of control young female macrophages in response LTE2 (p-value = 3.7x10^−3^; **Figure 5i**). In addition, macrophages from LTE2 females showed a non-significant trend of rescued glycolytic output to lower young-like levels (p-value = 0.64; **Figure 5j**). Together, our findings support the notion that sustained E2 levels throughout life are sufficient to blunt aspects of female macrophage aging (*i.e*. transcriptome, phagocytosis), but not others (*i.e*. metabolic preference). Importantly, the fact that LTE2 showed a more comprehensive prevention of aging phenotypes than STE2 suggests that the emergence of female age-related macrophage phenotypes can be prevented by sustained E2 exposure; however, once phenotypic defects are established, they cannot be fully restored to youthful levels.

Finally, since systemic administration of E2 through both STE2 and LTE2 could act on macrophages indirectly through niche cells, we performed *in vitro* E2 supplementation of the immortalized female macrophage cell line J774A.1 (**Extended Data Figure 14a**). We found evidence of profound transcriptional remodeling by MDS (**Extended Data Figure 14b**), which was consistent with “rejuvenation” of the transcriptome in response to E2 exposure according to GSEA (**Extended Data Figure 14c, Extended Data Table 5F**). In addition, *in vitro* E2-supplemented J774A.1 macrophages showed significantly increased phagocytic capacity (**Extended Data Figure 14d**). Thus, the effects of E2 supplementation on macrophage aging phenotypes are likely to be largely cell autonomous.

Our results suggest that age-related decline in circulating E2 in females can explain aspects of female-specific macrophage aging. Since *Esr1* was the highest expressed receptor for E2 in our peritoneal macrophage bulk RNA-seq data (**Extended Data Figure 12a**), we next evaluated potential changes in expression or activity aging peritoneal macrophages. To note, there was a non-significant trend for increased *Esr1* expression in aging female (but not male) macrophages by bulk RNA-seq (p-value = 7.1x10^−2^; **Extended Data Figure 12b**), consistent with the known negative feedback loop of Esr1 regulation^89^. *Esr1* expression levels were too low for robust detection in scRNA-seq (as expected for most genes encoding transcription factors^90^). To determine whether Esr1 may mediate the impact of reduced E2 levels with aging in peritoneal macrophages, we leveraged existing *Esr1* full body knockout mice^91^ (**Extended Data Figure 15a**). We isolated and characterized cells from 3-4 months old *Esr1* knockout and wild-type littermate female mice (**Extended Data Figure 15a; Extended Data Table 1G**). First, we evaluated the transcriptomic impact of constitutive *Esr1* knockout on peritoneal macrophages (samples with > 90% purity; **Extended Data Figure 15b)**. Although *Esr1* knockout led to substantial transcriptional remodeling (**Extended Data Figure 15d**; samples with > 90% purity; **Extended Data Table 5G**), GSEA analysis of *Esr1* knockout-regulated genes surprisingly showed a pattern consistent with transcriptional rejuvenation (**Extended Data Figure 15e**). However, consistent with cycling, OVX and LTE2 functional results, *Esr1* knockout peritoneal macrophages exhibited significantly decreased ability to perform phagocytosis, as determined by flow-cytometry (p-value = 3.7x10^−3^; **Extended Data Figure 15f**), but showed no difference in glycolytic output (p-value = 0.23; **Extended Data Figure 15g**).

Together, our results support that sustained youthful level of E2 exposure, at least in part through Esr1 signaling, are required to maintain youthful transcriptomic signatures and phagocytic capacity, in female peritoneal macrophages across the lifespan, but are not sufficient to preserve low glycolytic dependence.

### Identifying non hormonal mechanisms driving aspects of female-specific macrophage aging

Since age-related changes in E2 signaling can only account for aspects of female-specific age-related peritoneal macrophage remodeling (*i.e*. phagocytosis, transcriptome), we next sought to identify potential molecular mediators contributing to female-specific age-related macrophage remodeling. To do so, we leveraged our comprehensive ‘omic’ datasets (**Figure 1, 2**) as *in silico* screening tools to pinpoint candidate regulators (**Figure 6a**). We prioritized genes encoding transcription factors [TFs], given their broad regulatory influence over the expression of hundreds to thousands of other genes^92^.

**Figure 6.**
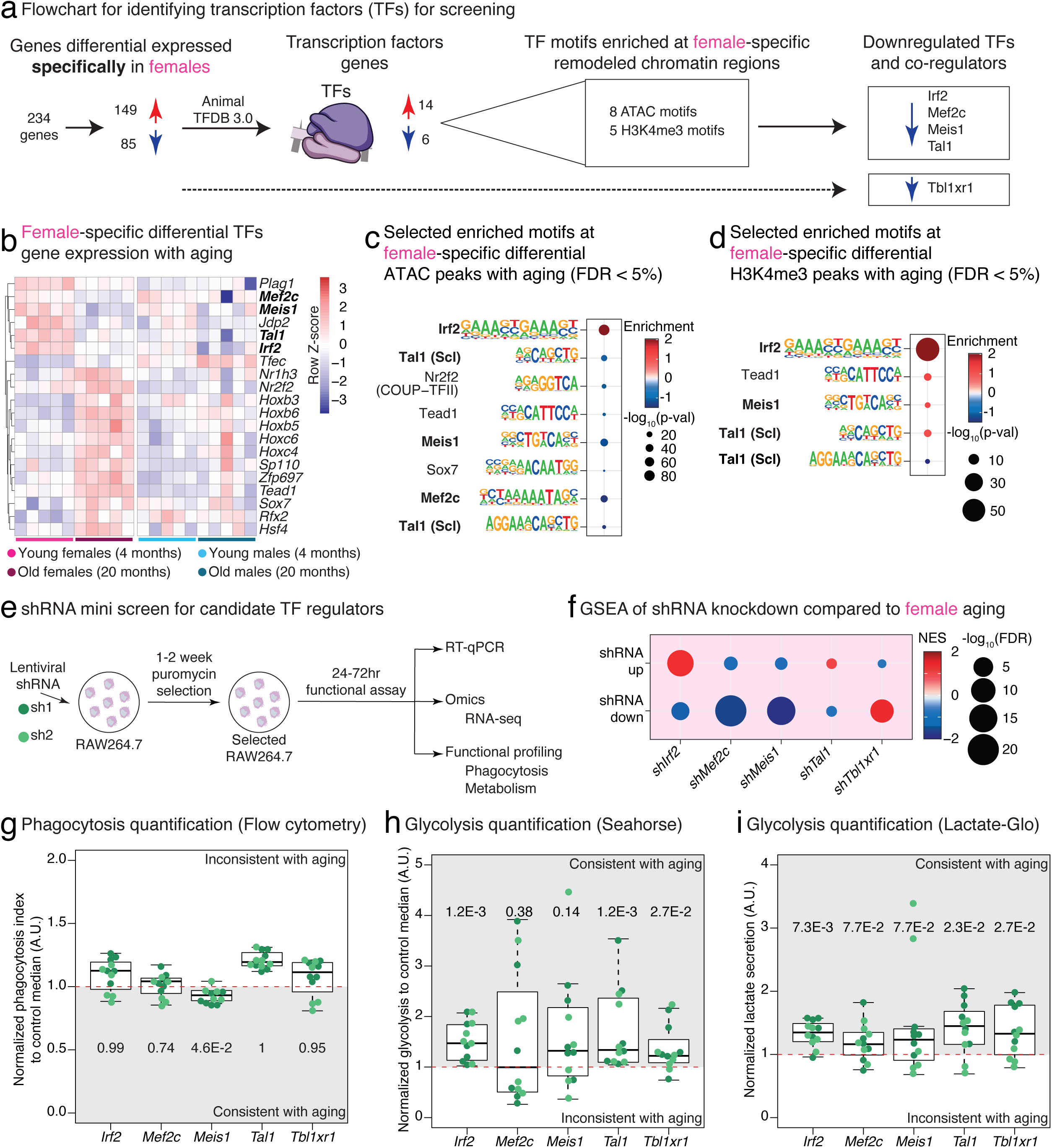
Functional genomics identification of candidate transcription factors driving female-specific aging macrophage ‘omic’ and functional remodeling. (**a**) Flowchart outlining the steps for the selection of top candidate transcription factors (TFs) for screening in RAW264.7 macrophages. (**b**) Heatmap displaying expression of transcription factor (TF) gene expression with female-specific age regulation (FDR < 5% in aging female macrophages, FDR > 10% in aging male macrophages based on DESeq2 analysis). (**c**) Select enriched TF motifs identified at female-specific differentially accessible ATAC-seq peaks with aging (HOMER; FDR < 5%). Only motifs corresponding to TF genes identified in (**b**) are plotted. (**d**) Select enriched motifs identified at female-specific differentially marked H3K4me3 peaks with aging (HOMER; FDR < 5%). Only motifs corresponding to TF genes identified in (**b**) are plotted. (**e**) Experimental scheme of mini shRNA-mediated TF knockdown screen in RAW264.7 macrophages. (**f**) GSEA of TF shRNA knockdown-regulated genes, vs. our foundation aging female peritoneal macrophage transcriptomic dataset. (**g**-**i**) Boxplots showing flow cytometry phagocytosis quantification (**g**), Agilent Seahorse glycolysis quantification (**h**), and Lactate-Glo glycolysis quantification (**i**) in RAW264.7 macrophages transduced with lentiviral TF shRNA against *Irf2*, *Mef2c*, *Meis1*, *Tal1*, and *Tbl1xr1* (n = 6 independent infections for each hairpin). Dark green/light green for sh1/sh2 samples. Significance in non-parametric one-sided Wilcoxon rank-sum tests were reported (Hypothesis: consistent direction as aging). The horizontal red line in the panel shows the expected control value for each assay (1, no effect of gene knockdown). For all boxplots, the center line represents the sample median, the box limits show the 25^th^ and 75^th^ percentiles, and the whiskers span 1.5 times the interquartile range.

First, we leveraged our bulk RNA-seq dataset to identify candidate TF-encoding genes that significantly changed in expression with aging in female, but not male, peritoneal macrophages (**Figure 6a**). We examined evidence for disrupted TF signaling specifically in aging female macrophages using: (i) TF binding motif analysis in ATAC-seq regions remodeled specifically in aging female macrophages and (ii) TF binding motif analysis in H3K4me3 CUT&RUN regions remodeled specifically in aging female macrophages (**Figure 6a**). Mouse TF-encoding genes were obtained from the AnimalTFDB3.0 database^93^ and overlapped with genes with significant age-regulation in females only, revealing 20 candidate TF-encoding genes (**Figure 6a, b**). To note, 14 of these TFs were upregulated with age, and 6 of them were downregulated with age **(Figure 6a, b; Extended Data Figure 16a**). We then performed TF motif analysis in the genomic regions with ATAC-seq and H3K4me3 CUT&RUN signal remodeled specifically in female macrophages with aging using HOMER^94^. Significantly enriched motifs in peaks with female-specific increased or reduced signals were then filtered based on correspondence with our 20 candidate TFs (**Figure 6c**, **d**), which led to the identification of 4 TF candidates with age-related <u>decreased</u> expression (*i.e. Irf2, Mef2c, Meis1, Tal1*), and 3 TF candidates with age-related <u>increased</u> expression (*i.e. Nr2f2, Sox7, Tead1*). For experimental simplicity, we focused on TFs specifically downregulated with aging in female macrophages (*i.e. Irf2, Mef2c, Meis1, Tal1*). Interestingly, these candidate TFs were identified as nodes of the predicted differential PPI network of aging female macrophages (**Figure 3a**), consistent with potentially central regulatory roles. Intriguingly, independent TF footprint analysis using chromVAR^95^ showed a strong increase in IRF2 footprint accessibility in ATAC-seq from old female peritoneal macrophages (**Extended Data Figure 16b; Extended Data Table 6A**). In addition, we also leveraged our scRNA-seq datasets to predict differential TF activity across sex and ages in macrophage single cell transcriptomes using SCENIC^96^, which revealed significant predicted decrease in Irf2 activity and significant predicted increase in Mef2c activity in aging female macrophages (**Extended Data Figure 16c, d; Extended Data Table 6B**), providing another layer of evidence that age-related disruption of these TFs may impact female macrophages. Additionally, *Tbl1xr1*, a candidate phagocytosis regulator (**Figure 4g-h**) that functions as a transcriptional co-regulator^97^, was also identified by SCENIC as showing significant age-related predicted activity decrease (**Extended Data Figure 16c, d ; Extended Data Table 6B**). In analyses thereafter, we considered a set of 4 TFs and 1 TF co-regulator (*i.e. Irf2, Mef2c, Meis1, Tal1, Tbl1xr1*; hereafter referred to as “TFs”) as candidate regulators of female macrophage aging (**Figure 6a**). The impact of candidate TFs on macrophage phenotypes (*i.e*. transcriptome, phagocytic capacity, metabolic preference) was evaluated using RAW264.7 macrophages using a mini lentiviral shRNA knockdown screen (**Figure 6e**; **Extended Data Figure 17a, b;** see **Methods**).

We first analyzed the impact of decreased TF expression on macrophage transcriptomes by mRNA-seq (**Extended Data Figure 17c**), which revealed that TF knockdown led to clear remodeling of macrophage transcriptomes (**Extended Data Figure 17d-e; Extended Data Table 6C-H**). Robust/consistent transcriptional targets of candidate TFs were then used as GSEA gene sets to evaluate whether transcriptional changes elicited by decreased TF expression were consistent with female macrophage aging (**Figure 6f**). Interestingly, knockdown of *Irf2* and *Tal1* produced transcriptional patterns concordant with female macrophage aging, with upregulated genes further upregulated (and downregulated genes further decreased) in aging female macrophages (**Figure 6f**). To note, *Irf2* targets showed substantially stronger collinearity than those of *Tal1* (**Figure 6f**). In contrast, knockdown of *Mef2c* and *Meis1* resulted in downregulation of genes that are strongly downregulated in aging female macrophages, whereas upregulated genes showed weak downregulation opposite to aging (**Figure 6f**). *Tbl1xr1* knockdown led to transcriptional changes in macrophages fully inconsistent with aging (**Figure 6f**). Finally, to assess whether candidate TFs may form a regulatory network, we found that *Mef2c* knockdown significantly reduces *Irf2* expression (**Extended Data Figure 17f**), suggesting that concomitant downregulation of *Irf2* and *Mef2c in vivo* may act synergistically to shape the female aging peritoneal macrophage transcriptome.

Lastly, we evaluated whether decreased expression of candidate TFs recapitulates female-specific age-related macrophage functional remodeling in phagocytosis and metabolic preferences (**Figure 6g-i**). We observed no significant recapitulation of age-associated decline in phagocytosis following TF knockdown, with the notable exception of *Meis1* (**Figure 6g**). Interestingly, *Rps6ka3*, the top hit from our mini-shRNA knockdown screen for candidate regulators of phagocytosis, is significantly downregulated after *Meis1* knockdown (FDR = 0.004; **Extended Data Figure 17g; Extended Data Table 6E**), supporting the notion that *Meis1* downregulation in aged female macrophages occurs upstream of *Rps6ka3* downregulation to promote reduced phagocytosis. In contrast to phagocytosis, both the Agilent Seahorse and Promega Lactate-Glo assays revealed a widespread impact of TF candidate knockdown on glycolytic output (**Figure 6h, i**). Specifically, knockdown of *Irf2*, *Tal1*, and *Tb11xr1* led to significantly increased macrophage glycolytic activity across both assays, consistent with the female aging macrophage phenotype (**Figure 6h, i**).

Thus, our mini-shRNA screen revealed that *Irf2* downregulation led to the strongest (i) transcriptional and (ii) metabolic rewiring consistent with female macrophage aging. Consequently, *Irf2* age-related downregulation represents the best candidate mechanism to explain observations of female-specific glycolytic shift in aging macrophages, which could not be explained by age-related estrogen deprivation (**Figure 5**).

### The Irf2 target network is derepressed in female peritoneal macrophages with aging

Due to its status as our top candidate TF, we next set out to validate regulation of Irf2 expression with aging in primary peritoneal macrophages at the RNA and protein levels (**Figure 7a-c**; **Extended Data Figure 18a-c**). First, RT-qPCR analysis of *Irf2* RNA expression in peritoneal macrophages from independent cohorts of aging C57BL/6JNia and C57BL/6J mice confirmed significant transcriptional downregulation of *Irf2* with aging (**Extended Data Figure 18a**), albeit in both sexes. Importantly, using intracellular immunolabeling and quantification by flow cytometry, we observed a significant decrease in Irf2 protein levels in peritoneal macrophages, specifically in aging female macrophages (**Figure 7c; Extended Data Figure 18b-c**), consistent with our previous analyses. Thus, Irf2 is robustly downregulated in aging female peritoneal macrophages at both the transcriptional and protein levels.

**Figure 7.**
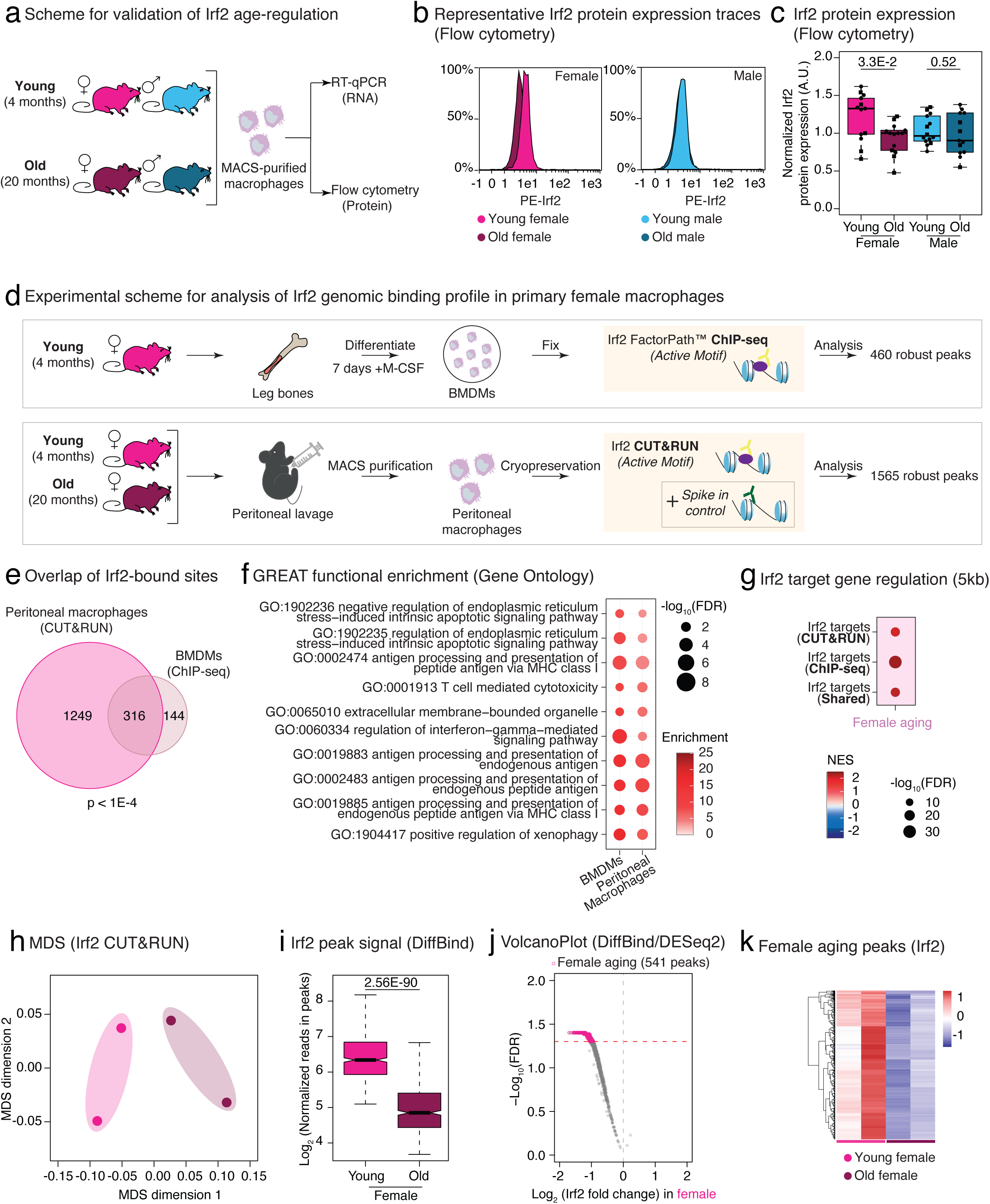
Age-related decrease in Irf2 expression impacts regulation of its target regulatory network. (**a**) Experimental scheme for analysis of *Irf2*/Irf2 expression levels in aging mouse peritoneal macrophages. (**b**) Representative histogram of female vs. male, young vs. old Irf2 protein expression in mouse peritoneal macrophages using intracellular immunolabelling and flow cytometry quantification. (**c**) Boxplot of Irf2 protein quantification by intracellular flow cytometry in peritoneal macrophages from female vs. male, young vs. old mice (young female n = 13, old female n = 14, young male n = 14, and old male n = 13; animals from 3 independent cohorts). Significance in non-parametric two-sided Wilcoxon rank-sum tests were reported (**d**) Experimental scheme for Irf2 genomic binding profiling by ChIP-seq in young female BMDMs and by CUT&RUN in young vs. old female peritoneal macrophages. (**e**) Venn diagram for overlapping robust consensus Irf2 binding sites identified in BMDMs vs. peritoneal macrophages. Significance of overlap from Fisher’s exact test (see **Methods**). (**f**) Top 10 common GREAT enriched terms for Irf2-bound genomic regions in BMDMs and peritoneal macrophages. (**g**) GSEA of direct Irf2 target genes (with a peak within 5kb of TSS) in BMDM ChIP-seq, peritoneal macrophage CUT&RUN or both, vs. our foundational aging female peritoneal macrophage transcriptomic dataset. NES: normalized enrichment score. FDR: false discovery rate. (**h**) Multidimensional scaling (MDS) plot of Irf2 binding profiles in aging female peritoneal macrophages by CUT&RUN, based on DiffBind/DEseq2 spike-in normalized signal. (**i**) Boxplot of DiffBind normalized Irf2 signal over robust Irf2 CUT&RUN peaks (minus those overlapping the mm10 blacklist), revealing overall decreased Irf2 binding on the chromatin of aged female peritoneal macrophages. Significance from DiffBind. (**j**) Volcano plot for differential Irf2 CUT&RUN binding in aged female peritoneal macrophages. 541 peaks were identified as significantly regulated (DiffBind/DEseq2FDR <5%). Note that the overwhelming majority of the signal is on the left-hand side of the plot, consistent with global loss of signal. (**k**) Heatmap of normalized Irf2 CUT&RUN signal in peaks with significant age-regulation. For all boxplots, the center line represents the sample median, the box limits show the 25^th^ and 75^th^ percentiles, and the whiskers span 1.5 times the interquartile range.

To evaluate the possibility that changes in Irf2 expression in aging female macrophages occur independently of sex hormone signaling, we next mined our RNA-seq datasets to extract Irf2 expression information (**Extended Data Figure 18d-e**). Interestingly, *Irf2* transcriptional levels in bulk RNA-seq datasets were not significantly affected in response to OVX (DESeq2 FDR = 0.69), late-life STE2 (DESeq2 FDR = 1), peri-estropausal LTE2 (DESeq2 FDR = 0.57), *Esr1* knock-out (DESeq2 FDR = 0.91), or *in vitro* E2 supplementation of J774A.1 macrophages (DESeq2 FDR = 0.88; **Extended Data Figure 14b**). Thus, our RNA-seq dataset data is consistent with the notion that *Irf2* downregulation occurs independently of age-related changes in E2 signaling.

To independently validate results from our shRNA screen in male RAW264.7 macrophages (**Figure 6**), we also performed *Irf2* shRNA-mediated knockdown in the female J774A.1 macrophage cell line (**Extended Data Figure 19a-d**). As before, we saw widespread transcriptional remodeling using MDS (**Extended Data Figure 19e**). Importantly, transcriptional remodeling following *Irf2* shRNA-mediated knockdown was consistent across both macrophage lines (**Extended Data Figure 19f**).

To better understand its regulatory network in female mouse macrophages, we evaluated the Irf2 chromatin binding landscape in primary macrophages from female C57BL/6JNia mice, using both BMDMs and MACS-purified peritoneal macrophages (**Figure 7d**). Specifically, we decided to use a robust approach to evaluate Irf2 genome-wide binding profiles in primary female mouse macrophages using both ChIP-seq (requiring ∼10 million cells per replicate; performed in abundant BMDM population from young female animals) and CUT&RUN (requiring ∼0.5 million cells per replicate; performed in peritoneal macrophages from young and old female animals). To better understand what impact decreased Irf2 expression may have with aging, we first analyzed the target network of Irf2 in female macrophages, regardless of age. We identified 460 ChIP-seq and 1565 CUT&RUN robust peaks across samples for each assay (**Figure 7d**; **Extended Data Table 7A, B**; see **Methods**), which showed significant overlap across macrophage niches (**Figure 7e**). Importantly, both peak sets showed expected significant enrichment for DNA binding motifs recognized by interferon regulatory factors (IRFs; **Extended Data Figure 20a; Extended Data Table 7C, D**). Irf2-bound regions were spread throughout the mouse genome (**Extended Data Figure 20b**) and mostly annotated to proximal regulatory regions/promoters (≤ 5kb from annotated transcription start sites [TSSs]) and distal intergenic regions (likely enhancers; **Extended Data Figure 20c**). To better define regulatory network of Irf2 in primary macrophages, we next leveraged the Genomic Regions Enrichment of Annotations Tool (GREAT)^98^, which revealed enrichments of GO terms related to immune response (*e.g*, GO:0060334 “regulation of interferon-gamma-mediated signaling pathway” or GO:0019883 “antigen processing and presentation of endogenous antigen”; **Figure 7f**; **Extended Data Table 7E, F**). Enrichments for similar gene sets and terms related to immune cell function regulation were observed when using the gene-centric ChIPseeker tool^99^ (**Extended Data Table 7G, H**). Importantly, GSEA revealed that Irf2 <u>direct</u> target genes (with ChIP-seq and/or CUT&RUN peaks within 5kb of their annotated TSSs) were significantly upregulated in aging female peritoneal macrophages (**Figure 7g**) and in response to Irf2 shRNA mediated knock-down in macrophage cell lines (**Extended Data Figure 20d**), consistent with Irf2’s primary role as a transcriptional repressor^100,101^. Even with a less sensitive overlap-based approach, the majority direct peritoneal CUT&RUN Irf2 targets overlapping with genes significantly regulated in aging female macrophages (FDR < 5%) were upregulated (19 out of 24 genes; **Extended Data Figure 20e**). Interestingly, at least 3 of the upregulated genes in this overlap, *Tm6sf1*, *Usp18*, and *Ifit3*, have been previously reported to be linked to cell metabolism^102–104^. Interestingly, 3 of the downregulated genes corresponded to TF genes (*Irf2, Jdp2, Mef2c*; **Figure 6b**), suggesting the existence of a complex dialog between such TF genes modulated in aging female macrophages.

Next, to evaluate the impact of aging on Irf2 genomic distribution, we focused on our CUT&RUN data, which we generated from peritoneal macrophages from young and old female mice. Since protein levels of Irf2 decreased with age in female peritoneal macrophages (**Figure 7b-c**), spike-in control nuclei from *D. melanogaster* were included in the CUT&RUN reaction at a 1:100 stoichiometry with mouse nuclei to enable proper sample-to-sample normalization despite potential global changes in binding^105^. Importantly, MDS analysis of Irf2 CUT&RUN samples revealed a dramatic separation of samples as a function of age, consistent with substantial remodeling of the Irf2 genomic binding profile with aging (**Figure 7h**). As expected from the change in overall protein levels, normalized Irf2 CUT&RUN signal was significantly reduced in robust peaks based on DiffBind analysis (**Figure 7i**), and all 541 peaks with significant age-related remodeling in Irf2-binding showed decreased binding with aging (**Figure 7j, k; Extended Data Table 7I**). Thus, our CUT&RUN experiment results show that decreased Irf2 expression in peritoneal macrophages with aging leads to significant unloading from target chromatin sites. Importantly, genes with CUT&RUN peaks that showed significant decrease in Irf2 signal in aging female macrophages (FDR < 5%) were significantly upregulated in aging female macrophages at the transcriptional level (**Extended Data Figure 20f**). Together, our analyses support the notion that decreased Irf2 expression with aging in female peritoneal macrophages leads to its decreased presence at target chromatin sites, ultimately promoting the reactivation of its repressed target gene network.

### Age-related downregulation of Irf2 expression may promote female macrophage metabolic reprogramming by lifting *Hk3* transcriptional repression

To gain further insights into how Irf2 downregulation impacts aging female macrophage phenotypes, we focused on peaks that (i) were robustly detected in both our BMDM ChIP-seq and peritoneal macrophage CUT&RUN datasets, and (ii) were within 5kb of annotated TSSs, and thus more easily linked to potential target genes (**Figure 8a**). This led to the identification of 38 robust Irf2 binding regions and associated target genes with potential loss of transcriptional repression during aging (**Figure 8a**). Importantly, we noted that one such robust/shared peak lay within the promoter of the *Hk3* gene (**Figure 8a, b**), and that region was among those with significantly reduced Irf2 CUT&RUN signal in aged female peritoneal macrophages (**Figure 8b; Extended Data Figure 21a**; **Extended Data Table 7I**). Intriguingly, *Hk3* encodes an isoform of hexokinase – the enzyme that catalyzes the rate-limiting step of glycolysis^106^, thus potentially providing a mechanistic explanation linking age-related Irf2 downregulation to increased glycolysis.

**Figure 8.**
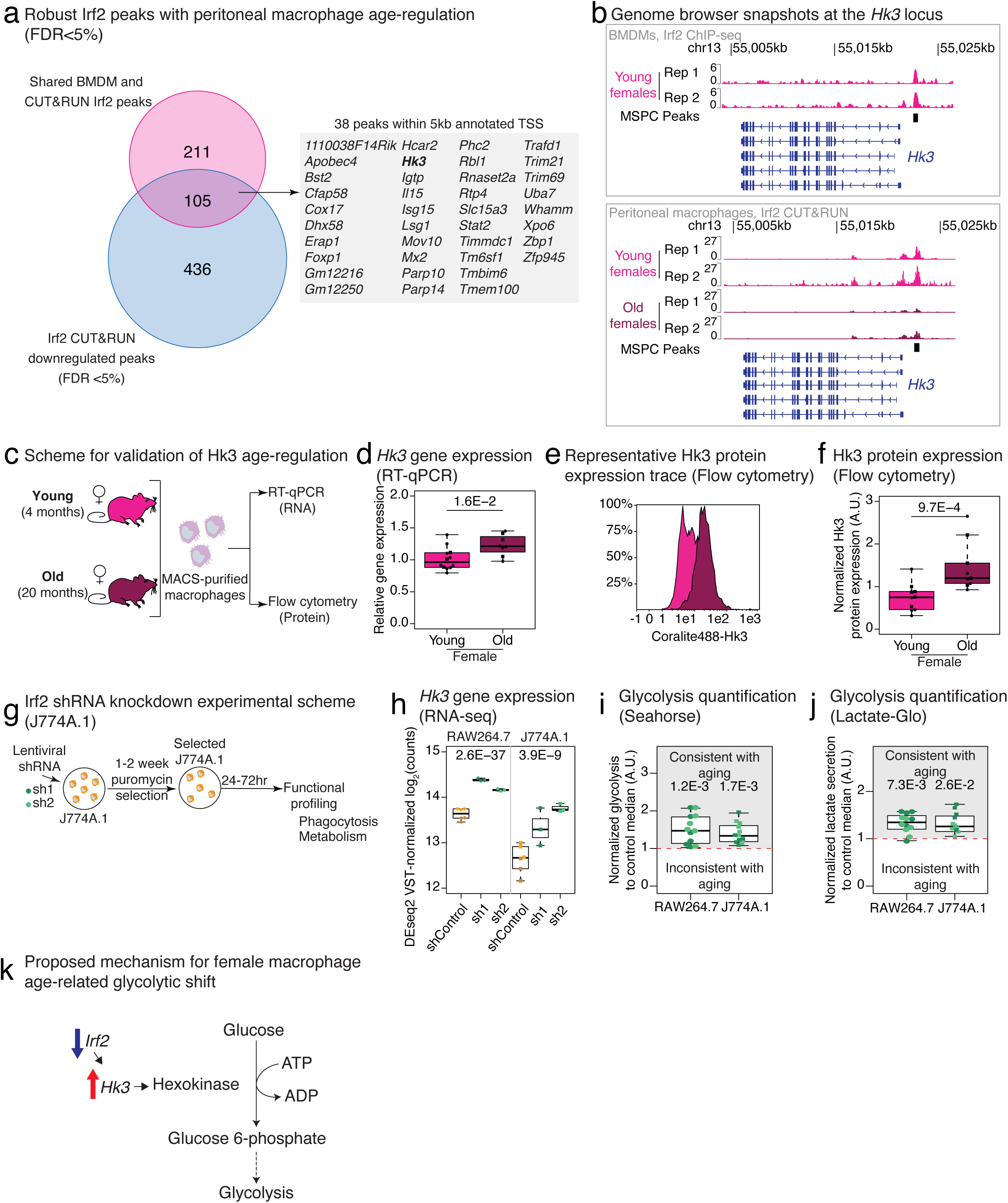
Decreased Irf2 expression and chromatin binding in mouse aging female peritoneal macrophages leads to Hk3 upregulation and increased glycolytic output. (**a**) Overlap analysis of robust Irf2-bound sites in female mouse primary macrophages vs. sites with significant age-related changes in Irf2 occupancy by Irf2 CUT&RUN in aging female peritoneal macrophages. Genes whose TSS is within 5kb of overlapping peaks are enumerated. (**b**) Integrated Genomics Viewer (IGV) snapshots of Irf2 BMDM ChIP-seq (top) or peritoneal macrophage CUT&RUN (bottom) signal at the *Hk3* gene locus on chromosome 13. (**c**) Experimental scheme for validation of age-related changes in *Hk3*/Hk3 expression in aging female peritoneal macrophages. (**d**) Boxplot of *Hk3* gene expression in mouse female peritoneal macrophages using RT-qPCR (n = 12 young, n = 8 old female; animals from 3 independent cohorts; circles/squares represent NIA/JAX mice, respectively). Significance in non-parametric two-sided Wilcoxon rank-sum tests. (**e**) Representative histogram of female Hk3 protein expression between young and old peritoneal macrophages based on intracellular immunolabelling and flow cytometry quantification. (**f**) Boxplot of Hk3 protein quantification by intracellular flow cytometry in peritoneal macrophages between young and old female mice (young female n = 10, and old female n = 9, animals from 3 independent cohorts; circles/squares represent NIA/JAX mice, respectively). Significance in non-parametric two-sided Wilcoxon rank-sum tests. (**g**) Experimental design for lentiviral knockdown of *Irf2* in J774A.1 cell. (**h**) Boxplot of *Hk3* expression in RAW264.7 and J774A.1 macrophage cells upon *Irf2* shRNA knockdown from RNA-seq. Data as DESeq2 VST-normalized log_2_ counts. Circles/squares for RAW264.7/J774A.1 samples; shLuciferase (goldenrod), non-targeting control (light goldenrod), sh1 (dark green), and sh2 (light green); n = 3 infections per hairpin. Significance as DESeq2 FDR. (**i**) Boxplot of glycolysis quantification using Seahorse assays on lentiviral knockdown of *Irf2* RAW264.7 and J774A.1 cell line (n = 6 infections per hairpin). Note that the RAW264.7 data is repeated from Figure 6h for visual convenience. (**j**) Boxplot of glycolysis quantification using Lactate-Glo on lentiviral knockdown *Irf2* in RAW264.7 and J774A.1 cells (n = 6 infections per hairpin). Note that the RAW264.7 data is repeated from Figure 6i for visual convenience. For (**i**, **j**), significance in non-parametric one-sided Wilcoxon rank-sum tests were reported (Hypothesis: same regulation direction as aging). The horizontal red line in the panel shows the expected median control value (1, no change from shRNA-mediated knock-down). (**k**) Proposed mechanism for Irf2-driven age-related glycolytic shift in aging female peritoneal macrophages. For all boxplots, the center line represents the sample median, the box limits show the 25^th^ and 75^th^ percentiles, and the whiskers span 1.5 times the interquartile range.

Thus, we asked how Hk3 expression was regulated in female aging peritoneal macrophages at the RNA and protein levels (**Figure 8c**). Although a trend towards increased *Hk3* expression is shown in our RNA-seq dataset, it did not reach statistical significance, likely due to a high outlier value in the young group (**Extended Data Figure 21b**). Thus, to independently determine whether *Hk3* transcription was regulated in peritoneal macrophages of aging females, we performed RT-qPCR analysis on peritoneal macrophage RNA from three additional independent cohorts. Consistent with predictions based on reduced Irf2 binding at the *Hk3* promoter, RT-qPCR revealed a significant increase in *Hk3* RNA levels in aged female peritoneal macrophages (p-value = 1.6x10^-2^; **Figure 8d**). Notably, no similar age-related trends in expression were observed for either of the other two hexokinase genes - *Hk1* and *Hk2* - in our RNA-seq dataset (**Extended Data Figure 21c, d**). Given that RNA and protein levels can become decorrelated with aging, we next assessed whether changes in *Hk3* transcriptional levels were accompanied by corresponding changes in protein expression. Indeed, quantification using intracellular immunolabelling by flow cytometry revealed significantly increased Hk3 protein levels in aged female peritoneal macrophages (p-value = 9.7 x10^-4^; **Figure 8e, f; Extended Data Figure21e, f**). Critically, *Irf2* knockdown also led to robust *Hk3* transcriptional upregulation in macrophage lines according to mRNA-seq (RAW264.7: FDR = 2.6x10^-37^; J774A.1: FDR = 3.9x10^-9^; **Figure 8h**), as expected from the status of *Hk3* as a robust Irf2 direct target. To note, the *Hk3* response to decreased Irf2 expression was specific, as neither *Hk1* nor *Hk2* was upregulated in response to *Irf2* knockdown (**Extended Data Figure 21g, h**), consistent with the fact that their regulatory regions do not seem to be bound by Irf2 in primary macrophages.

*Hk3* upregulation upon *Irf2* knockdown was accompanied by significantly increased glycolytic output according to the Seahorse and the Lactate-Glo assays in both our RAW264.7 screen (p-value = 1.2x10^-3^, 7.3x10^-3^ respectively; **Figure 6h-i**) and J774A.1 macrophages (p-value = 1.7x10^-3^, 2.6x10^-2^ respectively; **Figure 8i-j**). Consistent with RAW264.7 screen results, *Irf2* knockdown did not impact phagocytic capacity in J774A.1 macrophages (p-value = 0.51; **Extended Data Figure 19g**). Together, our data suggest that decreased *Irf2* expression with aging in female peritoneal macrophages leads to an upregulation of *Hk3*, which then leads to an aberrant age-related increase in glycolytic output (**Figure 8k**).

## Discussion

### A multi-omic resource to study murine tissue-resident macrophages as a function of age and biological sex

To understand how sex may influence aging of the immune system, specifically macrophages, we generated a multi-omic resource focusing on primary resident peritoneal macrophages from young and old, female and male C57BL/6J[Nia] mice, with high-quality transcriptomic (single-cell and bulk RNA-seq) and epigenomic (chromatin accessibility by ATAC-seq and H3K4me3 genomic distribution by CUT&RUN) maps. To our knowledge, our dataset is the largest multi-omic dataset for the study of primary macrophages during aging, and we believe it will be a rich resource to the scientific community in understanding the influence of biological sex on age-related immune remodeling. Analysis of these datasets revealed that, not only is there a baseline difference in genomic regulation of primary macrophages at a young age, but also that aging differentially impacts ‘omic’ profiles in female *vs*. male animals. Although there were shared features of molecular aging in female and male macrophages, we observed much larger sex-dimorphic age-related changes in gene expression and chromatin organization (**Figure 2**).

Notably, genes and genomic regions showing age-related sex-dimorphic regulation were enriched for processes key to macrophage function, including phagocytosis regulation, metabolic wiring, and inflammatory phenotypes (**Figure 3**). Although aging phenotypes of macrophages across various niches have been reported (*e.g*. visceral adipose macrophages, alveolar macrophages, Kupffer cells, *etc*.), previous studies largely focused on using animals of one sex, or mostly reported results combining sexes (reviewed in^36^). A notable exception includes a series of recent studies on aging murine microglia, the brain resident macrophage, which also reported large sex-differences in molecular landscapes with aging^39,107^.

### Peritoneal macrophages show female-specific age-related remodeling in phagocytosis and metabolic wiring

To determine whether sex-differential regulation of molecular landscapes with aging yielded functional consequences, we assessed phagocytic capacity, metabolic preference, and polarization surface markers in isolated peritoneal macrophages. Intriguingly, this revealed two significant female-specific age-related remodeling in primary peritoneal macrophages: (i) a decrease in phagocytic capacity, and (ii) a metabolic shift to increased glycolytic output. Different populations of tissue-resident macrophages have previously been observed to show age-related decreases in phagocytosis across a range of cargo types, but with no insights into the potential influence of sex on these phenotypes^36^. Our data suggest that age-related remodeling in macrophage functions is sex-dimorphic.

To understand the molecular mechanism driving female-specific regulation of phagocytosis, we tested candidate regulator genes of phagocytosis from CRISPR/Cas9 screens that were specifically downregulated in aged female macrophages through a mini shRNA knockdown screen (**Figure 4h**). Our results identified two phagocytosis regulators (*Magt1* and *Rps6ka3*) whose knockdown mimicked the age-related decrease in phagocytosis of female macrophages, consistent with the notion that their downregulation with aging may explain mechanistically our observed female-specific age-related decrease in macrophage phagocytic capacity. Although its impact during aging has not been evaluated before, *Magt1* is a Magnesium transporter located in the endoplasmic reticulum involved in protein glycosylation with a critical function in immune cells^108,109^, which was identified as required for proper phagocytosis of Zymosan in a whole genome CRISPR/Cas9 screen of human phagocytes^71^. Interestingly, *Rps6ka3* encodes a member of the S6 kinase family, whose activity is regulated by mTOR^110^, which is a key regulator of aging^111^, autophagy^112^, phagosome maturation^113^, and macrophage polarization^114^. *Rps6ka3* was also shown to play a direct role in innate immune responses^115^ and was found to be required for phagocytosis of Zymosan in human phagocytes^71^. Thus, downregulation of *Magt1* and *Rps6ka3* in aging female peritoneal macrophages is likely to mechanistically underlie the female-specific age-related phagocytosis defect that we observed.

Interestingly, despite ‘omic’-based predictions of male specific metabolic rewiring, we identified a metabolic shift specifically in aging female macrophages (both peritoneal macrophages and BMDMs), leading to increased glycolytic output. Increased glycolytic output is known to support inflammatory phenotypes and ROS-production in macrophages and is a characteristic of disease-associated macrophages^116^. Furthermore, a recent study of aging murine microglia also identified a female-specific increase in glycolytic output, which was driven by increased C3a – C3aR signaling^39^. Thus, our observations are consistent with a potential shift to a more dysfunctional state in female macrophages with aging.

### Sustained estrogen signaling is necessary for youthful macrophage phagocytosis

Since the age-related functional remodeling we uncovered were female specific, we reasoned that age-related decreases in circulating ovarian hormones may explain these phenotypes. Indeed, previous work found that E2 signaling can shape macrophage phenotypes^80,81^. Importantly, decreased exposure to youthful E2 levels by ovariectomy recapitulated aspects of female-specific macrophage aging, including transcriptional remodeling and decreased phagocytosis, but had no impact on glycolytic output of female macrophages. Similar functional outcomes were observed in macrophages from mice lacking estrogen receptor gene *Esr1*, the most abundantly expressed estrogen receptor in peritoneal macrophages, suggesting the functional impact of youthful E2 may be mediated through ERa.

Both late-life STE2 and peri-estropausal LTE2 led to partial rejuvenation of the peritoneal macrophage transcriptional landscape. In contrast, only long-term LTE2, initiated around the onset of estropause (9-12 months of age in C57BL/6 mice)^117^ led to rescue of the female-specific age-related phagocytosis decrease. Neither STE2 or LTE2 significantly rescue aged female macrophage metabolic defects. Our mouse observations are consistent with accumulating data showing that delayed onset of hormone therapy in post-menopausal women leads to decreased efficacy (and even to an increased risk of adverse side effects)^118–120^. Thus, our results suggest that youthful E2 levels are necessary (and sufficient if maintained throughout aging) to maintain (i) aspects of the youthful female macrophage transcriptome, and (ii) youthful phagocytosis capacity in peritoneal macrophages. However, neither decreased E2 signaling (*i.e*. OVX, low E2 estrus cycle phases, *Esr1* KO) nor sustained youthful E2 exposure significantly influenced glycolytic output, suggesting that the female-specific metabolic shift of peritoneal macrophages occurs independent of circulating sex hormones levels.

### TF-encoding genes are involved in female-specific age-related macrophage phenotypes

We also identified a set of putative TF encoding genes that were specifically downregulated in aging female macrophages and may shape female-specific aging phenotypic remodeling: *Irf2*, *Mef2c*, *Meis1*, *Tal1*, and *Tbl1xr1* (**Figure 6**). Interestingly, with the exception of *Tbl1xr1*, knockdown of candidate TF genes recapitulated phenotypic aspects of aging female macrophages, with *Irf2* and *Tal1* knockdown most closely resembling the effect of aging on macrophage transcriptomes (**Figure 6g**). Among candidates, only knockdown of *Meis1* led to aged-like decreased phagocytic capacity, likely due to downregulation of phagocytosis regulator *Rps6ka3*. To note, *Meis1* expression was unaffected in macrophages from OVX mice or in response to late-life short term E2 supplementation (**Extended Data Table 5A, C**), suggesting that its impact on phagocytosis may occur independently of E2 signaling. In contrast, knockdown of *Irf2*, *Tal1*, and *Tbl1xr1* promoted increased glycolytic output, suggesting that their age-related expression downregulation may underlie the female-specific glycolytic shift we observed in female peritoneal macrophages. How age-related decreased expression of these TFs in female macrophages may synergize together (and with decreased E2 signaling) to shape female-specific aging phenotypes is an interesting avenue for future studies.

### Female-specific age-related downregulation of Irf2 leads to increased *Hk3* expression and glycolytic output

Since *Irf2* knockdown best mimicked the aging female macrophage transcriptome and led to the strongest increase in glycolytic output, we focused on elucidating how it may drive aspects of female-specific peritoneal macrophage phenotypes (**Figure 7**). Interestingly, Irf2 shares a similar DNA-binding domain as Irf1, but works as a competitive antagonist of Irf1-mediated gene expression^121^. Consistent with decreased Irf2 activity driving aspects of aging female macrophage molecular landscapes, ChIP-seq analysis revealed that *bona fide* Irf2 target genes in primary macrophages were significantly upregulated and overlapped with remodeled chromatin regions in aging female macrophages. Most notably, ChIP-seq identified a strong region of Irf2 binding in the promoter of *Hk3* (the gene encoding Hexokinase 3). Both female macrophage aging and *Irf2* knockdown led to significant upregulation of *Hk3*, suggesting that decreased *Irf2* expression with aging is responsible for increased *Hk3* expression. It is noteworthy that, hexokinases catalyze the rate-limiting first step of glycolysis (the phosphorylation of glucose on C6)^122^, thus gatekeeping cellular glycolytic potential. Thus, our data suggest that the female-specific increase in glycolytic output during peritoneal macrophage aging is driven by downregulation of *Irf2* and loss of repression for downstream target *Hk3*. Interestingly, this mechanism that we identified in peritoneal macrophages is distinct from the one previously reported in microglia^39^, suggesting that metabolic rewiring of female macrophages is driven by niche-specific mechanisms.

Together, our study provides the first systematic characterization of sex-differences in immune aging in the context of macrophages and provides mechanistic insights into how sex-specific aging phenotypes can be achieved. Intriguingly, our data suggests that both mechanisms involving age-related changes in gonadal hormone levels and mechanisms independent of gonadal hormones can drive sex-dimorphism in the aging trajectories of immune cells. Ultimately, such mechanisms could be leveraged as potential targets to rejuvenate immune responses in a sex-specific manner, the first and easy step towards personalized geroscience.

## Material and Methods

### Mouse husbandry

All animals were treated and housed in accordance with the Guide for Care and Use of Laboratory Animals. All experimental procedures were approved by the University of Southern California’s Institutional Animal Care and Use Committee (IACUC) in accordance with institutional and national guidelines. Experiments described in this manuscript were performed on approved IACUC protocols 20770, 20804, 21004 and 21454. All animals were euthanized between 8-11 am using a “snaking order” to minimize batch-processing confounding factors related to circadian effects and post-mortem interval, by using by CO_2_ asphyxiation followed by cervical dislocation.

For intact aging analyses, male and female C57BL/6JNia mice were obtained from the National Institute on Aging (NIA) colony at Charles Rivers, or male and female C57BL/6J mice were obtained from the Jackson Laboratories (3 and 19 months, acclimated at USC for 4-6 weeks) **(Extended Data Table 1A).**

For estrus cycle tracking experiments male and female C57BL/6NTac were obtained from Taconic Biosciences (3 months, acclimated at USC for 2 weeks; **Extended Data Table 1B**). The cycle stage of adult female mice was characterized by vaginal lavage daily for 3 weeks until euthanasia on 3-4 months old mice. Mice that seen at least a full cycle or seen of the different phases during the three weeks of vaginal smear were taken for euthanasia (**Extended Data Table 1C**). To reduce animal waste, non-cycling mice were housed with a mixture of clean and used male bedding to elicit additional cycle to stimulate cycling. Mice were cycled for another two weeks and mice that were cycling were taken for euthanasia.

For ovariectomy (OVX) *vs*. sham surgery experiments, female C57BL/6J mice were obtained from Jackson Laboratories (8 weeks old animals, acclimated at USC for 4 weeks) (**Extended Data Table 1D**). C57BL6/J female mice gone through sham or OVX surgery at 12 weeks of age. All surgery procedures and post-operative care were conducted by University of Southern California Department of Animal Resources veterinarians and veterinarian technicians. Surgery entry point was from the back to preserve the peritoneal cavity as much as possible for post-mortem peritoneal cell collection. Mice went through a recovery period of 4 to 5 weeks before euthanasia.

For late-life short-term E2 supplementation experiments, female C57BL/6J mice were obtained from Jackson Laboratories at age 10 and 80 weeks and acclimated at USC for 2 weeks before treatment initiation (**Extended Data Table 1E**). For supplementation, 17b-estradiol (E2) was used in the purest chemical grade that could be obtained (³98% purity; Sigma-Aldrich #E8875). We generated our E2 stock solution of 5 ng/mL in pure 100% ethanol (Sigma-Aldrich #E7023). E2 stock solution and 100% ethanol were filtered sterilized on 0.22mm PES filters (VWR #76479-024) and stored at -20°C for no longer than 1 month. Drinking water of C57BL6/J mice was supplemented with either vehicle (0.1% EtOH) or 440 ng/mL of E2 for 4 weeks, corresponding to ∼94µg/kg of body weight/day^85,86^. Treatment water was changed every 2-3 days (3 times a week) to maximize treatment potency and limit potential E2 degradation. Water levels and mice were monitored daily. Water supplementation of E2 in this manner was found to be a safe and effective alternative to daily injections^85,86,123^. Mice were euthanized ∼4 weeks after treatment initiation for experimental phenotyping. For peri-estropausal long-term E2 supplementation experiments, female C57BL/6J mice were obtained from Jackson Laboratories at age 8 and 36 weeks and acclimated at USC for ∼4 weeks before treatment initiation (**Extended Data Table 1F**). E2 was prepared and stored in the same manner as for the late-life short-term supplementation paradigm. Long-term E2 supplementation was initiated in 10 months old animals with either vehicle (0.1% EtOH) or 440 ng/mL 17β-estradiol (E2) until they reached 20 months of age. Young controls received vehicle water for 4 weeks prior to experimental phenotyping, so that they would reach 4 months of age when old animals of the same cohort reached 20 months of age. Cohorts of animals were euthanized as sets for experimental phenotyping.

For B6N(Cg)-*Esr1^tm4.2Ksk^*/J [*Esr1* KO]^91^ (JAX #026176) experiments, matched female knockout [KO] and wild-type [WT] littermates were obtained from Jackson Laboratories (**Extended Data Table 1G**). After receiving mice at USC, *Esr1* KO/WT mice were acclimated and aged at USC until they reached 12-16 weeks before euthanasia.

Charles Rivers, Jackson Laboratories, and Taconic Biosciences have specific-pathogen-free (SPF) facilities. All animals were acclimated at the SPF animal facility at USC for >2 weeks before any processing or intervention to minimize the impact of travel stress. Some animals had to be euthanized prior to experimental endpoints upon recommendation of USC veterinarians due to welfare concerns and could thus not be included in the study.

### Cell hashing, single-cell RNA-seq library preparation & sequencing of peritoneal cells

Single-cell libraries were prepared either using Chromium Single Cell 3’ Library & Gel Bead Kit v2 (10X Genomics #PN-120237) [NIA pooled cohorts 1 & 2], or the Chromium Next GEM Single Cell 3L GEM, Library & Gel Bead Kit v3.1 (10X Genomics #PN-1000121) [JAX and NIA single animal cohorts with HTO labeling] according to manufacturer’s instructions (10x Genomics User Guide Chromium Next GEM Single-cell 3′ Reagent Kits v3.1 (CG000204, Rev D)). For the two experiments using the v2 system, isolated peritoneal cells were pooled from each biological group with equal cell number to ensure equal representation of all animals, based on cell concentration estimates from a COUNTESS cell counter (Thermo Fisher Scientific # AMQAF1000). This resulted in 4 libraries each (*i.e*. young female, young male, old female, old male) for each of 2 independent cohorts. In this case, cell suspensions were loaded for a targeted cell recovery of 3,000 cells per sample. For the two experiments using the v3.1 system, cell hashing was performed on isolated peritoneal cells from each individual using Biolegend TotalSeq^TM^-A Antibodies according to the manufacturer’s instructions (Biolegend #155801, #155803, #155805, #155807, #155809, #155811, #155813, #155815, #155817, #155819). Specifically, 0.5 million peritoneal cells were first incubated with mouse FcR-blocking Reagent (Miltenyi Biotec #130-092-575) at 4°C for 10 minutes. Then, each blocked sample was incubated with 1μg of specific TotalSeq Cell Hashing antibody (HTOs 1-10) at 4°C for 30 minutes. After incubation, stained cells were washed 3 times, and assessed for viability and yield using the COUNTESS cell counter (JAX) or the DeNovix CellDrop (Denovix #CellDrop FL-UNLTD) (NIA). Making sure no overlapping HTOs were included in a pool, samples were pooled in equicellular mixes for 3 reactions (JAX) or 2 reactions (NIA), with targeted cell recoveries of 14,000 and 30,000 cells, respectively. Samples were run on Chromium Single Cell A Chips (10X Genomics #PN-120236) [v2 reactions] or Chromium Next GEM Chip G (10X Genomics #PN-2000177) [v3.1 reactions] according to the manufacturer’s instructions. For the v3.1 reactions, HTO Additive primer v2 (Integrated DNA Technologies) was added to the cDNA amplification reaction to amplify the HTO molecules. The amplified HTOs were recovered in the cleanup step for the cDNA amplification, and HTO libraries were built using the 2x NEBNext PCR Master Mix and Truseq indexing oligos (Integrated DNA Technologies). Partially amplified and final single-cell RNA-seq libraries were quantified by a dsDNA High Sensitivity assay (Invitrogen #Q32854) on a Qubit 3.0 Fluorometer (Thermo Fisher Scientific #Q33216). Quality and size distribution of single-cell libraries were assessed on the 4200 TapeStation system (Agilent Technologies #G2991A) with High Sensitivity D1000 DNA ScreenTapes (Agilent Technologies #5067-5582) and D1000 reagent (Agilent Technologies #5067-5583). Libraries were sequenced on an Illumina Novaseq 6000 generating 150 bp paired-end reads at Novogene USA. Raw FASTQ reads have been deposited to the sequence read archive under accession PRJNA1094925.

### Computational processing of 10xGenomics sequencing libraries

#### Mapping, cell filtering and demultiplexing

For all datasets, raw reads were aligned to the mm10 mouse genome and processed using cellranger 6.0.2^124^ to generate gene-barcode matrices. For our hashed v3.1 datasets, HTO quantification was performed using CITE-seq-Count v.1.4.4^125^ with parameters “-cbf 1 -cbl 16 -umif 17 -umil 26” to parse UMI/HTO sequences. Gene-barcode matrices and HTO counts were loaded in R 3.6.3^126^ for downstream processing using Seurat 3.2.2^127,128^. We used SoupX v1.6.1^129^ to correct counts for any interference by ambient RNA in our libraries. Cells that were detected in the cellranger data (NIA v2 library) or in both cellranger and HTO data (NIA v3 and JAX v3 libraries) were filtered for downstream analysis. Dead and low-quality cells were filtered out by selecting for cells that contained between 300-6000 genes and < 25% mitochondrial reads. After removal of low-quality unwanted cells, UMI expression data was normalized for downstream processing using SCTransform(), regressing out the impact of number of genes and mitochondrial read load (“vars.to.regress = c(“nFeature_RNA”, “percent.mito”)”).

#### Singlet identification, doublet removal and cell demultiplexing

Dataset dimensionality was determined using (i) the dimension where the principal components only contribute 5% of standard deviation, (ii) cumulative PCs contribute 90% of the standard deviation, and (iii) the dimension where the percent change in variation between the consecutive PCs is less than 0.1%, using the smallest value. This was determined to be 16 (NIA v2), 21 (NIA v3) and 16 (JAX v3), respectively.

For all our datasets, potential doublets were annotated using Doubletfinder v2.0.3^130^ in each particular library, using the predicted doublet rate based on cell capture provided by 10x Genomics. For the NIA v2 dataset, all cells annotated as singlets by Doubletfinder were retained for downstream analyses.

In addition, for our NIA v3 and JAX v3 libraries, we took advantage of HTO libraries to add a second layer of doublet identification and removal. Specifically, HTO counts were added to the Seurat object as an independent assay and normalized using centered log-ratio (CLR) transformation. Cells were demultiplexed according to their HTO barcode of origin using the MULTIseqDemux() function into their component samples. For these datasets, cells were considered to be singlets if and only if they were annotated as singlets by Doubletfinder and MULTIseqDemux.

#### Dimensional reduction and cell clustering

Singlet UMI expression data was normalized and scaled for downstream analyses using SCTransform(), regressing out the impact of number of genes, mitochondrial read load and predicted cell cycle scores (“vars.to.regress = c(“nFeature_RNA”, “percent.mito”, “S.Score”, “G2M.Score)”). The first N principal components (PCs), where N is the estimated dimensionality of the dataset (see above) were used to compute the UMAP transformation with RunUMAP() and cluster cells using FindNeighbors() and FindClusters().

#### Cell type annotation using SingleR

To identify the most likely cell type annotation for each cell, singlets were annotated using SingleR v.1.0.1^131^, together with the reference ImmGen^132^ and the mouse RNAseq^133^ expression datasets. To guarantee annotation robustness, only cells annotated to the same cell type using both references were kept and annotated with the consensus prediction for all downstream analyses.

#### Seurat object merging

After independent pre-processing for each dataset, the 3 cohort datasets were also merged into a single Seurat object for analysis. Reciprocal PCA was used to integrate data from the 3 datasets and mitigate batch effects, using the top 3000 most variable genes and k = 10 anchors.

#### Cell type proportion analysis

We calculated a per library cell type proportion using annotations and analyzed changes in proportions for the most abundant cell types (B-cells, Macrophages and T-cells) across biological groups (**Figure 1D**). In addition, as an orthogonal approach, we computed changes in cell type proportions from the combined and dataset-specific Seurat objects using ‘scProportionTest’ v0.0.0.9000^45^, which runs a permutation test on two different single-cell groups for each cell type and returns the relative change in cell type abundance between groups with a confidence interval (**Extended Data Figure 1a-d**).

#### Cell type prioritization using AUGUR

We used the R package AUGUR v1.0.3^134^ to determine which peritoneal cell types were most transcriptionally impacted by age and sex, in the combined and separated objects (**Figure 1c, Extended Data Figure 3a-f**).

#### Muscat pseudobulking, differential expression analysis and meta-analysis

Since single-cell level differential analyses suffer from high false discovery rates, we used muscat to perform differential gene expression analyses on our scRNA-seq datasets^135^. Due to clear batching in our design, each dataset (NIA v2, NIA v3 and JAX v3) were processed separately for analysis in R v 3.6.3^136^. We used ‘muscat’ v1.0.1^135^ to derive pseudobulked count tables from our single cell gene expression data by biological sample and cell type. Only cell types with ≥ 5 cells across all samples in each dataset were used for downstream analyses to minimize technical noise and maximize interpretability. To diminish the potential impact of technical noise in each experiment on our result, we used ‘sva’ v3.34.0^137^ on the pseudobulked count tables to estimate spurious noise on gene expression, and limma v3.42.2 was used to regress estimated effects from the counts. We used ‘DESeq2’ v1.26.0^136^ to perform differential gene expression analysis on the cleaned gene counts matrices for each cell type in each dataset, modeling the effect of aging in each sex separately so as to determine how similar aging trajectories were between sexes. DESeq2 results for each cell type, in each sex for each dataset were next used for meta-analysis to determine robustly regulated genes as a function of sex in each cell type.

To identify consistently regulated genes with aging as a function of sex across datasets, we used a meta-analysis approach using ‘metaRNASeq’ v1.0.5^138^. We performed this analysis on B-cells, macrophages and T-cells, as these were the only cell types with robust detection across biological groups in our 3 scRNAseq datasets (**Extended Data Figure 1e**). We focused our analyses on genes detected across datasets for each cell type, and performed p-value combination using the inverse normal method^138^. Genes with (i) a combined adjusted p-value < 0.05 and (ii) with consistent directionality of regulation in the 3 datasets were considered significantly regulated with aging (**Extended Data Table 2A-F**). Next, we compared significantly differentially regulated genes with aging from our meta-analysis in aging female vs. male cells (**Figure 1e**), which showed moderate overlap. The scale of overlapping transcriptional changes with aging was determined using the Jaccard Index (corresponding to the size of the intersection of gene lists, over the size of their union). The Jaccard index is equal to 1 in cases of perfect overlap, and to 0 when there is no overlap^139^. Jaccard index similarity of age-regulated genes between sexes in each robust cell type is reported in **Figure 1d**.

### Peritoneal lavage immune cell proportion

Flow cytometry analysis was used to validate changes in proportions of immune cell types within the peritoneal cavity from at least two independent cohorts of aged C57BL/6JNia mice, aged C57BL/6J, OVX *vs*. sham C57BL/6J mice, E2-supplemented C57BL/6J mice, and *Esr1* KO mice. A panel of antibodies was used to evaluate differences in the proportion of B-cells, macrophages, and T cells (see below). Among live cell singlets, B-cells were defined as CD19^+^, T-cells were defined as CD3^+^, and macrophages were defined as F4/80^+^.

Cells were isolated from the peritoneal cavity using a peritoneal lavage. Then, 200,000 peritoneal cells were washed once by adding 1mL of PBS and 2mM EDTA pH 7.2 (Miltenyi Biotec #130-091-222) supplemented with 0.5% BSA (Miltenyi Biotec #130-091-376; MACS resuspension buffer) in a 5mL polystyrene round bottom tube (Falcon #352054), then centrifuged 300g for 10 minutes at 4°C. Cells were resuspended in MACS resuspension buffer in the presence of FcR-blocking solution (concentration following manufacturer recommended dilution) for 10 minutes at 4°C. Afterwards, cells were stained with antibodies for 10 minutes at 4°C protected from light, and excess antibody was washed away using MACS resuspension buffer. Stained cells were then analyzed by flow cytometry on a MACSQuant Analyzer 10 (Miltenyi Biotec #130-096-343), and flow cytometry data was analyzed using FlowLogic V8. Gating was determined using fluorescence-minus-one [FMO] controls for each color used in the experiment to ensure that positive populations were solely associated with the antibody for that specific marker. Raw cytometry data, including FMO controls, were deposited for naïve aging, sham vs. OVX, E2 supplementation, and *Esr1* KO to Figshare under 10.6084/m9.figshare.28934213, 10.6084/m9.figshare.29169674, 10.6084/m9.figshare.29169737, and 10.6084/m9.figshare.29169707, respectively.

### Peritoneal lavage immune cell yield

To assess the age-related impact to immune cell number within the peritoneal cavity, we took total peritoneal lavage cell counts from the DeNovix CellDrop and the identified the fraction of each immune cell types (B-cell, macrophage, T-cell; **Extended Data Figure 2b**) to calculate original immune cell numbers in the peritoneal cavity. To calculate the number of immune cell types (B-cell, macrophage, and T-cell), the total peritoneal lavage cell counts were multiplied by the fraction of immune cell obtained from flow cytometry (**Extended Data Figure 2c**).

### Isolation of primary peritoneal macrophages

Isolation of primary peritoneal macrophages was performed on aged C57BL/6JNia, aged C57BL/6J, estrus cycle tracking C57BL/6NTac, OVX *vs*. sham C57BL/6J, E2-supplemented C57BL/6J, and *Esr1* KO mice following a protocol from Lu et al. 2020^69^ with modifications to remove excess T-cells and B-cells in old animals. Briefly, peritoneal cells were obtained by using a 25G needle (BD #305125) to inject 10mL of filtered sterilized 3% bovine serum albumin (BSA) (VWR #0332-500G) in DPBS (Corning, 21-031-CV) into the peritoneal cavity of euthanized mice. Using a 20-gauge needle (BD #305175), the BSA solution was then harvested back from the peritoneum, now referred to as “peritoneal lavage”. Peritoneal lavages were centrifuged 300g for 10 minutes at 4°C, and cells were resuspended in 1mL of MACS resuspension buffer. To remove any potential contaminating red blood cells, samples were treated with 10mL of 1x Red Blood Cell [RBC] lysis solution (Biolegend #420301) inverted to mix and incubated at 2 minutes at room temperature, then centrifuged 300g for 10minutes at 4°C. To obtain single cell suspension and removal of cell clumps from RBC lysis, macrophages were filtered on 30µm MACS SmartStrainers (Miltenyi Biotec #130-110-915).

Peritoneal macrophages were then isolated using Miltenyi Biotec peritoneal isolation kit (Miltenti Biotec #130-110-434), with modifications to the magnetic labeling step improve purity of macrophage isolation regardless of age, treatment, and genotype. Indeed, while using the unmodified manufacturer’s protocol, we found that aged mice exhibited poor macrophage purity, with large contamination of B-cells (likely due to B-cell population expansion with aging; **Extended Data Figure 2b**). To improve peritoneal macrophage purity, we routinely added additional biotin-labelled antibodies during the “macrophage (peritoneum) biotin-antibody cocktail” labeling step to help deplete potential excess B-cells, T-cells, and monocytes from peritoneal lavages. 5mL of biotinylated anti-CD19 (Miltenyi Biotec #130-119-757), anti-CD79b (Miltenyi Biotec #130-105-832), anti-CD3e (Miltenyi Biotec #130-093-179), and anti-Ly6c (Miltenyi Biotec #130-111-776) antibodies were added during incubation for 10 minutes at 4°C. After primary antibody incubation, all other steps in isolating peritoneal macrophages were performed according to the manufacturer’s instructions. This allowed us to routinely achieve high purity for isolated peritoneal macrophages. Magnetic Activated Cell Sorting (MACS)-purified peritoneal macrophages were used for downstream experiments, including ‘omic’ and functional profiling.

### Purity and identity of MACS-purified peritoneal macrophages

Potential phenotypic differences in MACS-purified peritoneal macrophage purity were assessed from independent cohorts of aged C57BL/6JNia, aged C57BL/6J, OVX vs. sham C57BL/6J, E2-supplemented C57BL/6J, and *Esr1* KO mice. Briefly, 100,000 MACS-purified peritoneal macrophages were used for flow cytometry labeling. After FcR-blocking, macrophages were stained using F4/80-APC (1:50; Miltenyi Biotec #130-116-547) and CD11b-Vioblue (1:50; Miltenyi Biotec #130-113-238) antibodies according to the manufacturer’s instructions. Stained cells were analyzed by flow cytometry on the Miltenyi Biotec MACSQuant Analyzer 10. Flow cytometry results were processed using the FlowLogic V8 software. Pure macrophages were designated as double positive F4/80^+^ CD11b^+^ cells. From these, large peritoneal macrophages were defined as F4/80^+high^, and small peritoneal macrophages were defined as F4/80^+low^. The raw flow cytometry data for naïve aging, sham vs. OVX, E2 supplementation, and *Esr1* KO analyses has been deposited to Figshare under 10.6084/m9.figshare.28934198, 10.6084/m9.figshare.29169578, 10.6084/m9.figshare.29169581, and 10.6084/m9.figshare.29169587, respectively.

### RNA purification from primary macrophages and macrophage cell lines

For RNA isolation of primary peritoneal macrophages, frozen cell pellets were resuspended in 500mL of TRIzol reagent (Thermo-Fisher #15596018), and total RNA was purified following the manufacturer’s instructions. RNA quality was assessed using the Agilent Technologies 4200 TapeStation system using High Sensitivity RNA ScreenTapes (Agilent Technologies #5067-5579) together with RNA ScreenTape Sample Buffer (Agilent Technologies #5067-5577).

For RNA isolation of modified RAW264.7 and J774A.1 cells (see below), 60-80% confluency in a 6-well plate were lysed using 1mL of TRIzol reagent, and RNA was extracted using DirectZol columns (Zymo #R2052) following manufacturer’s protocol.

### RNA-seq library preparations for primary peritoneal macrophages and macrophage cell lines

For primary peritoneal macrophages, we constructed ribosomal RNA depleted strand-specific bulk RNA-sequencing libraries in house as previously described Lu et al. 2021^24^. Briefly, 450ng of total RNA was subjected to ribosomal-RNA depletion using the NEBNext rRNA Depletion Kit v2 (New England Biolabs #E705X), according to the manufacturer’s protocol. Strand specific RNA-seq libraries were then constructed using the SMARTer Stranded RNA-seq Kit (Clontech #634841, #634863), according to the manufacturer’s protocol. Since this version of the kit became discontinued, the LTE2 primary peritoneal macrophage libraries were constructed using its successor SMART-seq Total RNA Mid Input (Clontech #635049), according to manufacturer’s protocol. Libraries were quality controlled on the Agilent Technologies 4200 TapeStation system with Agilent Technologies D1000 ScreenTape and Agilent Technologies D1000 reagent, before multiplexing the libraries for sequencing. Paired-end 150bp reads were generated on the Illumina Novaseq X or Novaseq 6000 platforms at Novogene USA. Raw FASTQ sequencing reads were deposited to SRA under Bioprojects accession PRJNA1095555 and PRJNA1101936.

For macrophage cell line derived samples, total RNA samples were sent to the Novogene corporation for unstranded mRNA-sequencing. Briefly, the company performed mRNA selection using poly (A) capture, RNA fragmentation, cDNA reverse transcription and then unstranded RNA sequencing library preparation. RNA libraries were sequenced as paired-end 150bp reads on the Illumina Novaseq X or Novaseq 6000 platforms. Corresponding raw FASTQ sequencing reads were deposited to SRA under Bioprojects accession PRJNA1096052, PRJNA1222371 and PRJNA1313760.

### Bulk RNA-seq analysis pipeline

Paired end 150bp FASTQ reads were hard-trimmed to 100bp at their 3’ end to improve mapping rates and the first 9bp at the 5’ end were trimmed to eliminate priming biases^140^. This was done using Fastx Trimmer 0.0.13 (https://github.com/agordon/fastx_toolkit), or Trimgalore 0.6.6 (github.com/FelixKrueger/TrimGalore) after a system upgrade made it impossible to use Fastx Trimmer. For Trimgalore, additional required stringency filters were added to eliminate any bases with phred score < 15 (corresponding to < 1% of bases). Trimmed reads were mapped to the mm10 genome reference using STAR 2.7.7a^141^. Read counts were assigned to genes from the UCSC mm10 reference using featureCounts from subread 1.5.3^142^, using either the stranded or unstranded option based on library construction (see above). Read count tables were imported into R v3.6.3^126^ for downstream processing. Based on general RNA-seq processing guidelines, only genes with mapped reads in at least half of the samples in each analysis were considered to be expressed and retained for analysis.

The ‘DESeq2’ R package v1.26.0 was used for normalization and differential gene expression analysis of the RNA-seq data in R^136^ using appropriate analytical covariates (*e.g*. sex, age, treatment, etc.). Genes with a false discovery rate < 5% in any analysis were considered statistically significant. For analysis of sex-specific age-regulation of genes, we considered a gene to be specifically regulated with age in females (males) if the DESeq2 FDR was < 5% in females (males) and > 10% in males (females). DESeq2 results are reported in **Extended Data Table 2G**.

### ATAC-seq library generation for primary peritoneal macrophages

A second independent cohort of C57BL/6JNia mice was used to assess the impact of age and sex on peritoneal macrophage chromatin accessibility (**Extended Data Table 1A**). We constructed ATAC-seq libraries in house as described Lu et al. 2021^24^, using the omni-ATAC protocol and starting from 50,000 MACS-purified peritoneal macrophages^50^. Libraries were quality controlled using the Agilent Technologies 4200 TapeStation system with Agilent Technologies High Sensitivity D1000 DNA ScreenTape and reagent before multiplexing the libraries for sequencing. Paired-end 150bp reads were generated on the Illumina Novaseq 6000 platform at Novogene USA. Raw FASTQ sequencing reads were deposited to SRA under Bioproject accession PRJNA1095555.

### ATAC-seq preprocessing pipeline

Paired-end ATAC-seq reads were adapter-trimmed using NGmerge 0.3^143^, a tool recommended for use in ATAC-seq preprocessing which clips overhanging reads in a sequence-independent fashion. Trimmed paired-end were mapped to the mm10 genome reference using bowtie2 v2.5.1^144^. After alignment, PCR duplicates were removed using the ‘rmdup’ function of samtools 0.1.19^145^. Reads from all samples were pooled define “meta-peaks” for robust differential accessibility analysis using samtools ‘merge’. We used MACS2 v2.2.7.1 to identify ATAC-seq accessible regions^146^. A normalized read count matrix was obtained in R v3.6.3^136^ using our mapped read bam files over MACS2 meta-peaks using R package ‘Diffbind’ 2.14^147^, which was used for downstream analyses. ‘DESeq2’ 1.26.0 was used for normalization and differential gene expression analysis of ATAC-seq data in R^136^ using appropriate analytical covariates (*e.g*. sex, age). Peaks with a false discovery rate < 5% in any analysis were considered statistically significant. For analysis of sex-specific age-regulation of chromatin region accessibility, we considered a chromatin region to be specifically remodeled with age in females (males) if the DESeq2 FDR was < 5% in females (males) and > 10% in males (females). DESeq2 results are reported in **Extended Data Table 2H**.

### H4K4me3 Cleavage Under Targets and Release Using Nuclease [CUT&RUN] library generation for primary peritoneal macrophages

A third independent cohort of C57BL/6JNia mice was used to assess the impact of age and sex on peritoneal macrophage H3K4me3 landscapes (**Extended Data Table 1A**). We used a CUT&RUN protocol adapted from the kit manual from EpiCypher, with modifications to account for the use of immune cells (*i.e*. to avoid unscheduled immune activation from ConA signaling on cells). All buffers were made in-house.

#### Nuclei isolation

250,000 MACS-purified peritoneal macrophages were harvested and resuspended in 100µL of 1X DPBS. Samples were centrifuged for 3 min at 600g at room temperature, and supernatant was discarded. Cells were resuspended in 100µL of ice-cold Nuclear Extraction Buffer (20mM HEPES [Sigma-Aldrich #H3375], pH 7.9, 10mM KCl [Sigma-Aldrich # P9541], 0,1% Triton X-100 [VWR # M143], 20% Glycerol [VWR # BDH1172], 0.5mM Spermidine [EMD Millipore # 56766] and 1x Protease Inhibitor [Roche #11873580001]) and incubated on ice for 10 min. Samples were centrifuged for 3 min at 600g at 4 °C, and the supernatant was removed. Samples were resuspended in 100µL of ice-cold Nuclear Extraction Buffer and kept on ice until further processing.

#### Concanavalin A bead activation

11µL per sample of gently resuspended Concanavalin A (ConA) beads were transferred to fresh 1.5mL tubes. Tubes were placed on a magnetic separation rack until the slurry cleared and the supernatant was removed. 100µL of ice-cold Bead Activation Buffer (20 mM HEPES, pH 7.9, 10 mM KCl, 1 mM CaCl_2_ and 1 mM ManCl_2_) was added to each tube and pipetted 10 times to mix and wash the beads for a total of two washes. Washed beads were resuspended in 11µL of ice-cold Bead Activation Buffer. 10µL of activated bead slurry was transferred into fresh 1.5mL tubes. Beads were kept on ice until further processing.

#### Immunolabeling of chromatin

Isolated nuclei were collected by centrifugation at for 3 min at 600g at 4 °C, and the supernatant was discarded. Nuclei were resuspended in 100µL of Wash Buffer (20mM HEPES, pH 7.5, 150mM NaCl, 0.5mM Spermidine, and 1x Protease Inhibitor) and transferred to the tubes containing the activated Concanavalin A beads. Samples were incubated at room temperature for 10 min. Tubes were placed on a magnetic separation rack until the slurry cleared and supernatant was removed. 50µL of ice-cold Antibody Buffer (20mM HEPES, pH 7.5, 150mM NaCl, 0.5mM Spermidine, 1x Protease Inhibitor, 0.01% Digitonin and 2mM EDTA) was added to each sample. 0.5µL of anti-H3K4me3 antibody (Active Motif #39159) was added to each sample. Samples were incubated on a nutator overnight at 4°C. Tubes were placed on a magnetic separator rack until the slurry cleared and the supernatant was removed. While keeping the tubes on the magnetic separator, 250µL of ice-cold Digitonin Buffer (20mM HEPES, pH 7.5, 150mM NaCl, 0.5mM Spermidine, and 1x Protease Inhibitor and 0.01% Digitonin) was added to the tubes and supernatant was removed without disturbing the tubes. Digitonin Buffer wash step was repeated once more for a total of two washes. 50µL of ice-cold Digitonin Buffer was added.

#### pAG-MNase binding and DNA purification

2.5µL of CUTANA pAG-MNase (EpiCypher # 15-1016) was added to each sample and gently mixed by vortexing. Samples were incubated at room temperature for 10 min and placed on a magnetic separator. Supernatant was removed by pipetting. While the tubes were on the magnet separator, 250µL of ice-cold Digitonin Buffer was added to each sample and removed. Digitonin Buffer wash step was repeated once more for a total of two washes. 50µL of ice-cold Digitonin Buffer was added to the tubes and gently mixed by vortexing. 1µL of 100mM CaCl_2_ was added to each sample and gently mixed by vortexing. Samples were incubated on a nutator for 2 hours at 4°C. 33µL of Stop Buffer (340mM NaCl, 20mM EDTA, 4mM EGTA, 50µg/mL RNaseA, 50µg/mL GlycoBlue Coprecipitant) was added to each sample. Samples were incubated at 37 °C for 10 min in a thermocycler. Samples were placed on a magnetic separator until the slurry cleared and the supernatant was collected to new 1.5 ml tubes. DNA purification was performed using the DNA Clean & Concentration-5 kit (Zymo Research #D4013) according to the manufacturer’s instructions.

#### Sequencing library preparation and sequencing

CUT&RUN sequencing libraries were prepared using the NEBNext Ultra II Library Prep Kit for Illumina kit (NEBNext #E7645), according to the manufacturer’s instructions. H4K4me3 CUT&RUN libraries were multiplexed and sent for sequencing on the Illumina NovaSeq 6000 platform to Novogene USA. De-multiplexed FASTQ files were provided by Novogene for data analysis. Raw FASTQ sequencing reads were deposited to SRA under Bioproject accession PRJNA1095555.

### H4K4me3 CUT&RUN preprocessing pipeline

Paired-end H4K4me3 CUT&RUN reads were adapter-trimmed using NGmerge 0.3^143^. Trimmed paired-end were mapped to the mm10 genome reference using bowtie2 v2.5.1^144^. After alignment, PCR duplicates were removed using the ‘rmdup’ function of samtools 0.1.19^145^. Reads from sequenced samples were pooled in order to define “meta-peaks” for robust differential accessibility analysis using samtools ‘merge’. We used MACS2 v2.2.7.1 to identify H4K4me3 modified regions^146^. A normalized read count matrix was obtained in R v3.6.3^136^ using our mapped read bam files over MACS2 meta-peaks using R package ‘Diffbind’ 2.14^147^, which was used for downstream analyses. ‘DESeq2’ 1.26.0 was used for normalization and differential gene expression analysis of H4K4me3 CUT&RUN data in R^136^ using appropriate analytical covariates (*e.g*. sex, age). Peaks with a FDR < 5% in any analysis were considered statistically significant. For analysis of sex-specific age-regulation of H4K4me3 signal intensity, we considered a chromatin region to be specifically remodeled with age in females (males) if the DESeq2 FDR was < 5% in females (males) and > 10% in males (females). DESeq2 results are reported in **Extended Data Table 2I**.

### Multidimensional Scaling analysis

To perform Multidimensional Scaling (MDS) analysis, we used a simple distance metric between samples based on the Spearman’s rank correlation value (1-Rho), which was then provided to the core R command ‘cmdscale’. Dimensionality reduction was applied to DESeq2-normalized data, obtained as described in relevant sections above.

### Putative Protein-Protein Interaction [PPI] network analysis with NetworkAnalyst

Genes with significant regulation with aging in female or male peritoneal macrophages (DESeq2 FDR < 5%; **Extended Data Table 2G**) were used as input for network analysis with NetworkAnalyst 3.0^63^. Significant age-regulated genes in each sex and their corresponding DESeq2 computed log_2_(fold-change) were used as input. In both cases, we had NetworkAnalyst use PPI information derived from IMEx/InnateDB data, a knowledgebase specifically geared for analyses of innate immune networks^64^, to construct putative PPI networks. In each case, the largest subnetwork determined by NetworkAnalyst was used in our analyses (**Figure 3a, b**).

### Functional enrichment analysis by Gene Set Enrichment Analysis

We used the Gene Set Enrichment Analysis (GSEA) paradigm^148^ through the ‘phenotest’ v1.34.0 package in R v3.6.3^136^. Significance was calculated using 10,000 permutations.

#### GSEA analysis using Gene Ontology and Reactome gene sets

Gene Ontology term annotation were obtained from ENSEMBL through Biomart (Ensembl 108; downloaded on 2022-12-21), and gene-term association with only author statement support (GO evidence codes ‘TAS’ and ‘NAS’) or unclear evidence (GO evidence code ‘ND’) was filtered out. The Reactome annotations were obtained from the Molecular Signature Database mouse v2022.1^149^. For RNA-seq GSEA analysis, differential gene regulation information and gene-centric statistics were obtained directly from the DESeq2 result table without further processing. For ATAC-seq and H3K4me3 CUT&RUN GSEA analyses, the peak-centric DESeq2 result tables were first processed to generate gene-centric results compatible with GSEA. Specifically, only peaks with 10kb of annotated TSSs according to HOMER v5.1 ‘annotatePeaks.pl’ script^94^ were retained. For ATAC-seq, to capture the most strongly remodeled regulatory elements, the gene-centric data was obtained by capturing the DESeq2 statistics for the peak with the most significant DESeq2 FDR, and in the case of peaks tied for FDR, with the largest absolute log_2_(fold change) with aging among tied peaks for each annotated gene. For H3K4me3 CUT&RUN, to capture the peak most representative of gene promoters (since H3K4me3 is a modification typical of promoters), the gene-centric data was obtained by capturing the DESeq2 statistics for the peak closest to each annotated gene.

In all cases, from gene-centric result tables, the DESeq2 t-statistic was used to generate the ranked list of genes for functional enrichment analysis with aging in female and male macrophages separately. The ‘gsea’ function from phenotest’ was run on gene sets with a size of 20 to 5000 genes, filtering too small or too large gene sets, so as to improve biological robustness. For ease of reading, only the top 5 most significant sex-specific gene sets or top 10 most significant sex independent gene sets are shown in figures, with all shown gene sets passing FDR < 5%. All significantly regulated gene sets are reported in **Extended Data Table 3A-F**.

#### GSEA analysis of curated gene sets for top functional processes

For curated gene set analyses (**Figure 4e, Extended Data Figure 11a**), we used curated gene sets for validation of important top curated processes. Specifically, we curated candidate regulators of phagocytosis from CRISPR/Cas9 screens based on results reported by Haney et al. 2018 and Pluvinage et al. 2019^71,72^. We also obtained a set of genes upregulated in M1 and M2 *in vitro* polarized BMDMs, by reanalyzing a publicly available RNA-seq dataset (accession number GSE103958)^75^. Briefly, downloaded FASTQ reads were mapped using STAR, a count table generated using featureCounts from subread, and differential gene expression analysis was performed using DESeq2 comparing M0 (untreated) vs. M1 (IFNg LPS treated) BMDMs, and M0 (untreated) vs. M1 (IL4 or IL13 treated) BMDMs. Genes with FDR < 5% and upregulated upon M1/M2 treatments were used as signatures of polarization for analysis (M1: 1392 genes; M2: 760 genes; **Extended Data Table 4B**).

#### GSEA analysis of macrophage-regulated gene sets

To compare the impact of various interventions on macrophage transcriptomes to that of aging in peritoneal macrophage from female mice, we also leveraged a GSEA approach. Specifically, we used our various RNA-seq experiments to identify genes significantly regulated in response to *in vivo* interventions (*i.e*. OVX vs. Sham, E2-supplementation vs. Vehicle, *Esr1* KO) or *in vitro* modulation (*i.e*. TF gene shRNA knockdown; see below).

In all cases, we performed DESeq2 differential gene expression analysis of our datasets to identify genes with significant regulation across our conditions (see above). For RNA-seq datasets measuring the impact of *in vivo* interventions on primary peritoneal macrophage transcriptomes and for *in vitro* E2 supplementation of J774A.1cells, we obtained gene sets corresponding to genes significantly upregulated or downregulation in response to the intervention at FDR < 5%. These lists were then used as gene sets for GSEA analysis with ‘phenotest’ (**Figure 5c, h; Extended Data Figures 13e, 14c, 15e**). For *in vitro* modulation of TF gene expression datasets, we used a more stringent definition for target gene sets to limit the impact of hairpin-specific off-target effects. For each candidate TF, we performed differential expression analysis comparing control (i.e. non-mammalian and luciferase hairpins) to each hairpin separately. To identify robust and consistent transcriptional targets, we then performed Fisher combination of adjusted p-values across both hairpins (analytical tables provided in **Extended Data Table 6C-H**). Genes were considered to be significant transcriptional targets of a candidate TF if and only if (i) they showed consistent direction of transcriptional regulation across both hairpins, and (ii) if their Fisher combined adjusted p-values were < 10^-4^ as a stringent filter. Heatmaps showing consistent regulation for the chosen gene sets to be used for GSEA analysis are reported in **Extended Data Figure 17e**, showing that our approach identifies consistent transcriptional targets. These lists were then used as gene sets for GSEA analysis with phenotest (**Figure 6f**).

### Culture of primary peritoneal macrophages and BMDMs

All primary macrophages were cultured in a humidified incubator under 5% CO_2_-95% air at 37°C. MACS-purified peritoneal macrophages were cultured in macrophage media (DMEM/F-12 50/50 supplemented with 10% FBS, 1x antibiotic antimycotic (Gibco #15240-062), 10% L929 conditioned media and 2ng/mL M-CSF (Miltenyi Biotec #130-101-706)) in non-treated tissue culture plates. Bone marrow cells were obtained from the hind legs of aged C57BL/6JNia, aged C57BL/6J and estrus cycle tracked C57BL/6NTac mice following as before^69^. Briefly, after bone marrow harvest following Amend et al. 2016 protocol^69,150^, cells were resuspended in 1mL of MACS resuspension buffer and treated with 1x RBC lysis solution for 2 minutes at room temperature, then centrifuged at 300g for 10minutes at 4°C. Resuspended RBC-lysed bone marrow cells were then filtered on a 70mM filter to remove debris and clumps (Miltenyi Biotec #130-110-916). Cells were then transferred to macrophage media for culture in T75 vented untreated tissue culture flasks (Greiner Bio-One #658195) for differentiation into bone marrow derived macrophages. After 3 days, the media was replaced with fresh macrophage media and differentiation continued for a total of seven days after euthanasia and bone marrow harvest. Obtained BMDMs were then used for downstream experiments (*i.e.* phagocytosis assay (**Extended Data Figure 8b**) and Agilent Seahorse metabolic profiling (**Figure 4j, k, Extended Data Figure 10a, b**).

### Growth and maintenance of cell lines

All cell lines were cultured in a humidified incubator under 5% CO_2_-95% air at 37°C. L929 cell line (ATCC CCL-1) were cultured with Dulbecco’s Modified Eagle’s Medium/ Hams F-12 50/50 mix (DMEM/F-12 50/50; Corning #10-090-CV) supplemented with 10% fetal bovine serum (FBS) (Sigma-Aldrich #F0926), and 1x penicillin streptomycin L-glutamine (Corning # 30-009-Cl). L929 cells were seeded in vented T175 tissue culture flasks (VWR #10062-864) at 10% confluency. Once cells reached around 80% confluency, conditioned media was collected and filtered through a 0.45µM filter (GenClone #25-246). Conditioned media was stored at -20°C until use (see below). HEK293T (ATCC CRL-3216), RAW264.7 (ATCC TIB71), and J774A.1 (ATCC TIB-67) were cultured with Dulbecco’s Modified Eagle’s Medium (DMEM) (Corning #15-013-CV) supplemented with 10% FBS, and 1x penicillin streptomycin L-glutamine. Media was changed every 2-3 days. To passage cell lines, L929, HEK293T, and RAW264.7 cells were detached using 0.05% trypsin (Coring #25-052-Cl) for 5 minutes at 37°C. After incubation, trypsin was neutralized by adding double the volume of complete media to the trypsin used. Cells were pelleted by centrifugation at 300g for 5 minutes at room temperature. For passaging of J774A.1 cells, cells were detached using a cell lifter (VWR #76036-006). Cells were collected and pelleted at 300g for 5 minutes at room temperature. Supernatant was discarded and cells were resuspended with 1mL of complete media. Cells were seeded at lower densities in a new tissue culture plate as needed based on experimental needs. All cell lines were re-authenticated after establishing culture in the laboratory using the ATCC cell line STR profiling service to confirm their identity in presented experiments.

### 17β-estradiol treatment of J774A.1 female macrophage cell line

To study the impact of estradiol responses on the J774A.1female macrophage cell line, we first generated hormone-depleted Fetal Bovine Serum (FBS). A total of 50 mL of FBS (Sigma-Aldrich #F0926) was mixed with 126.5 mg of dextran-coated charcoal (Sigma, #C6241-5G), and vortexed to ensure thorough mixing. The mixture was then incubated at 4°C for 12 hours. Following incubation, the stripped BSA was sequentially filtered using 0.45μm and 0.22μm polyvinylidene fluoride (PVDF) filters to remove residual particulates.

A 1nM stock solution of 17β-estradiol (E2) was prepared by dissolving 2.5 mg of 17β-estradiol in 100% Ethanol. Cells were cultured in hormone-stripped media (DMEM with 10% stripped FBS and 1% penicillin streptomycin L-glutamine (Corning # 30-009-Cl)) for one passage in a 10 cm dish before being transferred to stripped media supplemented with (i) 17-β-estradiol (E2) at a final concentration of 100nM or (ii) an equivalent volume of 100% EtOH as the vehicle control. J774A.1cells were cultured for 48 hours with under these conditions before being harvested for mRNA-seq or profiled for phagocytic capacity by flow cytometry assay (**Extended Data Figure 14a**).

### Analysis of macrophage phagocytic capacity

We used our previously described microscopy-based phagocytosis assay protocol^69^ and a new streamlined flow cytometry-based phagocytosis assay protocol to quantify the phagocytic capacity of primary macrophages and macrophage cell lines.

Briefly, for the microscopy protocol, 75,000 macrophages were seeded in a 24-well plate with a 12mm glass coverslip (Electron Microscopy Sciences #72231-01), with technical duplicates for each animal. Macrophages were allowed to attach overnight, before treatment for 1 hour with 1mg/mL Alexa Fluor 488-labelled Zymosan bioparticles (Invitrogen #Z23373) diluted in macrophage media. After 1-hour, Zymosan particles were washed three times with ice-cold DPBS, and cells were fixed with 4% formaldehyde (Macron #H121-05) for 10 minutes at room temperature. After fixation, coverslips were washed three times with DPBS and mounted on microscope slides (VWR #48311-703) with prolong diamond antifade mounting medium with DAPI (Invitrogen #P36962). Images of phagocytosis were acquired using the Keyence microscope (Keyence #BZ-X710) with the 10x objective. Stitched images of a 3 by 3 field were then counted by blinded experimenters. To quantify Zymosan-internalized (green fluorescent) cells and total cell number, images were analyzed using ImageJ software^151^. Total cell counts were determined by identifying DAPI-stained nuclei using the “Analyze Particles” function following appropriate thresholding. Due to the possibility of multiple phagosomes within a single cell, Zymosan-positive cells were manually counted to ensure accurate identification of individual phagocytic cells The phagocytosis index was determined by calculating the ratio of Alexa Fluor 488–positive cells to the total number of cells. Phagocytosis images for aging and estrus cycling has been deposited to Figshare under 10.6084/m9.figshare.29286194 and 10.6084/m9.figshare.29286206, respectively.

For our new flow cytometry-based protocol, 100,000 macrophages were first blocked with FcR-blocking reagent for 10 minutes at 37°C. For primary MACS-purified peritoneal macrophages, we also included an additional staining step with F4/80-APC (Miltenyi Biotec #130-116-547) for downstream data filtering. For both primary and immortalized macrophages, cells were then treated with Alexa Fluor 488-labelled Zymosan bioparticles as above, targeting with a multiplicity of infection (MOI) of 2 per macrophage for 30 minutes at 37°C. After treatment, macrophages were fixed with a final concentration of ice cold 2% formaldehyde solution (Macron #H121-05) at 4°C protected from light. Macrophages were washed twice and resuspended in 500uL of MACS resuspension buffer, then analyzed on the MACSQuant Analyzer 10 to phagocytosis capacity. Gating was determined using fluorescence-minus-one [FMO] controls for each color used in the experiment to ensure that positive populations were solely associated with the antibody/particle Analysis was performed in Flowlogic V8 software. Flow cytometry data for aging, sham vs. OVX, E2 supplementation, *Esr1* KO, RAW264.7 phagocytosis regulator knockdown, RAW264.7 transcription factor candidate knockdown, and J774A.1 Irf2 knockdown phagocytosis has been deposited to Figshare under 10.6084/m9.figshare.29161955, 10.6084/m9.figshare.29169836, 10.6084/m9.figshare.29169821, 10.6084/m9.figshare.29169860, 10.6084/m9.figshare.29169821, 10.6084/m9.figshare.29169872, 10.6084/m9.figshare.29169899, and 10.6084/m9.figshare.29286278, respectively.

### Cell surface marker protein level quantification by flow cytometry

To estimate potential differences in peritoneal macrophage surface markers (*i.e*. Zymosan pattern recognition receptors, M1/M2 markers) with respect to sex and age, 100,000 MACS-purified peritoneal macrophages were subjected to Fc receptor blocking (Miltenyi Biotec #130-092-575) and stained with corresponding antibodies according to the manufacturer’s instructions. For analysis of pattern recognition receptors, cells were stained with F4/80-Brilliant Violet 421 (1:20) (Biolegend #123131), Dectin-1-PE (1:10) (Miltenyi Biotec #130-102-284), and CD282 (Tlr2)-APC (1:50) (Miltenyi Biotec #130-120-052). For M1 marker detection, cells were stained with CD80-FITC (1:20) (Biolegend #104705) and CD86-APC (1:20) (Biolegend #105011). For M2 marker detection, cells were stained with CD163-PE (1:20) (Biolegend #111803) and CD206-APC (1:20) (Biolegend #141707). After gating on F4/80 positive cells, we obtained median fluorescent intensity (MFI) of stained cells for surface proteins of interest using flow cytometry quantification on the Milteni Biotec MACSQuant Analyzer 10. Flow cytometry results were processed using the FlowLogic V8 software. Flow cytometry data has been deposited to Figshare under 10.6084/m9.figshare.29162144.

### Metabolic profiling of macrophages using the Agilent Seahorse platform

50,000 BMDMs or transduced macrophage cell lines (RAW264.7, J774A.1) cells were seeded in a 96-well Seahorse assay plate with macrophage media (or appropriate culture media) and incubated overnight for cells to attach. To characterize the bioenergetics of cells, a XF–96 Flux Analyzer (Agilent Technologies) was used. Growth media was replaced with 175μL of XF assay medium (specially formulated, unbuffered Dulbecco’s modified Eagle’s medium for XF assay; Agilent Technologies #103575-100) supplemented with 1mM sodium pyruvate at pH 7.4 (Agilent Technologies #103578-100). The cells were kept in a non-CO_2_ incubator for 60 minutes at 37°C before placing in the analyzer. Oxygen consumption rate [OCR] and extracellular acidification rate [ECAR] were measured using 3 cycles of 3 minutes wait and 4 minutes measure time. Each biological replicate had three technical replicates and the results obtained were averaged. Injection of compounds and calculating metabolic parameters were obtained by following an established protocols for macrophages^74^.

To normalize OCR and ECAR values to actual live/seeded cell number, the cell media was removed from 96-well plates at the end of the assay and air died for around 10 minutes before storage at - 80°C at least overnight to initiate cell lysis. To finalize cell lysis, 100mL of nuclease-free water was added to each well and incubated at 37°C for 1 hour, and plates were refrozen for > 1 hour at -80°C. Plates were thawed at room temperature, and 200mg/mL Hoechst (Thermo Scientific #H1399) in TNE buffer (10mM Tris [Thermo scientific #J22675-A1], 1mM EDTA [VWR #BDH7830-1], 2M NaCl [Sigma-Aldrich #S5150-1L], pH 7.4) was added to each well. Plates were read at excitation 346 and emission 460 on Molecular Devices SpectraMAX M3. Fluorescent values were then used to divide the OCR and ECAR values to obtain normalized, comparable values. Seahorse data for aging, estrus cycling, transcription factor candidate knockdown in RAW264.7, and *Irf2* knockdown in J774A.1 has been deposited under 10.6084/m9.figshare.29169911, 10.6084/m9.figshare.29169920, 10.6084/m9.figshare.29169926, and 10.6084/m9.figshare.29169938, respectively.

### Measurement of macrophage glycolytic function using Lactate-Glo

30,000 MACS-purified peritoneal macrophages or RAW264.7/J774A.1 cells were seeded into a 96-well plate tissue culture plate (GenClone #25-109) with 100mL of fresh media. Then, 48 hours after seeding (primary peritoneal macrophages) or 24 hours (macrophage cell lines), the conditioned media was collected and stored at -20°C until further processing. To note, for this experiment, transduced RAW264.7/J774A.1 cells were cultured without puromycin (see below) to avoid metabolic abnormalities due to the impact of puromycin on mitochondrial function^152^. To measure the amount of lactate in the condition media, Lactate-Glo (Promega #J5021) was used following to the manufacturer’s protocol and luminescence values were collected using Promega GLOMAX (Promega #GM3500) plate reader. Luminescence values were normalized to actual live/seeded cell numbers using the same method as for the Seahorse assay (see above).

### Lentiviral particle production for shRNA-mediated gene knockdown

In all cases, plasmid DNA was isolated from overnight cultures using mini-preps from the Macherey-Nagel NucleoSpin Plasmid kit (Macherey-Nagel #740588.250) according to the manufacturer’s protocol, and DNA was sent to GENEWIZ or Plasmidsaurus for sequence validation before use in any experiment.

We used a lentiviral shRNA particle approach to mediate gene-specific RNA interference and evaluate the impact of decreased gene expression of candidate phagocytosis regulators or TF-mediators on macrophage phenotypes. We obtained 3^rd^ generation of lentiviral vector packaging backbones from Addgene: pMDLg/pRRE (Addgene #12251), pRSV-Rev (Addgene #12253), VSV.G (Addgene #14888). To guarantee single clone origin of laboratory isolates, bacteria were re-streaked on a 100mg/mL carbenicillin (Sigma-Aldrich #C3416) Luria broth (LB; Thermo Scientific #J75852-A1) 1% agar (Invitrogen #30391-023) plates and incubated overnight at 37°C. For plasmid preparation, single colonies were isolated and inoculated into culture tube containing 100mg/mL carbenicillin in LB media for culture at 37 °C with shaking for 16-20 hours for each construct. Plasmid DNAs were obtained using the NucleoBond Xtra midi plus EF kit (Macherey-Nagel #740422.50) following manufacturer’s recommendation.

MISSIONâ shRNA control and target bacterial glycerol stocks were obtained from Sigma-Aldrich (**Extended Data Table 1H**). To obtain single bacteria colonies, frozen glycerol stocks were inoculated into sterile culture tubes (Genesee Scientific #21-129) containing 0.5mL terrific broth (TB) (Sigma-Aldrich #T0918-1KG) without antibiotics and incubated at 37 °C with shaking for 30 minutes. Next, 50mL of bacteria pre-cultures was spread on prewarmed 100mg/mL carbenicillin TB 1% agar plate, and incubated for 16 hours at 37 °C. For plasmid preparation, a single colony was isolated from the plate and inoculated into a culture tube containing 100mg/mL TB, incubated at 37 °C with shaking for 16-20 hours for each construct. To produce a large quantity and endotoxin free plasmid DNA, 200mL of 100mg/mL TB bacteria culture was inoculated and incubated at 37°C with shaking for 20 hours. Plasmid DNAs were obtained using the NucleoBond Xtra midi plus EF kit (Macherey-Nagel #740422.50) following manufacturer’s recommendation.

Lentiviral particles were produced by transfection of HEK293T cells using the calcium phosphate transfection method^153^. Transfection mixtures were prepared by combining plasmid DNA (contained pMDLg/pRRE (1,000 ng), VSV.G (500 ng), pRSV-Rev (500 ng), and the pLKO shRNA plasmid of interest (1000ng)), with 2.5 M calcium chloride (Sigma-Aldrich #C1016-500G) and mixing with 0.22mM filter sterilized 2xHEPES-buffered saline ([HBS] 140mM NaCl, 1.5mM Na_2_HPO_4_·2H_2_O (VWR #0348-500G), 50mM HEPES)) to form DNA-calcium phosphate complexes. The DNA-calcium phosphate complexes were formed for 2 minutes at room temperature and then added dropwise on HEK293T cells at 60-80% confluency. HEK293T were incubated with DNA-calcium phosphate complexes for 16 hours, then media was changed to fresh complete media. After 24 hours, supernatants containing viral particles were collected and filtered using a 0.45mM filter to remove cells.

To concentrate lentiviral particles before use, supernatant was carefully layered on top of a layer of 20% sucrose (Sigma-Aldrich #S0389-1KG) in DPBS solution in a Beckman Coulter polypropylene quick-seal centrifuge tube (Beckman Coulter #343414). Viral preps were centrifuged at 70,000 x g for 2 hours at 4°C in an Optima XPN-100 Ultracentrifuge (Beckman Coulter #B10053). After centrifugation, the layered supernatant was removed, and viral pellets were resuspended and concentrated 100x in ice cold DPBS. Lentiviral particles were aliquoted and stored at -80°C until experiments.

### Lentiviral transduction of macrophage cell lines

Macrophage cell lines were seeded at a density of 30,000 cells per well in a 96-well tissue culture plate (GenClone #25-109) and incubated for 48 hours. During this period, cells were treated with various concentrations of puromycin in complete media (10, 8, 6, 4, 2,1mg/mL) (Sigma-Aldrich #P8833), to assess cytotoxicity. On the second day, 100μL of the culture medium in each well was replaced with fresh medium containing the corresponding concentration of puromycin. Following the manufacturer’s instructions, CyQUANT™ XTT Cell Viability (Invitrogen #X12223) assay reagents were added to each well to assess cell viability. The plates were then incubated for 4 hours at 37 °C. Absorbance was measured at 450 nm and 660 nm using a microplate reader. Specific absorbance values were calculated by subtracting the 660 nm reference readings from the 450 nm readings.

For transduction, 200,000 cells from macrophage cell lines (RAW264.7, J774A.1) were seeded in a 12-well tissue culture plate (GenClone #25-109) one day before infection. The next day, cells were infected with viral particles at a MOI of 1 in the presence of 8mg/mL polybrene (Sigma-Aldrich #TR-1003-G). After 24 hours, the media was replaced with fresh complete media. 48 hours after infection, cells were selected using complete media with addition of 10mg/mL of puromycin (Sigma-Aldrich #P8833), as determined with our in house kill curve for these cells (see above). Molecular and/or functional profiling of modified cells was performed on puromycin-selected macrophage cells cultured in complete media supplemented with 10mg/mL puromycin, expect for metabolic assays where 24 hours prior, media was changed to complete media without puromycin.

### Reverse transcription quantified PCR verification of shRNA gene knockdown

Total RNA was reverse transcribed into complementary DNA (cDNA) using the Thermo Scientific™ Maxima H Minus First Strand cDNA Synthesis Kit (Thermo scientific #K1682), according to the manufacturer’s instructions. Quantitative PCR was performed using the SensiFAST SYBR No-ROX Kit (Bioline #BIO-98020) with gene-specific primers (For sequences see **Extended Data Table 1I**) to assess relative gene expression. cDNA, primers, and SYBR master mix were first combined in a PCR tube to ensure a homogeneous mixture, then transferred into Mic qPCR-specific tubes (QuantaBio #95910-20) for amplification using the Mic qPCR Cycler (Bio Molecular Systems #MIC-2). Gene expression levels were quantified using the ΔΔCt method and normalized to the geometric mean of three housekeeping genes (*18s*, *Hprt*, and *Ubc*).

### Vaginal cytology determination of estrus cycling

Vaginal lavages were obtained using a standard protocol^154^. Lavages were deposited to glass slides (VWR #48311-703) and left to dry for at least 24 hours. Vaginal cytology slides were stained for 30 seconds 1% Thionin (Alfa Aesar #A18192), in 20mM acetic acid (Sigma-Aldrich #695092-2.5L-GL), 0.36mM Sodium Hydroxide (Sigma-Aldrich #221465-500G) at pH 4.0. Excess thionin staining was washed twice in MilliQ water. Slides were dehydrated for 30 seconds in 70%, 95%, and 100% ethanol, then air dried overnight before microscopic analysis. Slides were viewed under a microscope at 20x magnification identify the estrus phase each mouse was in at the time of sample collection (**Extended Data Table 1C**).

### TF motif analysis in chromatin regions remodeled as a function of aging and sex

To identify potential TFs regulating sex-specific aging phenotypes in macrophages, ATAC-seq and H3K4me3 peaks with age-related remodeling in only one sex were obtained (see above for criteria). Motif finding was performed using HOMER v5.1 ‘findMotifsGenome.pl’ script^94^, using all robustly detected metapeaks as enrichment background and the mm10 reference genome. We focused on results for enrichment of known motifs reported in “knownResults.txt”. Top TF motifs corresponding to TF genes significantly regulated with female aging are shown in **Figure 6c, d**. All enriched motifs from our analyses are reported in **Extended Data Table 7A, B.** To confirm enrichment of Irf2-related binding motifs, Irf2 peaks (see below) were used with the same pipeline, and selected motifs are reported in **Extended Data Figure 20a, Extended Data Table 7C, D**.

### ChromVAR analysis of TF footprint accessibility in bulk ATAC-seq data

As an independent method to evaluate candidate TF chromatin binding in aging female and male macrophages, we leveraged the ‘chromVAR’ pipeline{Schep, 2017 #172}, through the R packages ‘ChrAccR’ 0.9.17 and ‘chromVAR’ 1.8.0, together with HOMER motifs from ‘chromVARmotifs’ 0.2.0. This paradigm identified TF footprints with significant variation in occupancy based on computed Z-scores across conditions using the ‘chromVAR’ FDR calculation paradigm (**Extended Data Table 6A**).

### SCENIC analysis of regulon activity in single cell RNA-seq analysis

As an orthogonal method to evaluate candidate TF activity in aging female and male macrophages, we took advantage of macrophages identified in our scRNA-seq datasets (see above) and applied the SCENIC algorithm to our data^96^. We applied the SCENIC pipeline using R packages SCENIC v1.1.2-01, RcisTarget v1.6.0, GENIE3 v1.8.0 and AUCell v1.8.0 following the package tutorial in R v3.6.3^136^. Specifically, SCT normalized UMI counts from a combined Seurat object with macrophages transcriptomes from our 3 datasets were used as the input expression matrix. Mouse feather datasets for the mm10 assembly were downloaded from the SCENIC website (*i.e*. mm10 refseq-r80 500bp_up_and_100bp_down_tss.mc9nr.feather and mm10 refseq-r80 10kb_up_and_down_tss.mc9nr.feather). Computed TF regulon activities were obtained for 82 TFs in every cell, and we used a Wilcoxon rank-sum test approach to determine whether regulon activity was significantly impacted by aging in female or male macrophages (**Extended Data Table 6B**). Dot plots and violin plots were generated for TFs also identified from our bulk ‘omic’ data (**Extended Data Figure 16c, d**).

### Irf2 protein level quantification using intracellular labeling and flow cytometry quantification

Cells were resuspended in 90μL of staining buffer and incubated with 10μL of FcR blocking reagent (Miltenyi Biotec #130-092-575) at 4 °C for 20 minutes. For MACS-purified peritoneal macrophage analysis, surface F4/80 staining was performed (to guarantee quantification of Irf2 expression solely on macrophages) by adding 2μL of anti-F4/80 antibody per sample, followed by a 10-minute incubation at 4 °C. Then, 2mL of resuspension buffer was added, and cells were centrifuged at 300g for 10 minutes to remove unbound antibodies. Supernatant was then removed, and cells were resuspended in 1mL of Foxp3 Fixation/Permeabilization working solution (Invitrogen #00-5523-00), pulse-vortexed, and incubated at 4 °C for 1 hour, protected from light. Following fixation, 2 mL of 1x Permeabilization Buffer (Invitrogen #00-5523-00) was added, and samples were centrifuged at 600g for 5 minutes at room temperature. Supernatants were carefully decanted, leaving approximately 200μL, and cells were gently resuspended. To block nonspecific binding, 10μL normal rat serum was added, and samples were incubated at room temperature for 15 minutes. Each sample was split into 98μL aliquots for intracellular staining with 2μL of directly conjugated PE-IgG isotype control (Abcam #ab209478) or PE-Irf2 (Abcam #ab318372) antibodies, and incubated at 4 °C for 1 hour, protected from light. After staining, cells were washed twice with 2mL of 1x Permeabilization Buffer by centrifugation at 600g for 5 minutes at room temperature. Finally, cells were resuspended in eBioscience Flow Cytometry Staining Buffer (Invitrogen #00-4222-26). Stained cells’ median fluorescent intensity (MFI) was analyzed by flow cytometry on the Miltenyi Biotec MACSQuant Analyzer 10, and flow cytometry results were processed using the FlowLogic V8 software. To obtain the TF expression level, non-specific background signal was removed by subtracting the MFI of IgG isotype control from the MFI of Irf2 for each specific cell sample (i.e. isolate for a single animal, or transduced cell aliquot)^155^. TF levels were then normalized by dividing the median of the entire cohort per value to mitigate the impact of cohort-specific variation in fluorescence intensity. Raw flow cytometry data for aging, RAW264.7 *Irf2* KD, J774A.1 has been deposited to Figshare under 10.6084/m9.figshare.29285708, 10.6084/m9.figshare.29169947, and 10.6084/m9.figshare.29169959 respectively.

### Active Motif FactorPath Irf2 ChIP-seq in young female BMDMs

Red-blood cell lysed bone marrow cells from 4 months old female C57BL/6JNia mice were differentiated into BMDMs in two independent biological replicates (pool of 2 animals per replicate) as above. For each replicate ChIP experiment from an independent primary culture, we plated 10 million BMDMs, which were then fixed by adding 1:20 (v/v) ratio of freshly prepared formaldehyde solution (11% formaldehyde (Sigma-Aldrich #F8775-25ML), 100 mM NaCl,1 mM EDTA pH 8.0 (EMD Millipore #324506), and 50 mM HEPES pH = 8) to the cell culture media, followed by incubation for 15Lmin at room temperature. Fixation was stopped by adding 2.5 M glycine (Sigma-Aldrich #G7403-100G) solution at a 1:20 (v/v) ratio and incubating for 5 min at room temperature. Cells were collected by scraping on ice and spun down at 4°C 800g for 10 minutes. All subsequent steps were performed on ice. Cells were collected and washed twice with chilled PBS (Sigma-Aldrich #P4474-500ML) with 0.5% Igepal (Sigma-Aldrich #I8896-50ML). 1mM of phenylmethylsulfonyl fluoride (PMSF) (Sigma-Aldrich #P7626-250MG) was added to the second wash before pelleting. Cell pellets were snap-frozen on dry ice and sent to Active Motif for further processing.

Subsequent processing steps, including chromatin extraction, fragmentation, antibody precipitation, input, and library preparation, were performed by Active Motif (Carlsbad, CA) using a ChIP validated antibody to IRF2 (Abcam #ab124744). Active motif generated ChIP-seq libraries from the ChIP-DNA and a pooled Input control library using a custom Illumina library type protocol on an automated system (Apollo 342, Wafergen Biosystems/Takara). A pooled input library was also generated to provide a background for peak calling. ChIP-seq and input libraries were sequenced on Illumina NovaSeq 6000 as 75 nucleotides single-end reads. Demultiplexed FASTQ files were provided by Active Motif, and the ChIP-seq FASTQ reads were deposited to SRA under Bioproject PRJNA1126274.

### Female BMDM Irf2 ChIP-seq data processing and peak calling

Single-end Irf2 ChIP-seq reads were mapped to the mm10 genome reference using bowtie1 v1.2.3^156^. After alignment, PCR duplicates were removed using the FIXSeq algorithm^157^, as previously described^133^. We used HOMER v4.10.4 findPeaks with the “-style factor” option, and provided the pooled input reads with “-i”, to identify Irf2-enriched regions in our BMDM ChIP-seq replicate libraries^94^. Consensus peaks found in both replicates were identified using MSPC v5.5.0^158^, which identified 460 reproducibly bound regions (**Extended Data Table 7A**). Peaks were annotated to the closest TSS using the HOMER ‘annotatePeaks.pl’ script^94^ (**Extended Data Table 7A**), and enriched DNA motifs were identified using the HOMER ‘findMotifGenome.pl’ script^94^ (**Extended Data Figure 20a**, left panel, **Extended Data Table 7C**).

Genome browser bedgraph files to visualize signal were generated using the HOMER ‘makeUCSCfile’ script^94^, including input normalization and the “-style chipseq” option.

### Peritoneal macrophage isolation and preservation for CUT&RUN

Primary peritoneal macrophages were purified from 4 and 20 months old female C57BL/6J using MACS. Equal number of cells with purity >90% from 2-3 female C57BL/6J mice were used to generate independent pools of young and of old macrophages. Macrophages were combined from independent animals to obtain >500,000 macrophages per independent pool. After pooling, samples were recounted using the CellDrop counter (DeNovix), before resuspension in 1mL of Synth-a-Freeze cryopreservation media (Gibco #A1254201), to avoid biologic material contributions from FBS. Cells were slowly frozen for cryopreservation using a Mr. Frosty freezing container (ThermoFisher Scientific #5100-0001) for 24 hours at -80 °C, before moving into liquid nitrogen for storage prior to shipment. Final cell numbers were: young female pool 1 (n=2 mice): 1.14x10^6^ macrophages, young female pool 2 (n=2 mice): 1.52x10^6^ macrophages, old female pool 1 (n=3 mice): 0.51x10^6^ macrophages, and old female pool 2 (n=2): 0.86x10^5^ macrophages.

### Irf2 CUT&RUN and library generation for primary peritoneal macrophages

Cryopreserved peritoneal macrophage pools from young and old female C57BL/6J were sent to Active Motif (Carlsbad, CA) for the CUT&RUN assay using the same ChIP-validated antibody to IRF2 (Abcam #ab124744). Each cryopreserved sample comprised at least 500,000 MACS-purified peritoneal macrophages. Since our data suggested that total IRF2 levels were significantly reduced in aging female peritoneal macrophages, CUT&RUN reactions were designed to include Active Motif’s CUT&RUN Spike-In Control.

After nuclei isolation by Active Motif, peritoneal macrophage nuclei were combined with *Drosophila melanogaster* spike-in nuclei (Active Motif #53183) at a 1:100 ratio. In particular, in each CUT&RUN reaction, the Active Motif team included a cognate spike in antibody (anti-H2Av) to target the *Drosophila* nuclei. All subsequent steps, including chromatin extraction, fragmentation, antibody precipitation, input, and library preparation, were performed by Active Motif. CUT&RUN libraries were sequenced on Illumina NovaSeq 6000 as 38 nucleotides paired-end reads. Demultiplexed FASTQ files were provided by Active Motif, and corresponding FASTQ reads were deposited to SRA under Bioproject PRJNA1377026.

### Female peritoneal macrophage Irf2 CUT&RUN data processing and peak calling

Paired-end Irf2 CUT&RUN reads were mapped separately to the mm10 genome reference (sample) and dm6 (*Drosophila melanogaster* spike-in reads) using bowtie2 v2.5.4^144^. After alignment, PCR duplicates were removed for each set of reads using the ‘rmdup’ function of samtools 0.1.19^145^. We used HOMER v4.10.4 findPeaks with the “-style factor” option to identify Irf2-bound regions in each CUT&RUN samples^94^. Consensus robust peaks were identified using MSPC v5.5.0^158^, which identified 1565 reproducibly bound regions. Peaks were annotated to the closest TSS using the HOMER ‘annotatePeaks.pl’ script^94^ (**Extended Data Table 7B**), and enriched DNA motifs were identified using the HOMER ‘findMotifGenome.pl’ script^94^ (**Extended Data Figure 20a**, right panel, **Extended Data Table 7D**).

To take into account spike-in signal in track generation, scaling factors were computed using ratios of mapped reads as evaluated by samtools ‘flagstat’ from mm10 (signal) to dm6 (spike-in), as recommended ^159,160^. Genome browser bedgraph files to visualize signal were generated using the HOMER ‘makeUCSCfile’ script^94^, including the “-style chipseq” option and including computed spike-in scaling factors using the “-norm” option.

### Functional enrichment analysis of Irf2-bound regions in BMDMs and peritoneal macrophages

To compute functional enrichment for Irf2 target genes, we leveraged the ‘GREAT’ 4.0.4 tool^98^, a tool specifically optimized to identify functional enrichment starting from genomic regions. The MSPC peaks for BMDM ChIP-seq or peritoneal macrophage CUT&RUN were used as foreground, and the mm10 genome was used as the background for enrichment, with default parameters for all other options. Results were exported as ‘tsv’, processed in R to filter significant enrichments with FDR < 5% and are reported in **Extended Data Table 7E, F**. We also used R package ‘OmicCircos’ v1.24.0^161^ to visualize genomic distribution of peaks along chromosomes (**Extended Data Figure 20b**), ‘ChIPSeeker’ v1.22.1^162^ and ‘TxDb.Mmusculus.UCSC.mm10.knownGene’ v3.10.0 to analyze distribution of peaks relative to genes (**Extended Data Figure 20c**), together with clusterProfiler v3.14.3 as an orthogonal approach to GREAT for functional enrichment analysis of Irf2-bound regions (see results in **Extended Data Table 7G, H**). To evaluate the overlap between Irf2 ChIP-seq and CUT&RUN peaks, we used ChIPpeakAnno v3.20.1^163^ to draw Venn diagrams and perform significance of overlap analyses using the hypergeometric distribution. As recommended by the package developers, we used the number of genome-wide putative Irf2 binding sites in mm10 as background, as determined using the HOMER Irf2 motif PWM and ‘scanMotifGenomeWide.pl’ (yielding 144107 potential sites).

### Differential Irf2 binding analysis in aging female peritoneal macrophages

Since only the most recent versions of the package include options for spike-in normalization, differential CUT&RUN analysis was performed in R 4.5.1^136^ using ‘Diffbind’ v3.20.0 and ‘DESeq2’ v1.48.2, leveraging our mapped read bam files (both mm10 for signal and dm6 for spike-in control) over MSPC the consensus Irf2 peritoneal macrophage peaks. The mm10 blacklist was used to filter potential spurious peaks as recommended in the package vignette. Read counting, normalization and differential binding analyses were performed using Diffbind’s core functions, choosing the DESeq2-powered methods. Peaks with FDR <5% were considered to be differentially bound with aging in female peritoneal macrophages (**Extended Data Table 7I**).

### Hk3 protein level quantification using intracellular labeling and flow cytometry quantification

Cells were resuspended in 90μL of staining buffer and incubated with 10μL of FcR blocking reagent at 4 °C for 20 minutes. Surface staining was performed by adding 2μL of anti-F4/80 antibody per sample, followed by a 10-minute incubation at 4 °C for peritoneal macrophage analysis. 2mL of RSB was added, and cells were centrifuged at 300 × g for 10 minutes to remove unbound antibodies and resuspended in 100 ul RSB. 100 μL of IC Fixation Buffer (Invitrogen# 00-8222-49) and pulse vortex to mix. Incubate 60 minutes at 4 °C. 2mL of 1X Permeabilization Buffer was added and centrifuged at 400-600 x g for 5 minutes at room temperature. To block nonspecific binding, 10μL normal rat serum was added, and samples were incubated at room temperature for 15 minutes.

Each sample was split into 98μL aliquots for intracellular staining with 2μL of directly conjugated IgG (Proteintech # CL488-65124) or Hk3 (Proteintech #CL488-67803) antibodies, and incubated at 4°C for 1 hour, protected from light. After staining, cells were washed twice with 2mL of 1X Permeabilization Buffer by centrifugation at 600 × g for 5 minutes at room temperature. Finally, cells were resuspended in eBioscienceTM Flow Cytometry Staining Buffer (Invitrogen #00-4222-26). Flow cytometry results were processed using the FlowLogic V8 software. To obtain the transcription factor level, the background was removed by subtracting the MFI of Igg from the MFI of Hk3 ^155^. The transcription factor levels were normalized by dividing by the median of the entire cohort for each value.

## Data availability

Sequencing data has been deposited to the Sequence Read Archive, under Bioprojects PRJNA1094925, PRJNA1095555, PRJNA1096052, PRJNA1101936, PRJNA1126274, PRJNA1222371, PRJNA1313760, PRJNA1377026.

Raw flow cytometry data (10.6084/m9.figshare.28934213, 10.6084/m9.figshare.29169674, 10.6084/m9.figshare.29169737, 10.6084/m9.figshare.29169707, 10.6084/m9.figshare.28934198, 10.6084/m9.figshare.29169578, 10.6084/m9.figshare.29169581, 10.6084/m9.figshare.29169587, 10.6084/m9.figshare.29161955, 10.6084/m9.figshare.29169836, 10.6084/m9.figshare.29169821, 10.6084/m9.figshare.29169860, 10.6084/m9.figshare.29169821, 10.6084/m9.figshare.29169872, 10.6084/m9.figshare.29169899, 10.6084/m9.figshare.29285708, 10.6084/m9.figshare.29169947,10.6084/m9.figshare.29169959, and 10.6084/m9.figshare.29286278), microscopy images (10.6084/m9.figshare.29286194, 10.6084/m9.figshare.29286206), and Agilent Seahorse data (10.6084/m9.figshare.29169911, 10.6084/m9.figshare.29169920, 10.6084/m9.figshare.29169926) have been deposited to Figshare.

## Code Availability

R code for this study was run using R version 3.6.0-3.6.3. ‘Omic’ data processing scripts have been made available on the Benayoun lab GitHub repository at https://github.com/BenayounLaboratory/Macrophage_SexDim_Aging.

## Acknowledgements

We are grateful to Dr. Caleb Finch, Dr. Raphael Williams, Clayton Baker, Dr. Marc Vermulst, Sarah Shemtov, Allison Chae, Courtney Shen, Wyatt Web, Evelyn Navar, Emily K. Wang, Prakroothi S. Danthi, Amy Huang, Rochelle W. Lai, Guan J. Lai and Claire Chung for scientific insights and assistance during the preparation of this manuscript. We thank USC DAR veterinarians Dr. Lynlee M. Stevey-Rindenow and Dr. Bradley Ahrens for performing the ovariectomy and sham surgeries. We thank Hemal Mehta at the USC Leonard Davis school of Gerontology Seahorse core facility for helping run Seahorse experiments. We thank Dr. Didier Trono for providing the pMDLg/pRRE (Addgene #12251) and pRSV-Rev (Addgene #12253) plasmids. We thank Dr. Tannishtha Reya for providing VSV.G (Addgene #14888) plasmid. We thank Dr. Kenneth S. Korach for providing the B6N(Cg)-Esr1*^tm4.2Ksk^/J* (*Esr1* KO) mice^91^ (JAX #026176).

This work was supported by a Diana Jacobs Kalman/AFAR Scholarships for Research in the Biology of Aging, NIA T32 AG052374, and AthenaDAO student leader award to R.J.L.; USC Provost’s Undergrad Research Fellowships to E.E.W., E.H.L., and C.R.; GCRLE-2020 post-doctoral fellowship from the Global Consortium for Reproductive Longevity and Equality at the Buck Institute, made possible by the Bia-Echo Foundation to M.K.; NIAID R01 AI134987 to H.S.G.; core vouchers from the USC Leonard Davis School of Gerontology, NIA R00 AG049934, Pew Biomedical Scholar award #00034120, an innovator grant from the Rose Hills foundation, the Kathleen Gilmore Biology of Aging research award, a generous gift from Dr. Eric Hennigan, and Hevolution grant HF-GRO-23-1199072-28 to B.A.B. The authors acknowledge the Center for Advanced Research Computing [CARC] at the University of Southern California for providing computing resources that contributed to the research results reported within this publication (URL: https://carc.usc.edu). The funders had no role in study design, data collection and analysis, decision to publish, or preparation of the manuscript.

## Author contributions

R.J.L., N.S.K., and B.A.B. designed the study. R.J.L., N.S.K, A.P., performed live animal experiments. A.P. read vaginal cytology slides. R.J.L., S.C., M.K., L.A., I.Y.L., K.M., N.S.K., and B.A.B. dissected mouse tissues. R.J.L. and M.K. generated omics libraries. B.A.B. performed computational analyses. R.J.L. performed flow cytometry experiments. R.J.L., E.E.W., I.Y.L., C.R., J.J., and S.B. imaged and analyzed microscopy phagocytosis data. R.J.L., S.C., and A.X. performed phagocytosis flow cytometry and quantification analysis. R.J.L., S.C., and E.H.L. performed metabolic profiling experiments. R.J.L., A.X., and B.A.B analyzed Seahorse experiments. R.J.L., E.H.L, S.P., C.R., I.Y.L. performed RT-qPCR analyses. R.J.L., and S.C. perform intracellular Irf2 quantification. C.D.L. and H.S.G. contributed scientific insights to the project. R.J.L., S.C., and B.A.B. wrote the manuscript with input from all authors. All authors edited and commented on the final manuscript.

