## Supplementary material for "Mouse peritoneal macrophages undergo female-specific remodeling with aging driven by both hormone-dependent and -independent mechanisms"

**Extended Data material**

**Legends to Extended Data Figures**

**Extended Data Figure 1. Age- and sex-associated changes in peritoneal immune cell composition based on scRNA-seq analysis.**

(**a-d**) Cell type proportions differences according to scProportionTest as a function of age and sex for the combined (**a**), NIA 10x v2 (**b**), NIA 10x v3 (**c**), and JAX 10x v3 (**d**) datasets. Left-shifted cell types are more abundant in young animals and right-shifted cell types are more abundant in old animals. (**e**) Boxplots of peritoneal cell composition quantified by scRNA-seq; n = 8 libraries per sex and age; 4 independent cohorts in 3 independent datasets. For boxplots in panels (**e**), circles/squares represent NIA/JAX mice, respectively. Significance in non-parametric two-sided Wilcoxon rank-sum tests are reported in **e**. The center line of the box plots represents the sample median, the box limits consist of the 25^th^ and 75^th^ percentiles, the whiskers span 1.5x the interquartile range.

**Extended Data Figure 2. Age- and sex-associated changes in peritoneal immune cell composition based on flow cytometry.**

(**a**) Representative flow cytometry gating strategy for peritoneal lavage cell proportion analysis; B-cells are defined as CD19^+^; macrophages are defined as F4/80^+^; T-cells are defined as CD3^+^. (**b**) Boxplots of peritoneal cell composition quantified by flow cytometry; n ³ 25 per sex and age (6 independent cohorts). (**c**) Boxplots of peritoneal immune lavage cell number calculated by the proportion obtained from flow cytometry and total peritoneal lavage cell count; n ³ 25 per sex and age (6 independent cohorts). For boxplots in panels (**b, c**), circles/squares represent NIA/JAX mice, respectively. Significance in non-parametric two-sided Wilcoxon rank-sum tests are reported in **b, c**. The center line of the box plots represents the sample median, the box limits consist of the 25^th^ and 75^th^ percentiles, the whiskers span 1.5x the interquartile range.

**Extended Data Figure 3. Augur cell type prioritization analysis for peritonel cavity scRNA-seq.**

(**a-c**) UMAP of Augur AUC scores from scRNA-seq of NIA 10x v2 (**a**), NIA 10x v3 (**b**), and JAX 10x v3 (**c**) datasets. Values close to 0.5 denote smaller differences (random classification accuracy; blue shades), while larger values reflect larger differences (better classification accuracy; red shades). (**d-f**) Lollipop plot of Augur AUC results by immune cell type of NIA 10x v2 (**d**), NIA 10x v3 (**e**) and JAX 10x v3 (**f**). Not that there were too few enough monocytes (~100) in the NIA 10x v3 for Augur AUC score calculations.

**Extended Data Figure 4. Analysis of purity and identity of MACS-isolated mouse peritoneal macrophages.**

(**a**) Representative flow cytometry gating strategy for MACS-purified peritoneal macrophage (CD11b^+^ F4/80^+^) purity in young females. (**b**) Boxplot of MACS-purified peritoneal macrophage purity of ‘omic’ datasets from young and old female and male (yellow = RNA-seq, blue = ATAC-seq, purple = H3K4me3 CUT&RUN). Macrophage purity below 90% are excluded from any downstream ‘omic’ analysis. (**c**) Representative flow cytometry gating strategy for distinguishing small and large peritoneal macrophages [SPM/LPMs] in young female mice. SPM are defined as CD11b^+^F4/80^+Low^, while LPM are identified as CD11b^+^F4/80^+High^. (**d, e**) Quantification of peritoneal macrophage heterogeneity in NIA (**d**) and JAX (**e**) mice (n = 30 for young males, n = 28 for young females, n = 27 for old females, and n = 27 for old males; 7 independent cohort (JAX = 3 and NIA = 4)).

**Extended Data Figure 5. Differential gene and peak analysis in MACS-purified peritoneal macrophages as a function of age and sex.**

(**a**) Scatterplot of sex-independent transcriptomic aging changes, reported as DESeq2 log_2_ fold changes per month. (**b**) Scatterplot of sex-independent chromatin accessibility by ATAC-seq changes, reported as DESeq2 log_2_ fold changes per month. (**c**) Scatterplot of sex-independent H3K4me3 CUT&RUN signal changes, reported as DESeq2 log_2_ fold changes per month. (**d-f**) Circular genome plot of the genomic positions of peaks with significant aging regulation in females and males in RNA-seq (**d**), ATAC-seq (**e**), and H3K4me3 CUT&RUN (**f**) (FDR < 5%).

**Extended Data Figure 6. Analysis of X-linked genes.**

(**a**) Boxplot showing *Xist* expression using bulk RNA-seq DESeq2 VST-normalized log_2_ counts with DESeq2 FDR. Significance as FDR from DESeq2. (**b**) Violin plot of *Xist* expression using scRNA-seq using 10xGenomics NIA v2, NIA v3, and JAX datasets. Significance in two-sided Wilcoxon rank-sum tests. (**c**) GSEA of X chromosome inactivation (XCI) and X-linked genes in the aging female peritoneal macrophage transcriptome.

**Extended Data Figure 7. Gene set enrichment analysis of sex-independent and sex-dimorphic aging signatures across transcriptomic and epigenomic landscapes.**

(**a**, **b**, **c**) Top Gene Ontology GSEA terms for sex-independent aging from RNA-seq (**a**), ATAC-seq (**b**), and H3K4me3 (**c**). (**d**, **e**, **f**) Top Reactome GSEA terms for sex-independent aging from RNA-seq (**d**), ATAC-seq (**e**), and H3K4me3 CUT&RUN (**f**). (**g**, **h**, **i**) Top Reactome GSEA terms for sex-dimorphic aging from RNA-seq (**g**), ATAC-seq (**h**), and H3K4me3 CUT&RUN (**i**).

**Extended Data Figure 8. Analysis of peritoneal macrophage and BMDM phagocytosis as a function of aging and sex.**

(**a**) Representative flow cytometry gating strategy for MACS-purified peritoneal macrophage phagocytosis in young female mice. The macrophage population is defined as F4/80^+^ cells, and the phagocytosing population are identified by Alexa Fluor 488-Zymosan bioparticle signals. (**b**) Quantification of bone marrow-derived macrophage (BMDM) phagocytosis using microscopy; n = 29 per sex and age (6 independent cohorts). (**c**) Representative flow cytometry gating strategy for MACS-purified small and large peritoneal macrophage phagocytosis in young female mice. SPM are defined as CD11b^+^F4/80^+Low^, while LPM are identified as CD11b^+^F4/80^+High^, and the fraction of phagocytosing cells in each population is identified by the Alexa Fluor 488-Zymosan bioparticle signal. (**d**) Quantification of MACS-purified small and large peritoneal macrophage phagocytosis using flow cytometry. (**e**) DESeq2 VST-normalized log_2_ count expression of zymosan pattern recognition receptors (PRRs) *Clec7a* (Dectin-1) (left panel) and *Tlr2* (right panel) in peritoneal macrophages using bulk RNA-seq. Significance as FDR from DESeq2. (**f**) Representative flow cytometry gating strategy for surface marker median fluorescence intensity (MFI). The macrophage population is defined as F4/80^+^ cells. Zymosan PRR-expressing macrophages are identified as Dectin-1^+^ or Tlr2^+^ cells. (**g**) Boxplot of protein expression levels quantification on Dectin-1 and Tlr2 in peritoneal macrophages (n = 20 for young males, n = 19 for young females, n = 13 for old females, and n = 16 for old males; 3 independent cohorts). In panels (**b**, **d**, **e**, **g**), circles/squares represent NIA/JAX mice, respectively. Significance in non-parametric two-sided Wilcoxon rank-sum tests are reported in (**b**, **d**, **g**). The center line represents the sample median, the box limits consist of the 25^th^ and 75^th^ percentiles, and the whiskers span 1.5x the interquartile range.

**Extended Data Figure 9. Validation of shRNA-mediated knock-down for candidate phagocytosis regulators genes in RAW264.7 cells.**

(**a**) RT-qPCR validation of shRNA-mediated gene knockdown for *B4galt*, *Dpy19l3*, *Man1a2*, *Rps6ka*, *Tbl1xr1*, *Tm9sf3*, *Gna13*, *Hmgcs1*, *and Magt1*(n = 6) in RAW264.7 cells. Significance in non-parametric one-sided Wilcoxon rank-sum tests.

**Extended Data Figure 10. BMDM metabolism analysis using the Agilent Seahorse platform.**

(**a**) Representative normalized oxygen consumption rate curve (pmol/min/DNA) using the Agilent Seahorse platform.(**b**) Normalized respiration quantification (n = 15 for young old females and males). BMDMs were differentiated from 3 independent cohorts of C57BL/6JNia mice. Significance in non-parametric two-sided Wilcoxon rank-sum tests are reported.

**Extended Data Figure 11. Macrophage polarization analysis.**

(**a**) GSEA of M1/M2 polarization genes, vs. our foundational aging female and male peritoneal macrophage transcriptome dataset. NES: Normalized Enrichment Score. FDR: False Discovery Rate. (**b**) Representative flow cytometry gating strategy for surface marker median fluorescence intensity (MFI). The macrophage population is defined as F4/80^+^ cells. M1 macrophages are characterized by CD80^+^ and CD86^+^ expression, while M2 macrophages are defined as CD163^+^ and CD206^+^. (**c)** Boxplot of protein expression quantification for pro-inflammatory cell surface markers [CD80 (left panel); CD86 (right panel)]; n = 15 for young females, n = 12 for old females n = 13 for young males, and n = 14 for old males. MACS-purified peritoneal macrophages were obtained from 3 independent cohorts. (**d**) Boxplot of protein expression quantification for anti-inflammatory cell surface markers [CD163 (left panel); CD206 (right panel)]; n = 15 for young females, n = 12 for old females, n = 13 for young males, and n = 14 for old males; MACS-purified peritoneal macrophages were obtained from 3 independent cohorts. Significance in non-parametric two-sided Wilcoxon rank-sum tests is reported. (**c-d**) The center line represents the sample median, the box limits consist of the 25^th^ and 75^th^ percentiles, and the whiskers span 1.5x the interquartile range.

**Extended Data Figure 12. Sex steroid hormone receptors expression and effect of estrus cycling on primary macrophage functional phenotypes.**

(**a**) DESeq2-derived FPKM expression of sex hormone receptor genes in peritoneal macrophages bulk RNA-seq (*Esr1*, *Esr2*, *Gper1*, *Pgr*, and *Ar*). **(b)** Bulk RNA-seq analysis of Esr1 transcript expression (DESeq2 VST log_2_ counts) in young female (YF), old female (OF), and young male (YM) or old male (OM) peritoneal macrophages. Significance as FDR from DESeq2. (**c**) Experimental scheme illustrating the mouse estrus cycle, including predicted relative estrogen concentration in blood and representative vaginal cytology images. (**d**) Boxplot of phagocytosis quantification via microscopy in 4-month-old female mice at proestrus (n = 28), estrus (n = 20), metestrus (n = 9), and diestrus (n = 7), alongside male control mice (n = 19). Data from 4 independent cohorts of C57BL/6NTac. (**e**) Representative normalized oxygen consumption rate (OCR; Top plot) and extracellular acidification rate (ECAR; bottom plot) (pmol/min/DNA) measured using Seahorse assays in estrus cycle stages and male mice. (**f, g**) Quantification of metabolic parameters from Seahorse extracellular flux analysis. Normalized glycolysis (**g**) and respiration (**f**) of BMDMs from various hormone conditions and sexes (high E2 females, low E2 females, and males) [proestrus (n = 28), estrus (n = 20), metestrus (n = 9), and diestrus female mice (n = 7), alongside male mice (n = 19)]. BMDMs were obtained from 3 independent cohorts. Triangles/squares for Taconic/JAX. (**h**) Macrophage purity in Sham (n = 17) Vs. OVX (n = 17) mice. (**d, f, g, h**) Significance in non-parametric two-sided Wilcoxon rank-sum tests is reported. (**b, d, f, g**,**h)** The center line represents the sample median, the box limits consist of the 25^th^ and 75^th^ percentiles, and the whiskers span 1.5x the interquartile range.

**Extended Data Figure 13. Analysis of macrophages from short-term late-life E2 supplementation (STE2) and long-term periestropausal E2 supplementation (LTE2).**

(**a**) Experimental scheme describing short-term late-life 17β-estradiol (E2) supplementation in female mice. **(b)** Purity of MACS-purified peritoneal macrophages from young vehicle female (n = 8), old vehicle female (n = 9), and E2-supplemented old female mice (n = 9). Cells obtained from 2 independent cohorts. Significance in non-parametric two-sided Wilcoxon rank sum test. (**c**) MDS analysis of gene expression profiles in young females, old females, and STE2-treated old females; n = 5 for young females, n = 5 for old females, n = 4 for STE2-treated old females. (**d**) Scatterplot showing the relationship between log₂ fold changes in gene expression with aging in macrophages from vehicle treated females and log₂ fold changes in gene expression in macrophages from old STE2 vs. vehicle treated females. The Spearman rank correlation Rho value, as well as the significance of the correlation are reported. (**e**) GSEA of STE2-regulated genes in old female macrophages, *vs*. the foundational aging female peritoneal macrophage transcriptomic dataset. (**f**) Boxplot of phagocytosis of peritoneal macrophages in young females (n = 13), old females (n = 13), and STE2-treated old females (n = 14; animals from 3 independent cohorts). (**g**) Boxplot of glycolysis quantification of peritoneal macrophages in young females, old females, and STE2-treated old females (n = 14 for young females, old females, and E2-treated old females; animals from 3 independent cohorts). (**h**) Boxplot of LTE2 macrophage purity. (**i**) Scatterplot showing the relationship between log₂ fold changes in gene expression with aging in macrophages from vehicle-treated females and log₂ fold changes in gene expression in macrophages from old LTE2 vs. vehicle treated females. (**b, c, d, h, i**) Significance in Kruskal-Dunn test are reported. The center line represents the sample median, the box limits consist of the 25^th^ and 75^th^ percentiles, and the whiskers span 1.5x the interquartile range.

**Extended Data Figure 14. Analysis of female J774A.1 immortalized macrophages treated with 100nM E2 *in vitro*.**

(**a**) Experimental scheme for cell autonomous response to 100nM E2 treatment using female J774A.1 immortalized macrophages. (**b**) MDS analysis of gene expression profiles in vehicle and 100nM E2 treated J774A.1 (n = 3 for vehicle, n = 4 for 100nM E2 treated). (**c**) GSEA of E2-regulated genes in J774A.1, *vs*. the foundational aging female peritoneal macrophage transcriptomic dataset. (**d**) Boxplot of phagocytosis of vehicle and 100nM E2 treated J774A.1 (n = 9 for vehicle, n = 9 for 100nM E2 treated; from 3 independent experiments). Significance in non-parametric two-sided Wilcoxon rank-sum tests. The center line represents the sample median, the box limits consist of the 25^th^ and 75^th^ percentiles, and whiskers span 1.5x the interquartile range.

**Extended Data Figure 15. Analysis of macrophages from young adult Esr1 knock-out (KO) female mice.**

(**a**) Experimental scheme for *Esr1* knockout (KO) mice. **(b)** Purity of MACS-purified peritoneal macrophages from WT and Esr1 KO mice (n = 15). Purified peritoneal macrophages were obtained from 4 independent cohorts of mice. **(c)** Bulk RNA-seq analysis of Esr1 expression (DESeq2 VST-normalized log_2_ counts) in peritoneal macrophages from WT and Esr1 KO mice (n = 6). Significance as FDR from DESeq2. (**d**) MDS analysis of gene expression profiles in WT and *Esr1* KO mice (n = 5 for WT, n = 6 for *Esr1* KO). (**e**) GSEA of *Esr1* KO-regulated genes, *vs*. the foundational aging female peritoneal macrophage transcriptomic dataset. (**f**) Boxplot of phagocytosis of peritoneal macrophages in WT and *Esr1* KO mice (n = 15 for WT, n = 13 for *Esr1* KO; animals from 4 independent cohorts). (**g**) Boxplot of glycolysis quantification of peritoneal macrophages in WT and *Esr1* KO mice (n = 15 for WT and *Esr1* KO; animals from 4 independent cohorts). (**b**, **c**, **f, g**) Significance in non-parametric two-sided Wilcoxon rank-sum tests were reported. The center line represents the sample median, the box limits consist of the 25^th^ and 75^th^ percentiles, and the whiskers span 1.5x the interquartile range.

**Extended Data Figure 16. Expression and activity of candidate TF genes.**

(**a**) DESeq2 VST-normalized log_2_ count gene expression of *Irf2*, *Mef2c*, *Meis1*, *Tal1*, and *Tbl1xr1* (Bulk RNA-seq). Significance as DESeq2 FDR. The center line represents the sample median, the box limits consist of the 25^th^ and 75^th^ percentiles, the whiskers span 1.5x the interquartile range. (**b**) Top 15 most significantly variable footprint scores across samples according to ChromVAR analysis of ATAC-seq footprint using HOMER motifs. Left: heatmap of median footprint accessibility score values in each biological group. Right: barplot of -log_10_(FDR) for significance of variability in footprint accessibility according to ChromVAR (**Extended Data Table 6A**). (**c, d**) Dotplot (**c**) and violin plot (**d**) of predicted TF activity in peritoneal macrophages from scRNA-seq using SCENIC (**Extended Data Table 6B**).

**Extended Data Figure 17. Impact of shRNA-mediated knock-down of candidate TFs in RAW264.7 macrophages.**

(**a**) RT-qPCR validation of gene knockdown using shRNA targeting different TFs (n = 6 infections per hairpin) in RAW264.7 macrophages. Significance in non-parametric one-sided Wilcoxon rank-sum tests (Hypothesis: gene expression is decreased upon knock-down). (**b**) DESeq2 VST-normalized log_2_ count gene expression of TFs after gene knockdown using shRNA (Bulk RNA-seq). Goldenrod (shLuciferase; n = 3), light goldenrod (non-targeting control; n = 3), dark green (sh1; n = 3), and light green (sh2; n = 3). (**c**) Multidimensional scaling (MDS) plot of transcriptomic profiles. Goldenrod (shLuciferase; n = 3), light goldenrod (non-targeting control; n = 3), dark green (sh1; n = 3), and light green (sh2; n = 3) indicate distinct transcriptomic patterns based on gene expression knockdown. (**d**) Heatmap of differential gene expression after TF knockdown (FDR < 10^-4^). (**e**) Heatmap of TFs expression upon perturbation based on DESeq2 VST-normalized log_2_ counts (row scaled). **(f)** RT-qPCR analysis showing Mef2c knockdown (*shMef2c*) significantly reduces Irf2 expression in RAW264.7 macrophages. Relative gene expression is normalized to housekeeping control (n = 3 infections per hairpin). Significance in non-parametric two-sided Wilcoxon rank-sum tests. **(g)** DESeq2 VST-normalized log_2_ count gene expression for Rps6ka3 in RAW264.7 macrophages RNA-seq upon Meis1 knockdown. (**b**, **g**) Significance as FDR from DESeq2. For boxplots in panels (**a, b, f, g**), the center line represents the sample median, the box limits consist of the 25^th^ and 75^th^ percentiles, the whiskers span 1.5x the interquartile range.

**Extended Data Figure 18. Regulation of Irf2 expression with aging and in response to changes in ovarian hormone signaling.**

(**a**) Boxplot of *Irf2* gene expression in mouse peritoneal macrophages using RT-qPCR (young female n = 12, old female n = 8, young male n = 12, and old male n = 9; animals from 4 independent cohorts). Significance in non-parametric two-sided Wilcoxon rank-sum tests. (**b**) Representative flow cytometry gating strategy for intracellular Irf2 detection at the protein level using PE-conjugated antibody for Irf2 and isotype control. (**c**) Comparison of Irf2 and isotype control median fluorescence intensity in peritoneal macrophages from young and old mice, stratified by sex. (**d**) Boxplots of *Irf2* gene expression as DESeq2 VST-normalized log_2_ count in peritoneal macrophages from OVX vs. sham, STE2-treated, LTE2-treated, and *Esr1* KO peritoneal macrophages. (**e**) DESeq2 VST-normalized log_2_ count gene expression for Irf2 in J774A.1 immortalized macrophages treated with 100nM E2 vs. vehicle. (**d, e**) Significance as DESeq2 FDR.

**Extended Data Figure 19. Validation of Irf2 shRNA-mediated knock-down phenotypes in female J774A.1 immortalized macrophages.**

(**a**) Experimental design for lentiviral knockdown of *Irf2* in J774A.1 cells. (**b**) Irf2 expression in J774A.1 cells after *Irf2* knockdown using shRNA by RT-qPCR (n = 6 infections per hairpin). Significance in non-parametric one-sided Wilcoxon rank-sum tests compared to control non targeting and luciferase hairpins. (**c**) Irf2 expression in J774A.1 cells after *Irf2* knockdown using shRNA by RNA-seq expressed as DESeq2 VST-normalized log_2_ count (n = 3 infections per hairpin). Significance as DESeq2 FDR. (**d**) Intracellular flow cytometry quantification of normalized Irf2 protein expression in RAW264.7 (left panel; circle; n = 6 infections per hairpin) and J774A.1 (right panel; square; n = 6 infections per hairpin) macrophages transduced with two independent non-targeting control hairpins [shLuciferase (goldenrod), non-targeting control (light goldenrod)] or Irf2-targeting shRNAs hairpins [sh1 (dark green), or sh2 (light green)]. Significance in non-parametric one-sided Wilcoxon rank-sum (Hypothesis: protein expression is decreased with knock-down). (**e**) Multidimensional scaling (MDS) plot of Irf2-knockdown and control J774A.1 cells. shLuciferase (goldenrod), non-targeting control (light goldenrod), dark green (sh1), and light green (sh2); n = 3 infections per hairpin. (**f**) Venn diagrams showing the overlap of significantly upregulated (left) and downregulated (right) genes (FDR < 1%) following Irf2 knockdown in RAW264.7 and J774A.1 macrophages. Significance of overlap in Fisher’s exact test. (**g**) Phagocytosis quantification Irf2 knockdown in RAW264.7 (data repeated from **Figure 6h** for visual convenience) and J774A.1 cells compared to control, using flow cytometry. Circles/squares for RAW264.7/J774A.1 samples; dark green (sh1) and light green (sh2); n = 6 infections per hairpin. Significance in non-parametric one-sided Wilcoxon rank-sum tests were reported (hypothesis: same direction as aging). (**b**, **c**, **d, g**) The center line represents the sample median, the box limits consist of the 25^th^ and 75^th^ percentiles, and the whiskers span 1.5x the interquartile range.

**Extended Data Figure 20. Quality control analysis for Irf2 genomic profiling experiments.**

(**a**) Top 5 enriched transcription factor binding motifs found at Irf2 ChIP-seq peaks in BMDMs (left) and CUT&RUN peaks in peritoneal macrophages (right). (**b**) Circular genome plot of the genomic positions of Irf2-bound peaks in BMDMs (outer circle, light pink) or peritoneal macrophages (inner circle, bright pink). (**c**) Stacked barplots of the genomic distribution of Irf2 ChIP-seq peaks in BMDMs and CUT&RUN peaks in peritoneal macrophages according to ChIPseeker. (**d**) GSEA of genes associated to Irf2 ChIP-seq peaks in BMDMs and/or CUT&RUN peaks in peritoneal macrophages, *vs*. Irf2 shRNA knockdown transcriptomic datasets in RAW264.7 or J774A.1 cells. (**e**) Venn diagrams (left) show the overlap between Irf2 CUT&RUN binding sites in peritoneal macrophages (within 5kb of gene TSS) and genes that are significantly downregulated (left; FDR < 5%) or upregulated (right; FDR < 5%) with aging in female mouse macrophage, based on transcriptomic profiling. (**f**) GSEA plot for genes associated to CUT&RUN peaks with significantly decreased Irf2 binding in aged female peritoneal macrophages, *vs*. the foundational aging female peritoneal macrophage transcriptomic dataset.

**Extended Data Figure 21. Regulation of hexokinase expression levels in mouse macrophages.**

(**a**) DiffBind/DESeq2 spike-in normalized, VST-normalized log_2_ *Irf2* CUT&RUN signal for the peak situated in the Hk3 promoter region (chr13:55022528-55022928, mm10 genome reference). Significance as DESeq2 FDR. (**b-d**) DESeq2 VST-normalized log_2_ count gene expression for Hk3 (**b**), *Hk1* (**c**), and *Hk2* (**d**) in our peritoneal macrophage aging transcriptomic dataset. Significance as DESeq2 FDR. (**e**) Representative flow cytometry gating strategy for intracellular Hk3 detection using Coralite488-conjugated antibody for Hk3 and isotype control. (**f**) Comparison of Hk3 and isotype control median fluorescence intensity in peritoneal macrophages from young and old female mice. (**g-h**) DESeq2 VST-normalized log_2_ count gene expression for *Hk1* (**g**), and *Hk2* (**h**) in our Irf2 shRNA knockdown transcriptomic datasets in RAW264.7 or J774A.1 cells. Significance as DESeq2 FDR.

**Inventory of Extended Data Tables**

**Extended Data Table 1. Resource table mouse information shRNA qPCR primers.**

(**A**) Animal metadata for aging C57BL/6JNia and C57BL/6J used in this study. (**B**) Animal metadata for C57BL/6NTac estrus cycle mice used in this study. (**C**) Vaginal cytology estrus cycle phase calling data used in this study. (**D**) Animal metadata for C57BL6/J ovariectomy (OVX) vs. sham mice used in this study. (**E**) Animal metadata for short-term late-life C57BL6/J 17b-estradiol supplementation [STE2] mice used in this study. (**F**) Animal metadata for long-term peri-estropausal C57BL6/J 17b-estradiol supplementation [LTE2] mice used in this study. (**G**) Animal metadata for C57BL6/J B6N(Cg)-Esr1^tm4.2Ksk^/J (Esr1 KO) mice used in this study. (**H**) List of control plasmids and packaging plasmids. List of MISSION shRNA for phagocytosis regulators and transcription factor candidates. (**I**) List of qPCR primers used for RT-qPCR.

**Extended Data Table 2. Differential expression/accessibility/modification analysis of genes/peaks regulated with aging in males vs. females.**

(**A-F**) MetaRNAseq results for pseubulk-level differential expression analysis with aging in male vs. female cells: (A) female B- cells, (B) male B-cells, (C) female macrophages, (D) male macrophages, (E) female T0-cells, (F) male T-cells. (**G**) DEseq2 analysis of gene regulation with aging in male vs female peritoneal macrophages by RNA-seq (Merged Table). (**H**) DEseq2 analysis of peak accessibility remodeling with aging in male vs female peritoneal macrophages by ATAC-seq (Merged Table). (**I**) DEseq2 analysis of peak CUT&RUN signal remodeling with aging in male vs female peritoneal macrophages by H3K4me3 (Merged Table).

**Extended Data Table 3. Functional enrichment analysis aging and sex.**

(**A**) RNA-seq GSEA Gene Ontology (FDR < 5%) for differential regulation with respect to age and sex. (**B**) RNA-seq GSEA Reactome (FDR < 5%) for differential regulation with respect to age and sex. (**C**) ATAC-seq GSEA Gene Ontology (FDR < 5%) for differential regulation with respect to age and sex. (**D**) ATAC-seq GSEA Reactome (FDR < 5%) for differential regulation with respect to age and sex. (**E**) H3K4me3 CUT&RUN GSEA Gene Ontology (FDR < 5%) for differential regulation with respect to age and sex. (**F**) H3K4me3 CUT&RUN GSEA Reactome (FDR < 5%) for differential regulation with respect to age and sex. (**G**) GSEA Macrophage CRISPR phagocytosis screens female and male analysis. (**H**) GSEA Macrophage MsigDB hallmark genes related to metabolism female and male analysis. (**I**) GSEA Macrophage polarization markers female and male analysis.

**Extended Data Table 4. List of curated phagocytosis regulators and polarization genes.**

(**A**) List of phagocytosis regulators from CRISPR/Cas9 phagocytosis screens: Haney et al. 2018, Nature genetics ^1^ and Pluvinage et al., 2019 Nature ^2^ (**B**) List of curated genes involved in M1-like and M2-like polarization phenotypes based on a publicly available bone marrow-derived macrophage RNA-seq dataset ^3^ (GSE103958).

**Extended Data Table 5. Ovarian hormone signaling differential regulated genes in macrophages.**

(A) DEseq2 differential analysis of genes between ovariectomy (OVX) vs. sham peritoneal macrophages by RNA-seq (All genes). (B) DEseq2 differential analysis of genes between vehicle-treated young vs short-term vehicle-treated old peritoneal macrophages by RNA-seq (All genes; *in vivo* STE2 experiment). (C) DEseq2 differential analysis of genes between short-term old vehicle control vs. short-term E2 supplemented peritoneal macrophages by RNA-seq (All genes; *in vivo* STE2 experiment). (D) DEseq2 differential analysis of genes between vehicle-treated young vs long-term vehicle-treated old peritoneal macrophages by RNA-seq (All genes; *in vivo* LTE2 experiment). (E) DEseq2 differential analysis of genes between long-term old vehicle control vs. long-term treated E2 supplemented peritoneal macrophages by RNA-seq (All genes; *in vivo* LTE2 experiment). (F) DEseq2 differential analysis of genes between vehicle control vs. E2 treated J774A.1 macrophages by RNA-seq (All genes; *in vitro* supplementation experiment). (G) DEseq2 differential analysis of genes between female *Esr1* knockout and wildtype peritoneal macrophages by RNA-seq (All genes).

**Extended Data Table 6. Transcription factor candidate knockdown genes.**

(**A**) ChromVar HOMER TF footprint accessibility variability analysis in aging male and female macrophages (ATAC-seq) (**B**) SCENIC TF regulon activity impacted by aging in female or male macrophages (scRNA-seq) (Wilcoxon rank-sum test) (**C**) DEseq2 differential gene regulation analysis between *Irf2* shRNA knockdown and non-targeting control in RAW264.7 by RNA-seq. (**D**) DEseq2 differential gene regulation analysis between *Mef2c* shRNA knockdown and non-targeting control in RAW264.7 by RNA-seq. (**E**) DEseq2 differential gene regulation analysis between *Meis1* shRNA knockdown and non-targeting control in RAW264.7 by RNA-seq. (**F**) DEseq2 differential gene regulation analysis between *Tal1* shRNA knockdown and non-targeting control in RAW264.7 by RNA-seq. (**G**) DEseq2 differential gene regulation between *Tbl1xr1* shRNA knockdown and non-targeting control in RAW264.7 by RNA-seq. (**H**) DEseq2 differential gene regulation analysis between *Irf2* shRNA knockdown and non-targeting control in macrophage cell line J774A.1 by RNA-seq.

**Extended Data Table 7. Irf2 binding analysis in primary mouse macrophages in young cells and with aging.**

(A) Annotated HOMER/MSPC consensus Irf2 ChIP-seq peaks in young female BMDMs. (B) Annotated HOMER/MSPC consensus Irf2 CUT&RUN peaks in young and old female peritoneal macrophages. (C) HOMER Enriched Irf2 Motifs (Chip-seq; FDR < 5%). (D) HOMER Enriched Irf2 Motifs (CUT&RUN; FDR < 5%). (E) GREAT Gene Ontology analysis for Irf2 gene targets (BMDM ChIP-seq peaks). (F) GREAT Gene Ontology analysis for Irf2 gene targets (Peritoneal macrophage CUT&RUN peaks). (G) ChIPSeeker Gene Ontology analysis for Irf2 gene targets (BMDM ChIP-seq peaks). (H) ChIPSeeker Gene Ontology analysis for Irf2 gene targets (Peritoneal macrophage CUT&RUN peaks). (I) DiffBind/DEseq2 analysis of differential Irf2 CUT&RUN signal in female aging peritoneal macrophages (All peaks).

**Extended Data material references:**

1	Haney, M. S. *et al.* Identification of phagocytosis regulators using magnetic genome-wide CRISPR screens. *Nat Genet* **50**, 1716-1727, doi:10.1038/s41588-018-0254-1 (2018).

2	Pluvinage, J. V. *et al.* CD22 blockade restores homeostatic microglial phagocytosis in ageing brains. *Nature* **568**, 187-192, doi:10.1038/s41586-019-1088-4 (2019).

3	Das, A. *et al.* High-Resolution Mapping and Dynamics of the Transcriptome, Transcription Factors, and Transcription Co-Factor Networks in Classically and Alternatively Activated Macrophages. *Front Immunol* **9**, 22, doi:10.3389/fimmu.2018.00022 (2018).
