## Extended data figures for "Mouse peritoneal macrophages undergo female-specific remodeling with aging driven by both hormone-dependent and -independent mechanisms"

**Extended Data Figure 1**

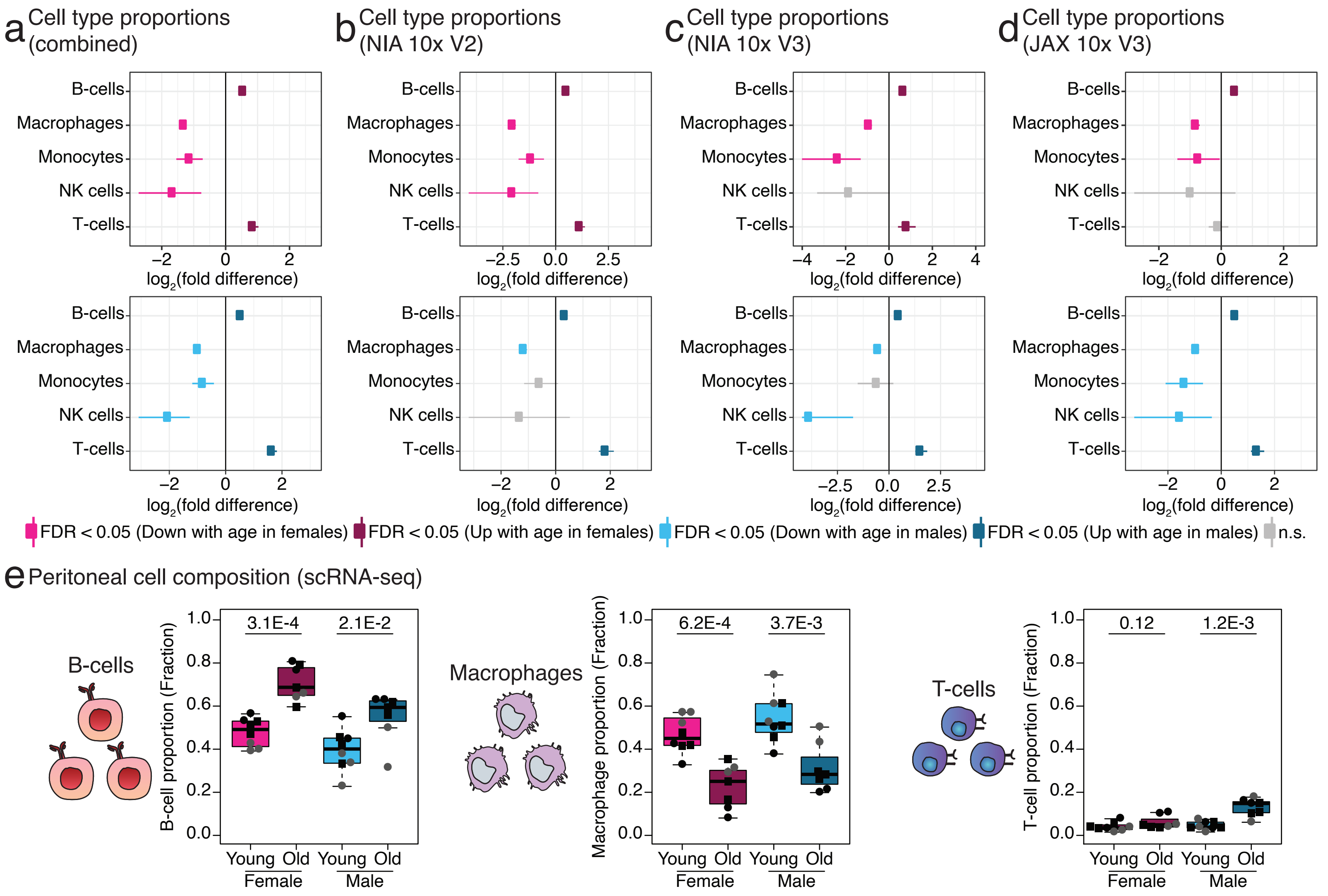

**Extended Data Figure 2**

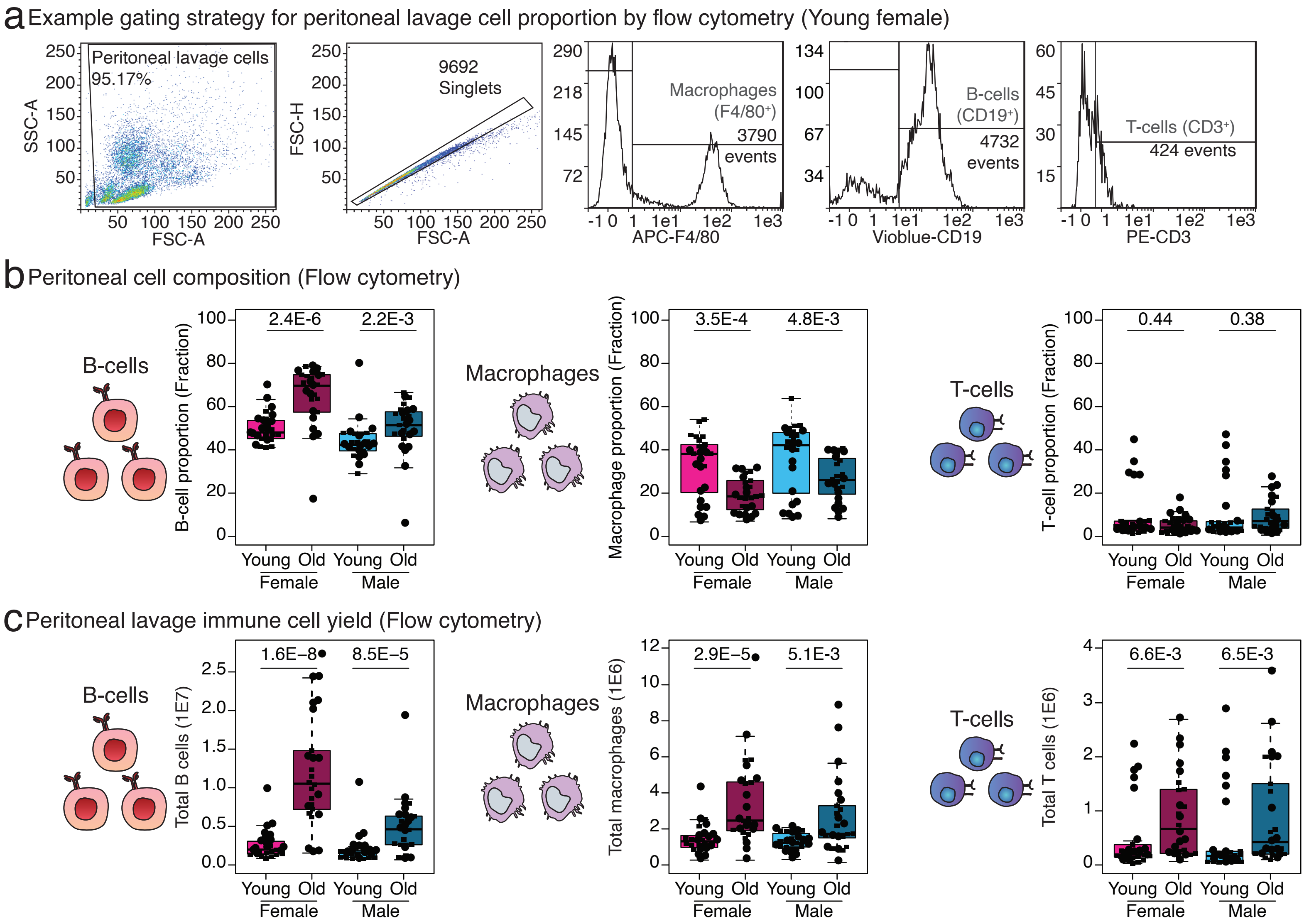

#### Extended Data Figure 3

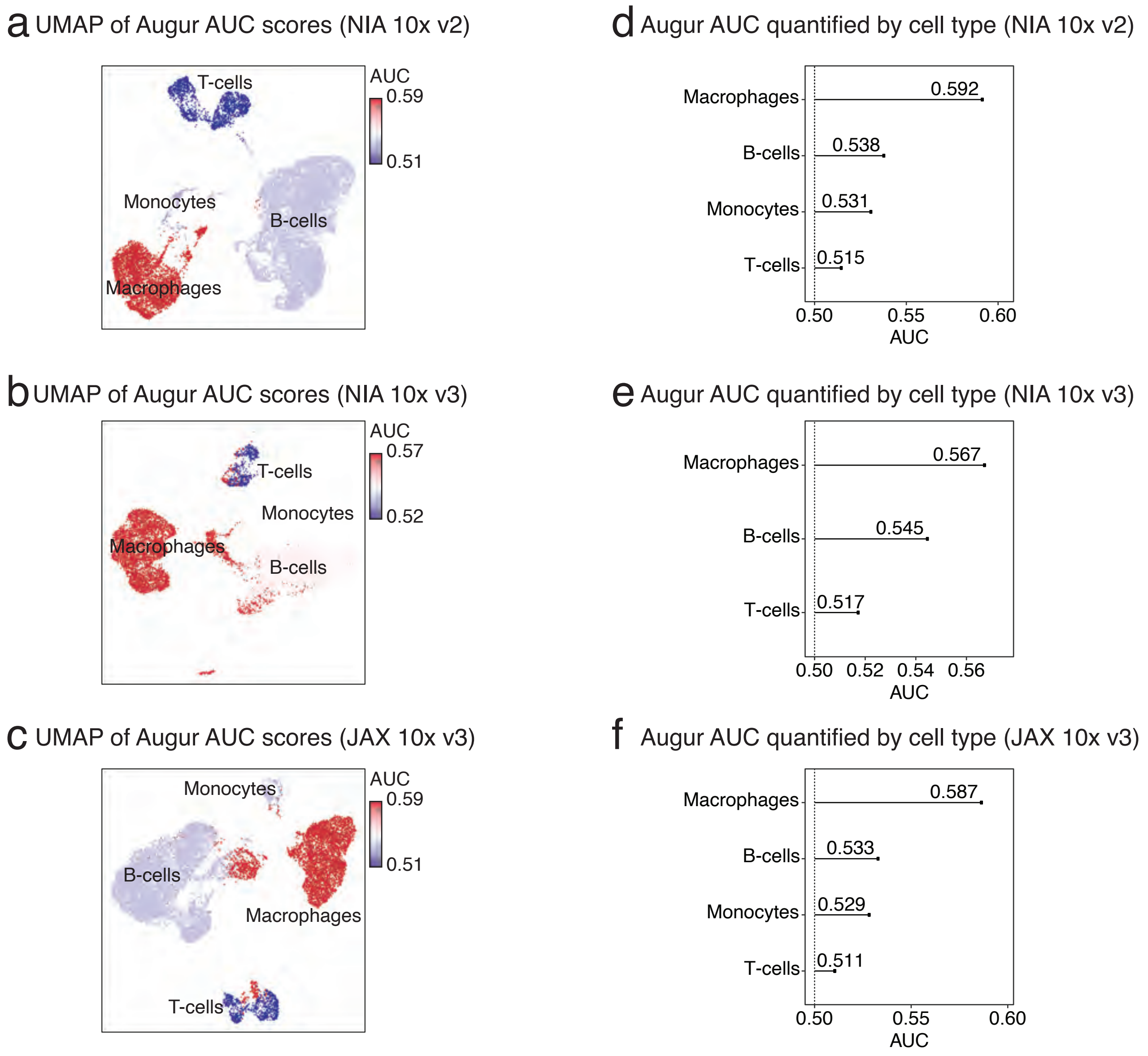

**Extended Data Figure 4**

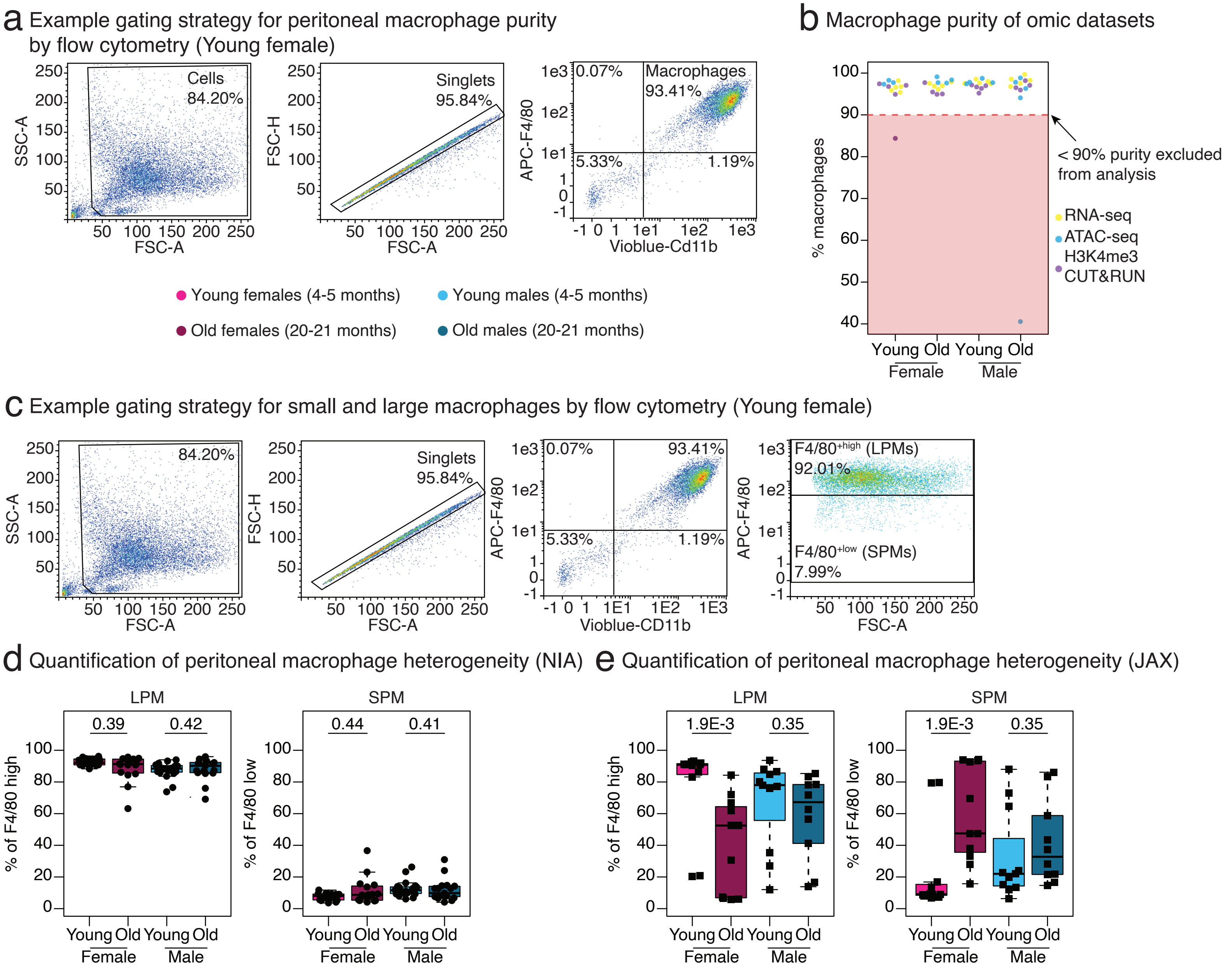

**Extended Data Figure 5**

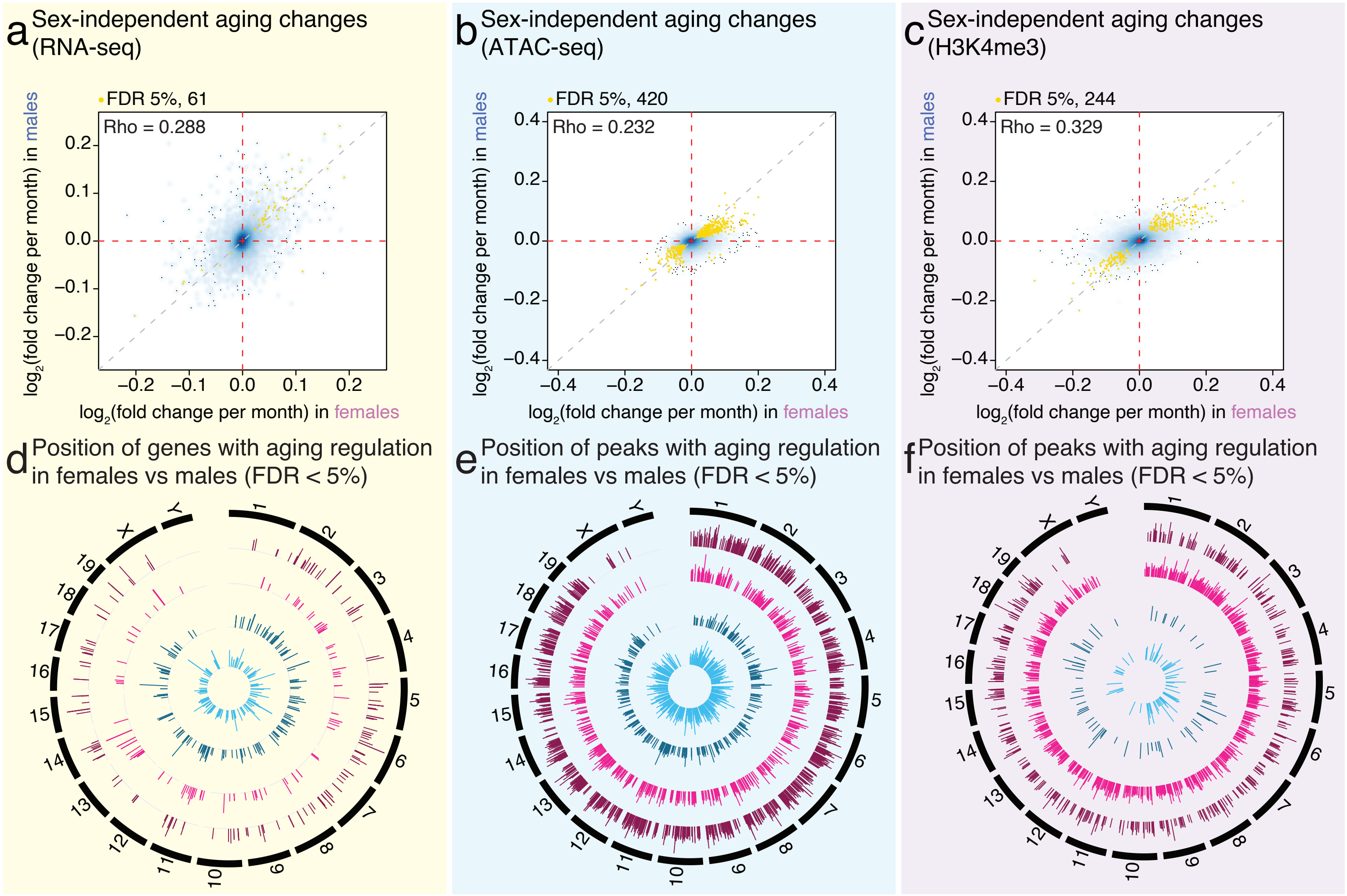

#### Extended Data Figure 6

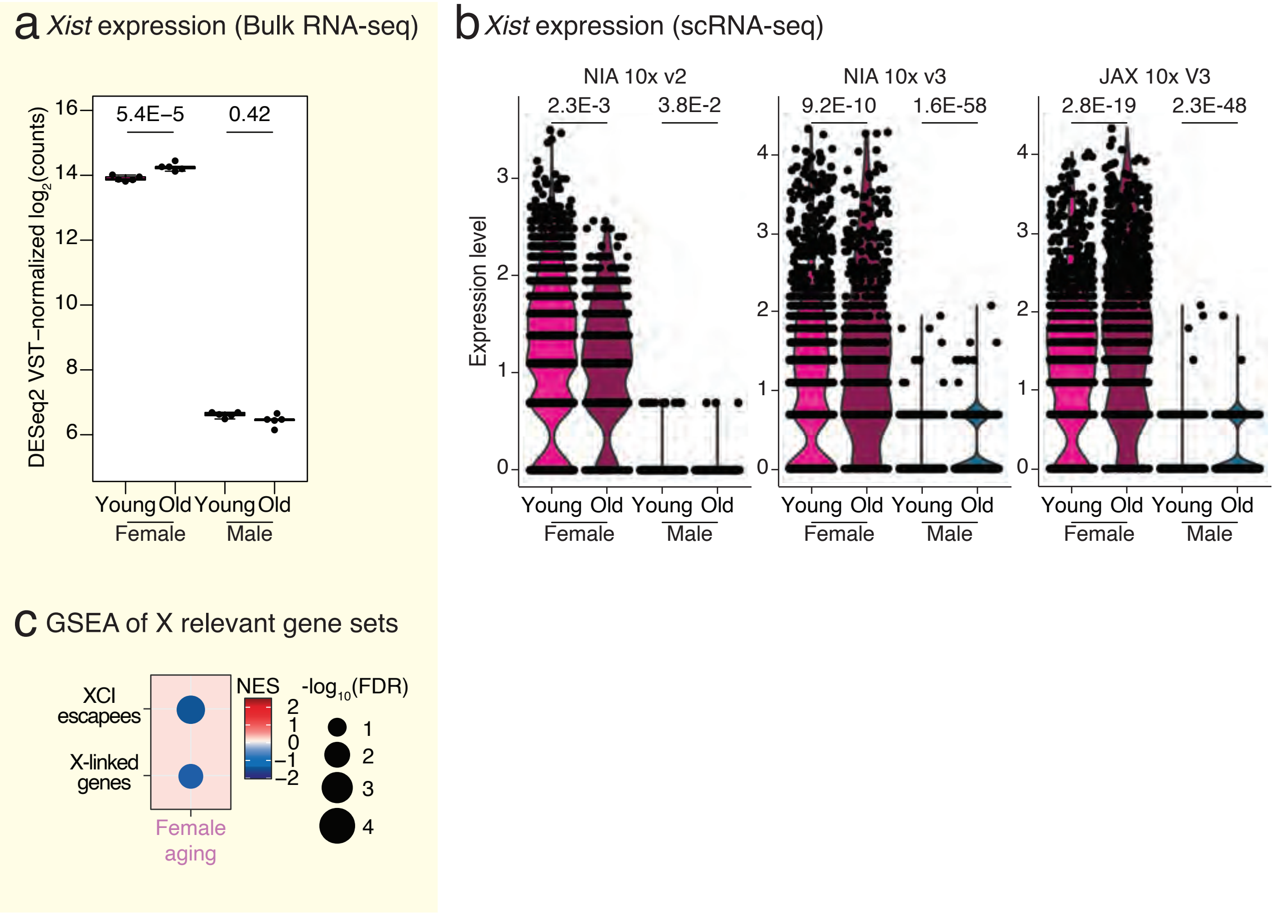

**Extended Data Figure 7**

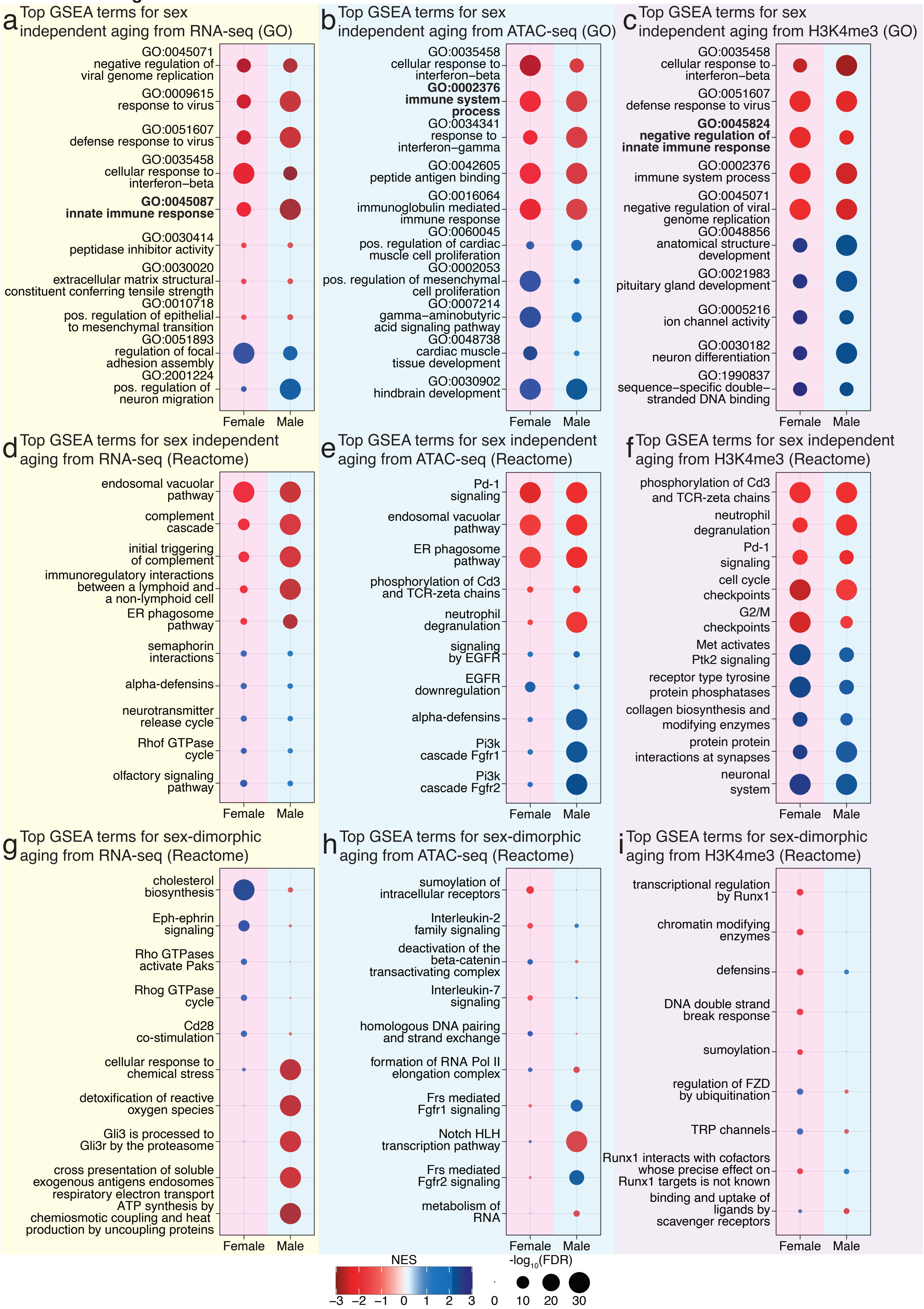

**Extended Data Figure 8**

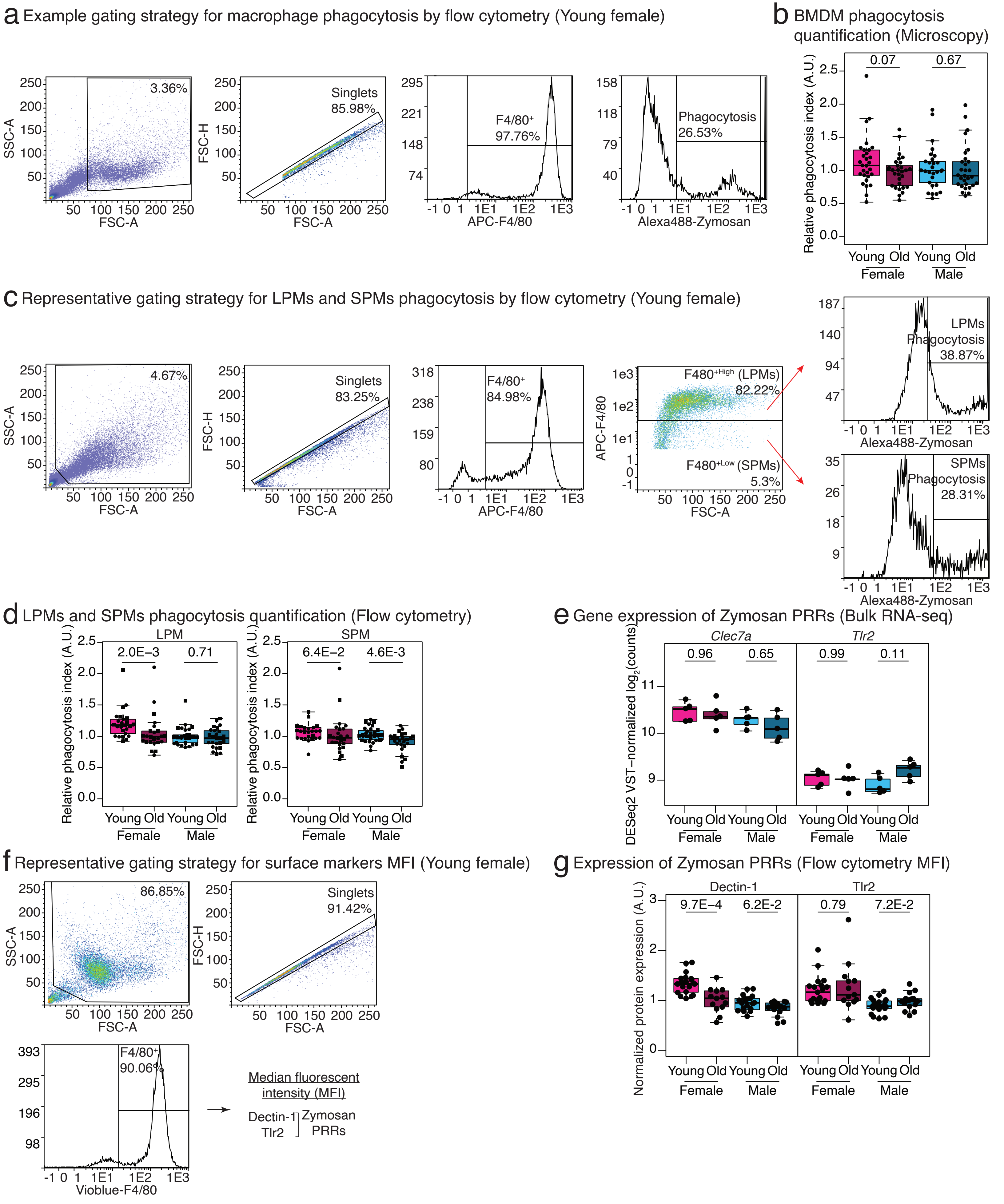

#### Extended Data Figure 9

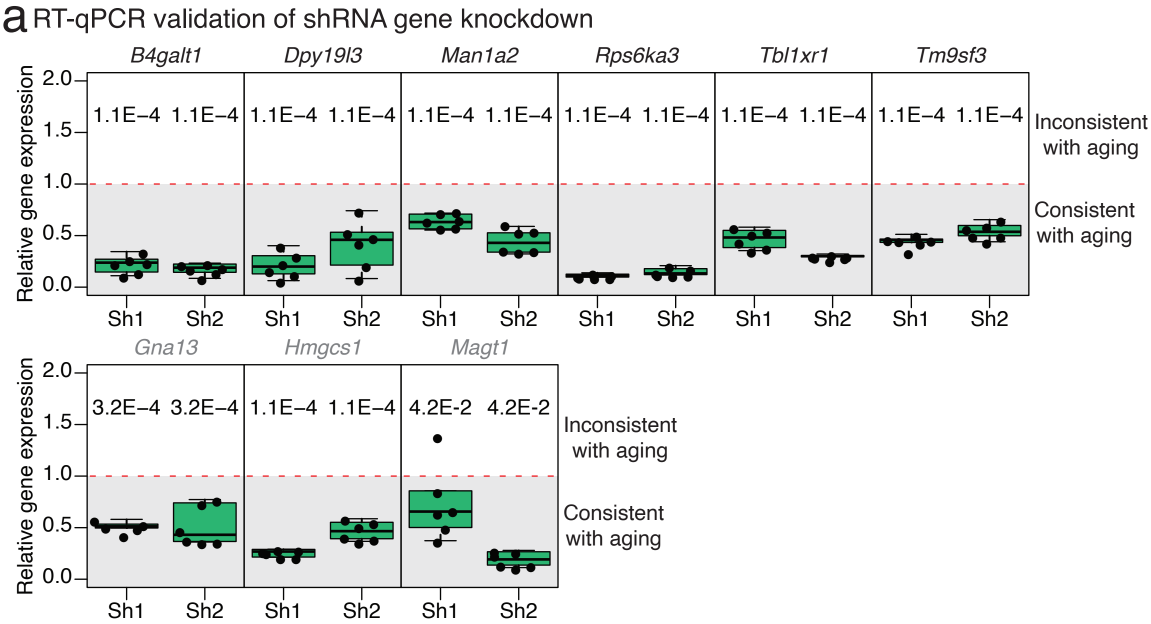

### Extended Data Figure 10

a Representative normalized OCR (Seahorse)

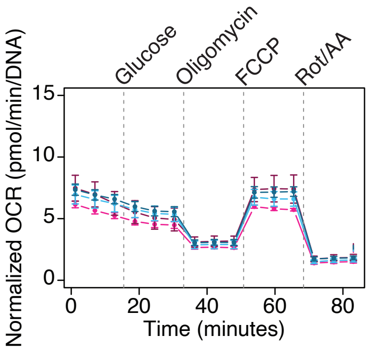


b Respiration quantification (Seahorse)

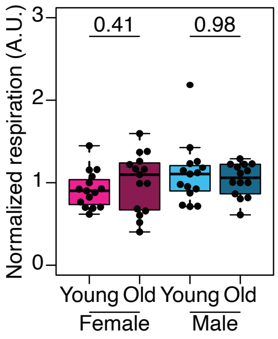

**Extended Data Figure 11**

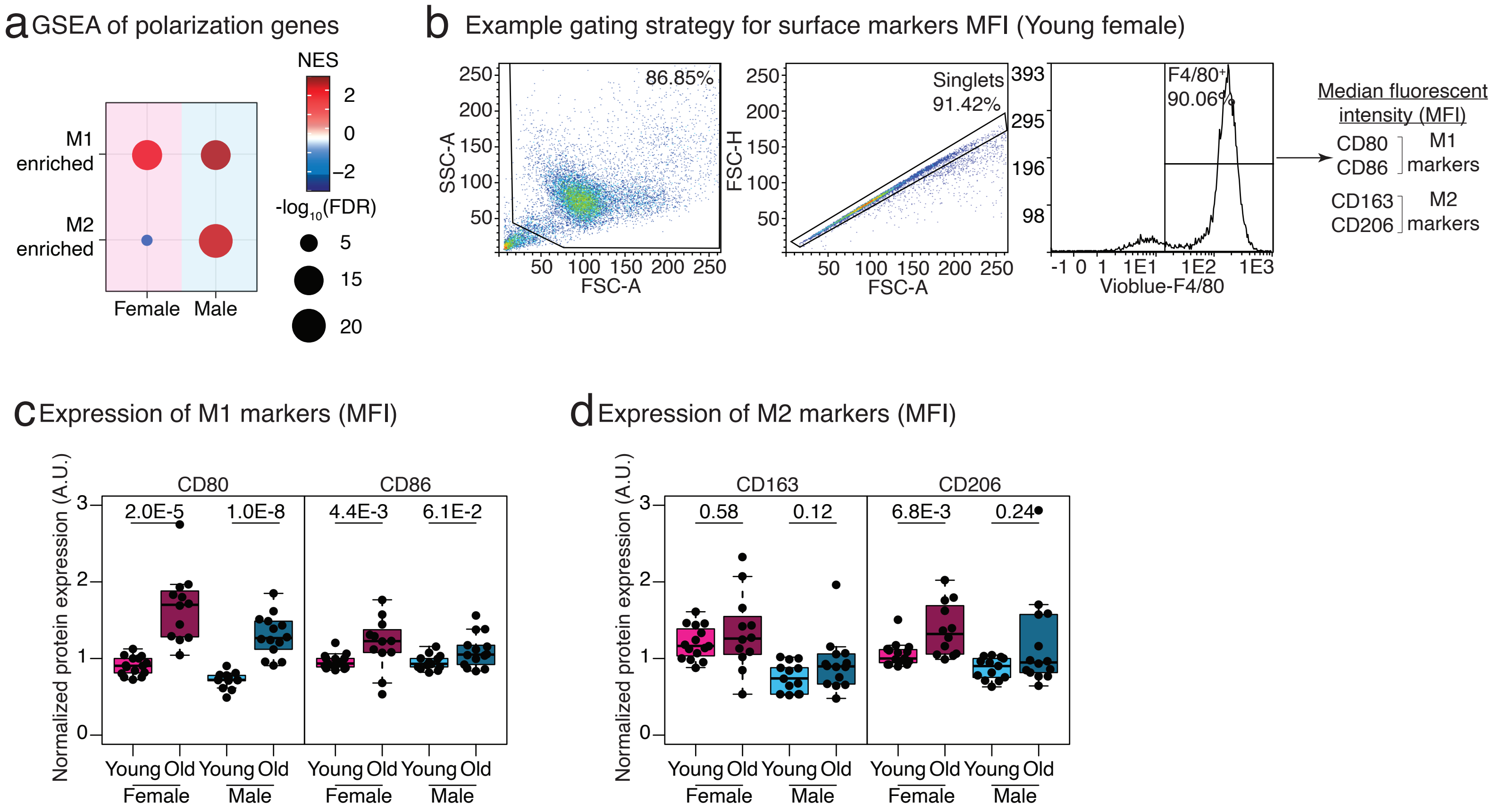

**Extended Data Figure 12**

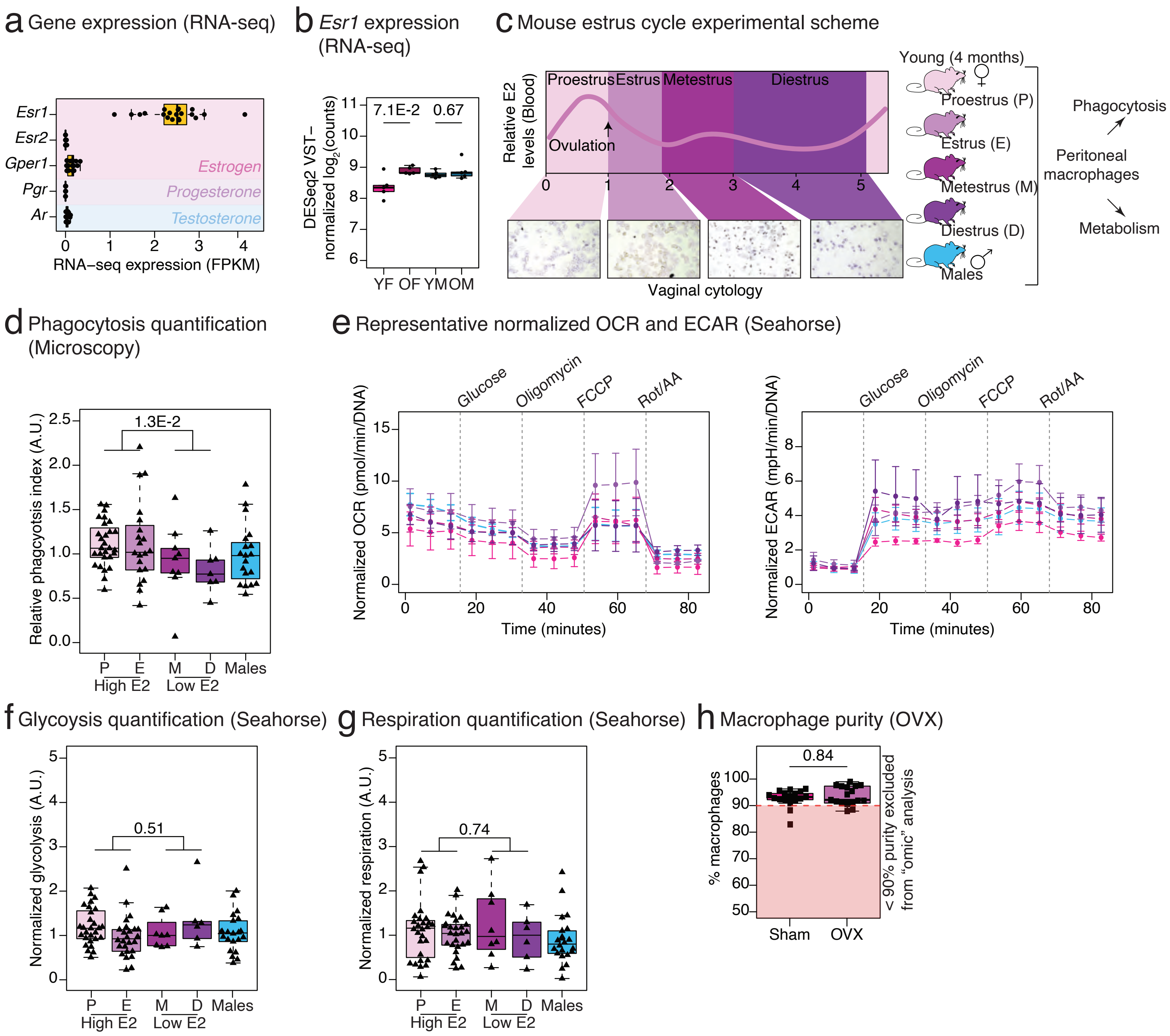

#### Extended Data Figure 13

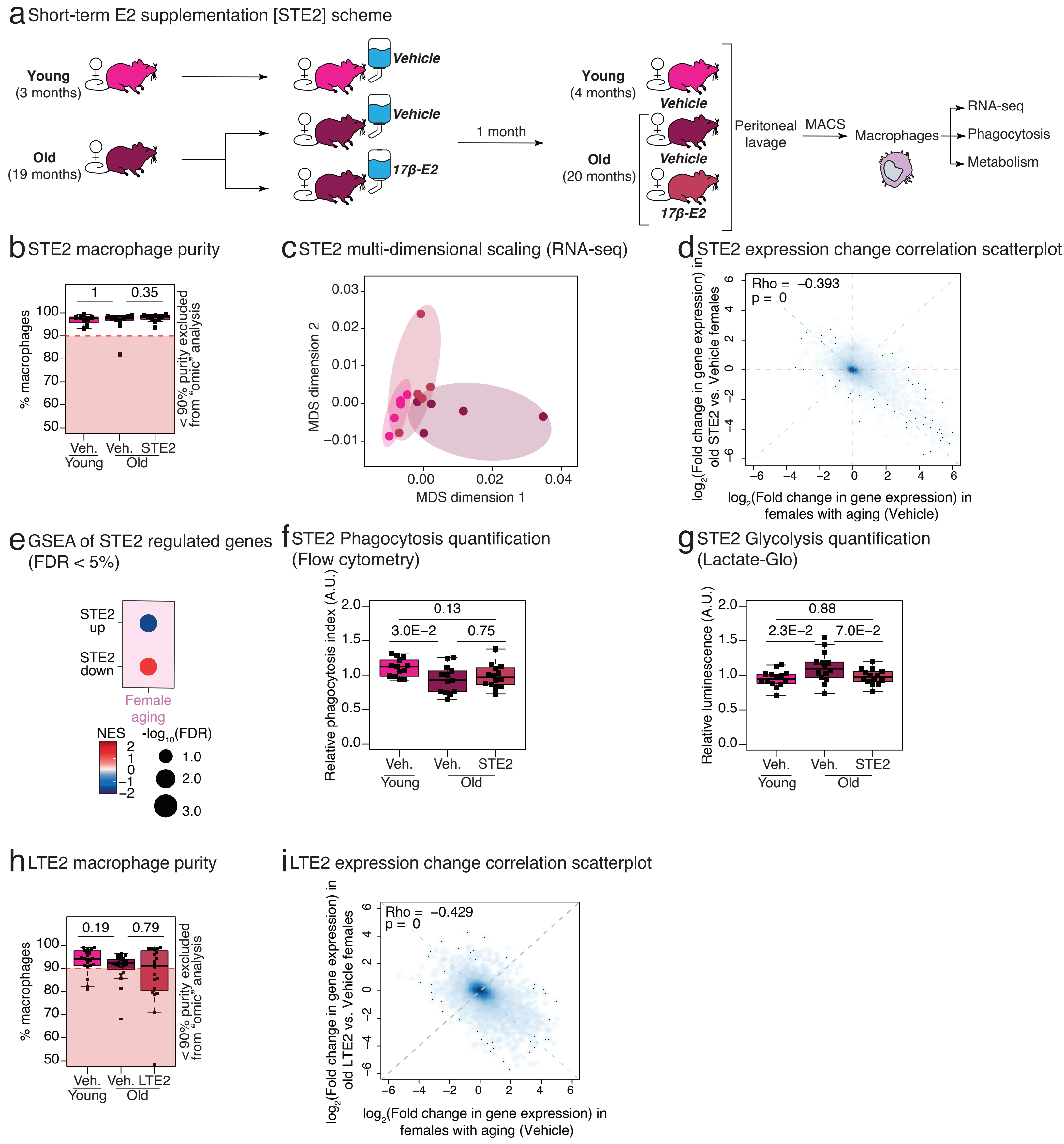

**Extended Data Figure 14**

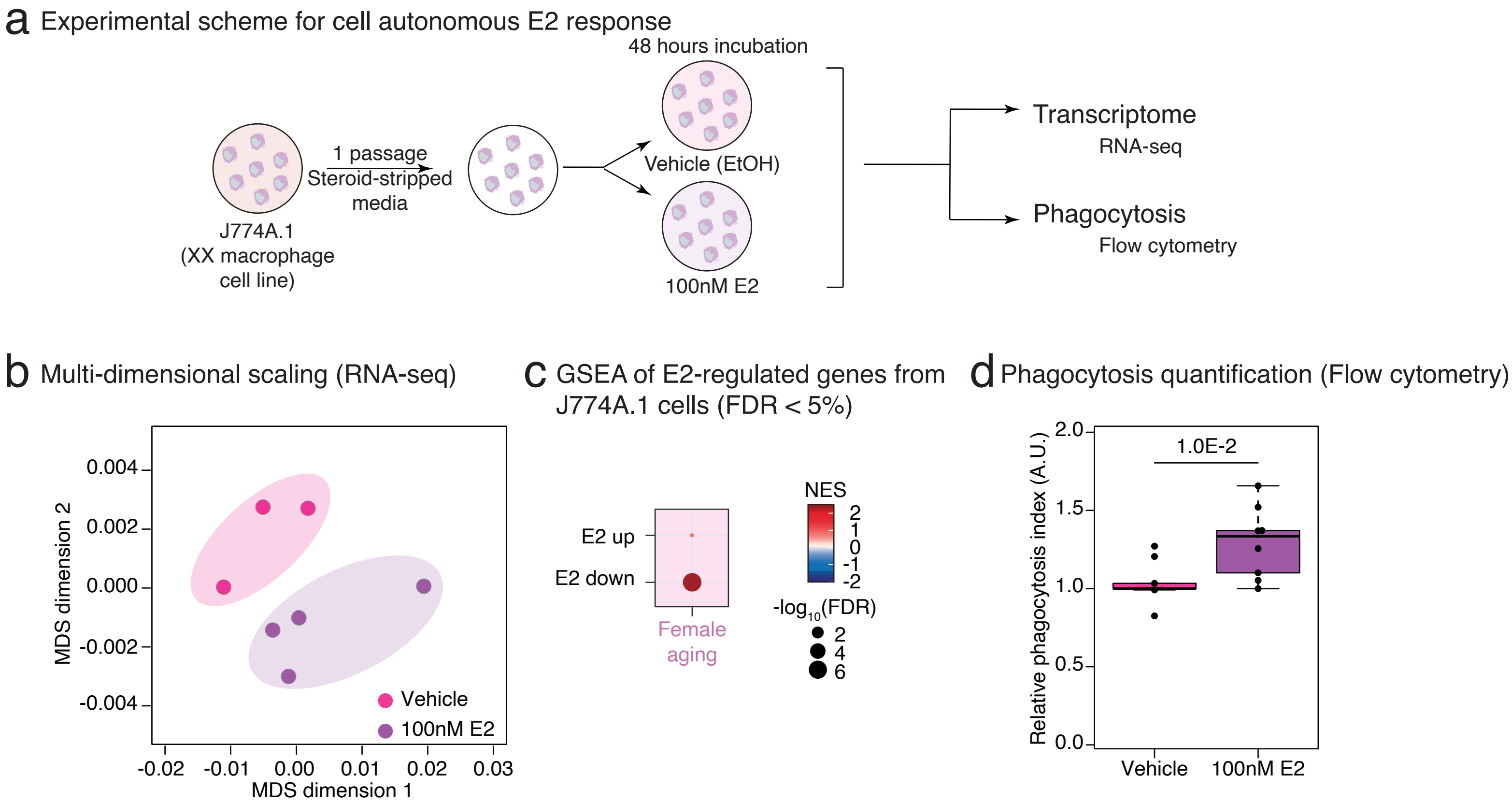

#### Extended Data Figure 15

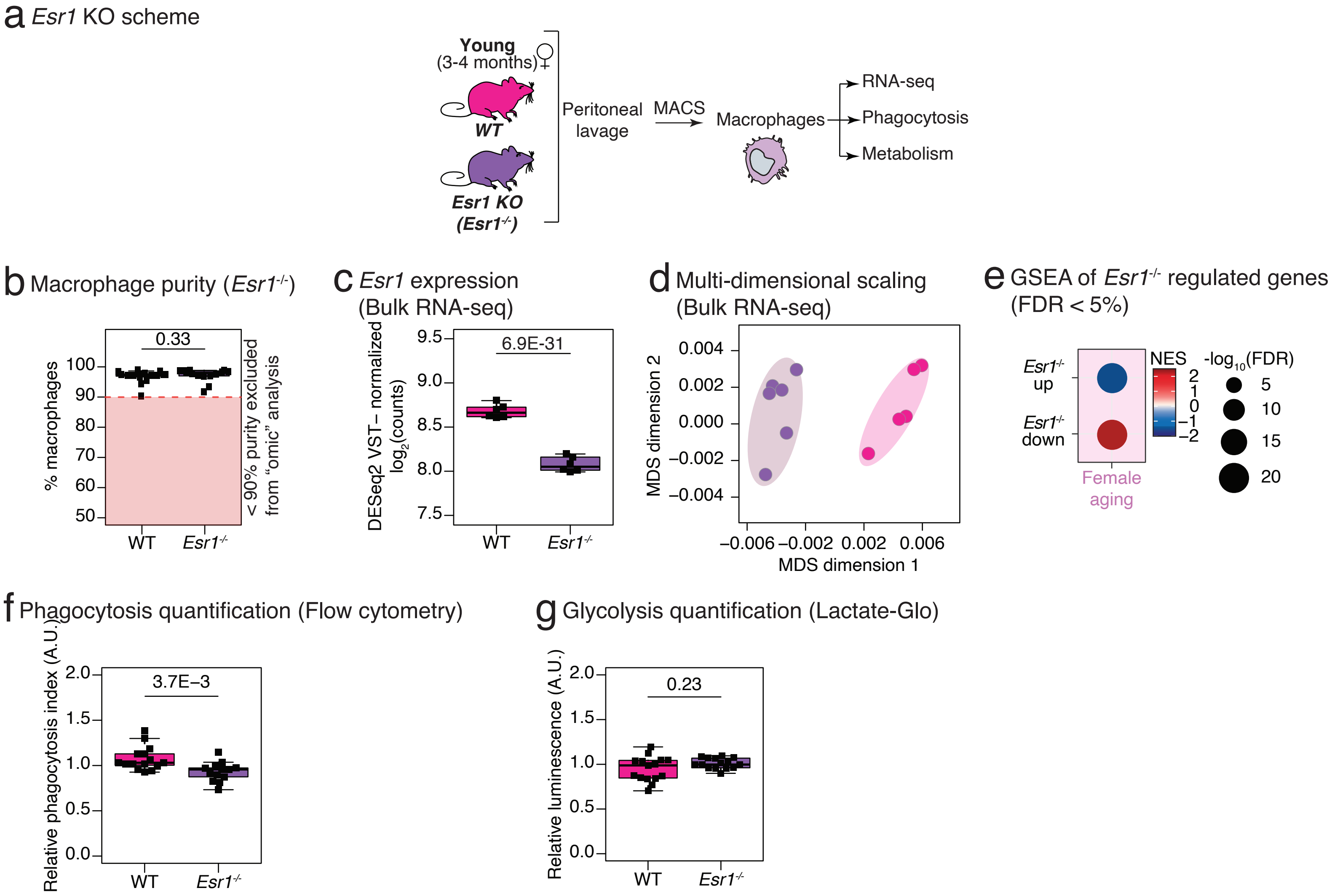

#### Extended Data Figure 16

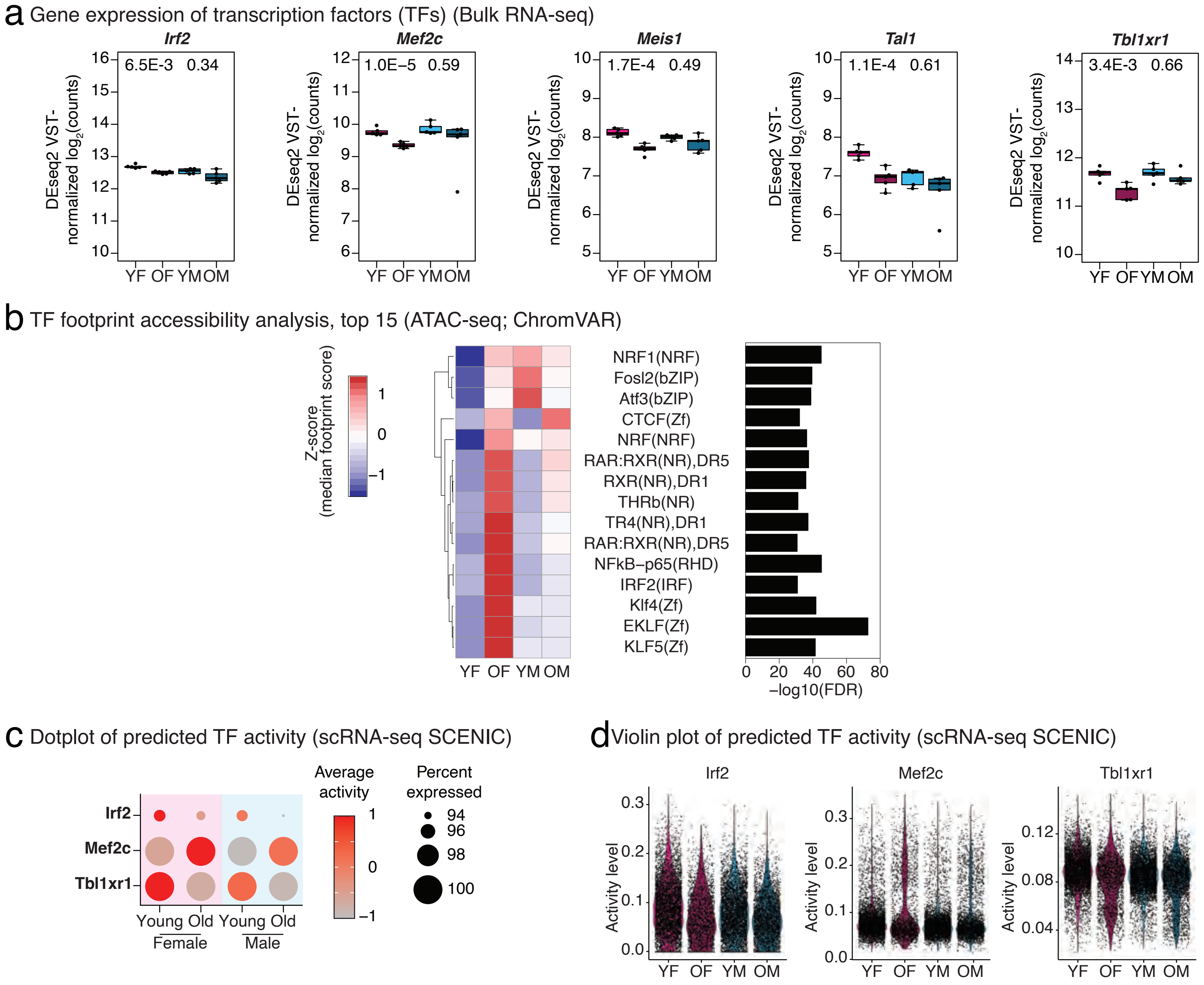

#### Extended Data Figure 17

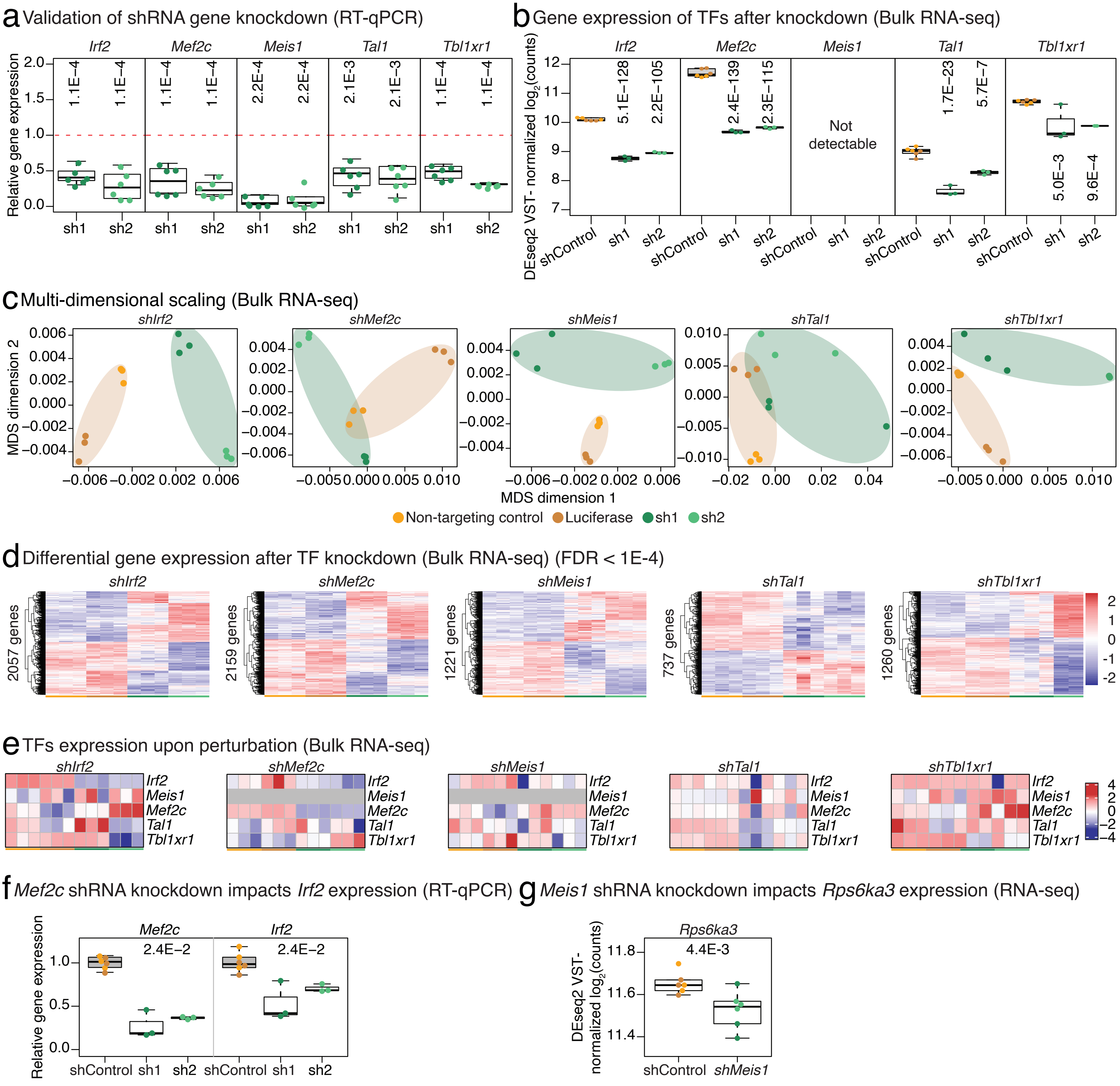

**Extended Data Figure 18**

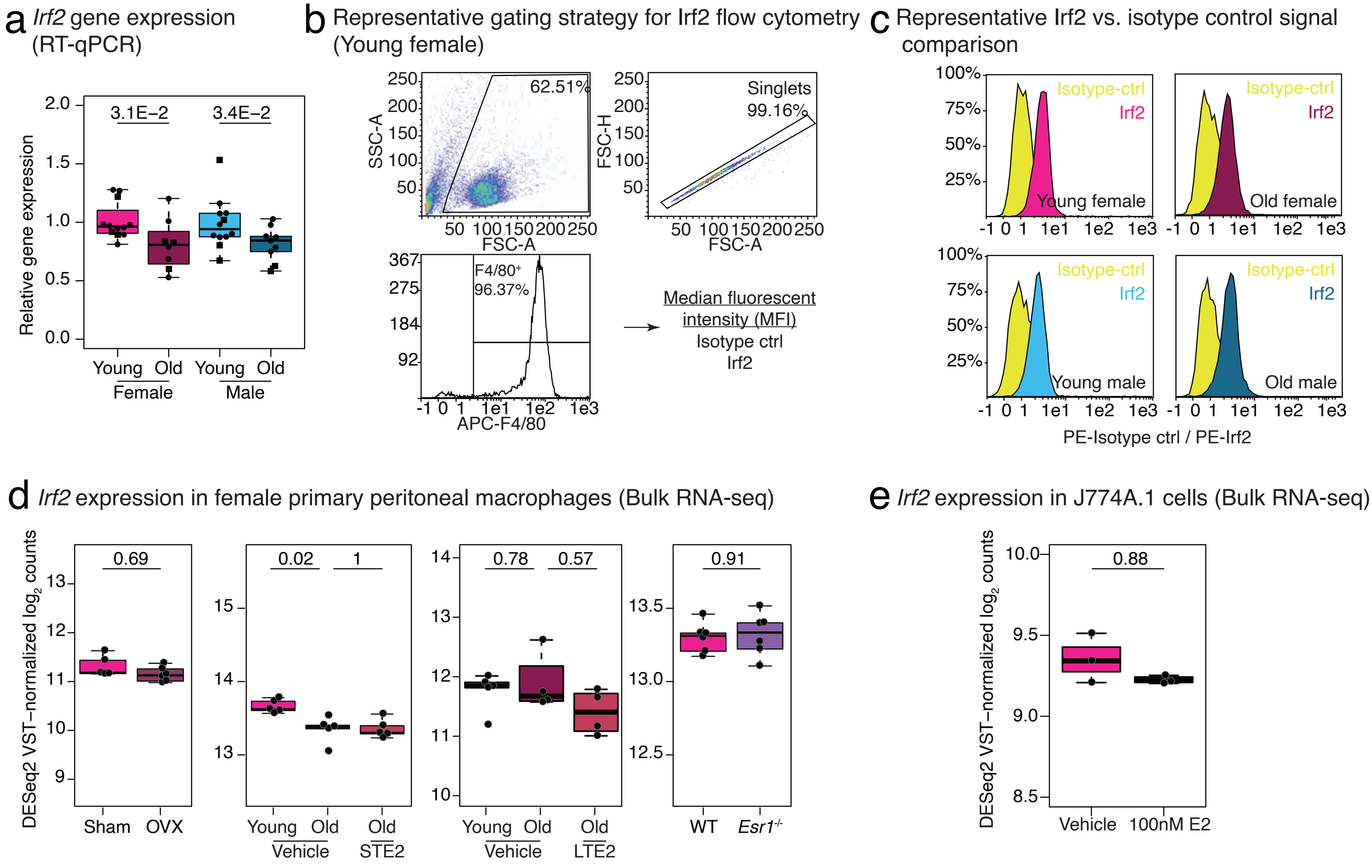

**Extended Data Figure 19**

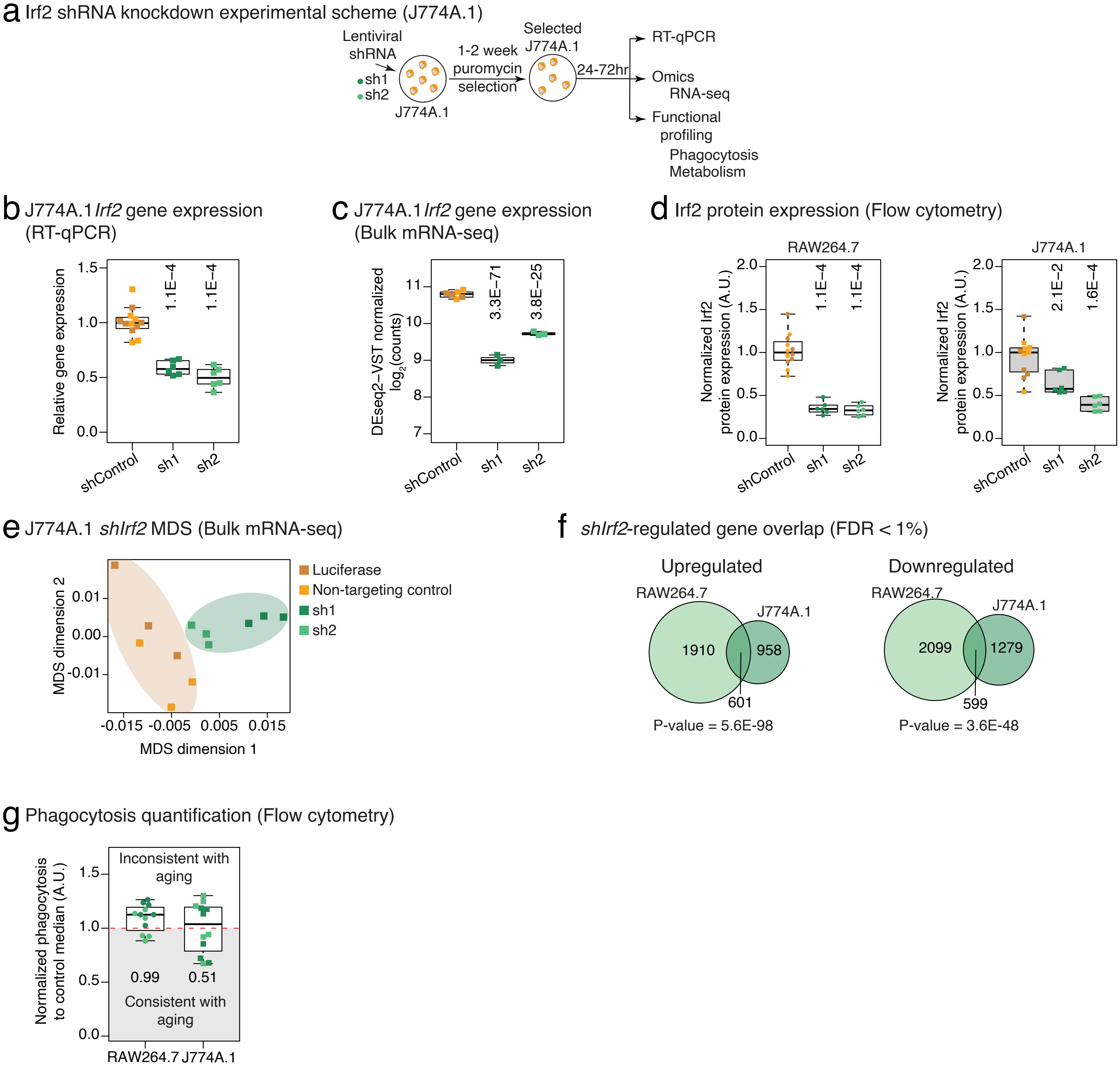

Extended Data Figure 20

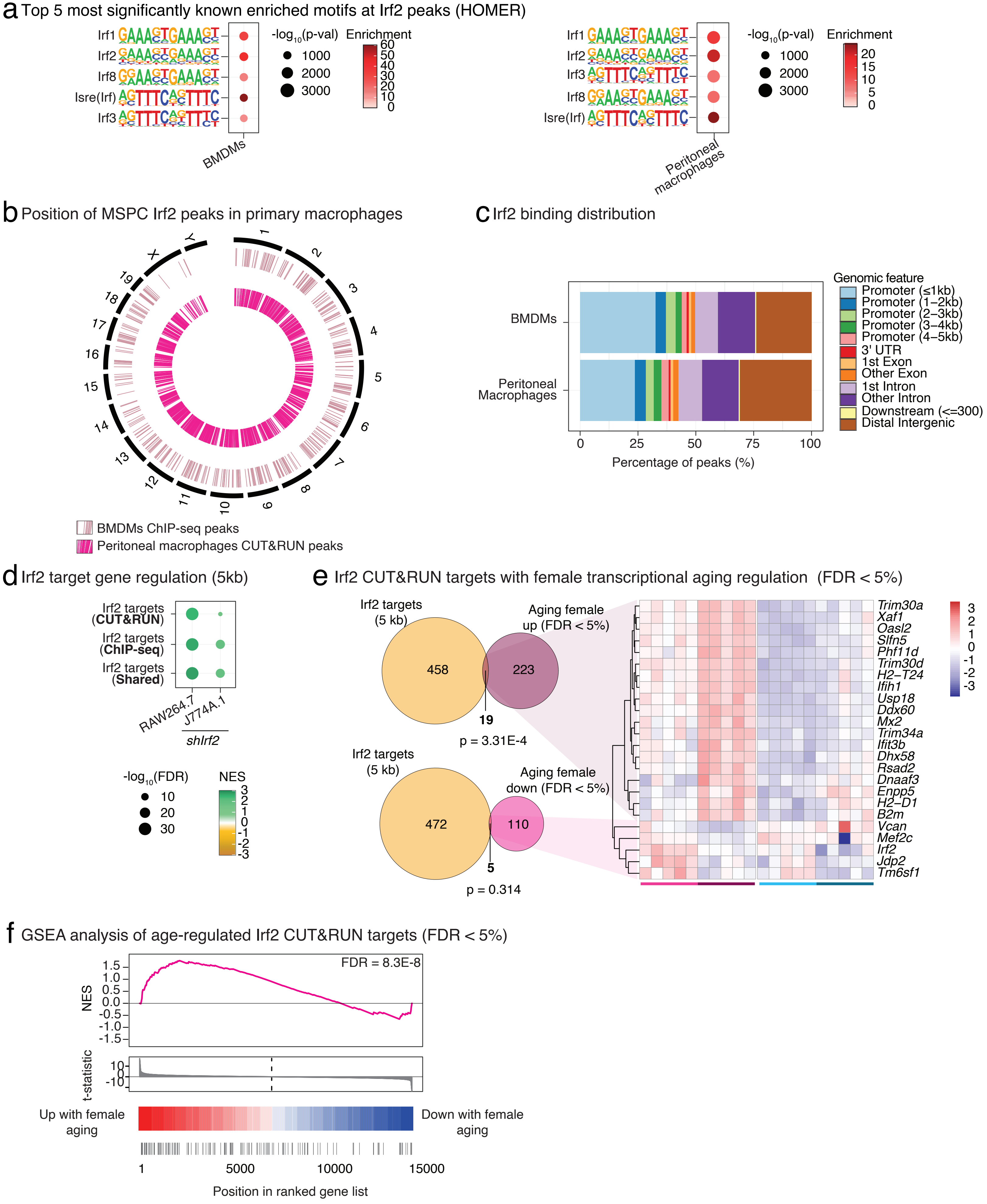

#### Extended Data Figure 21

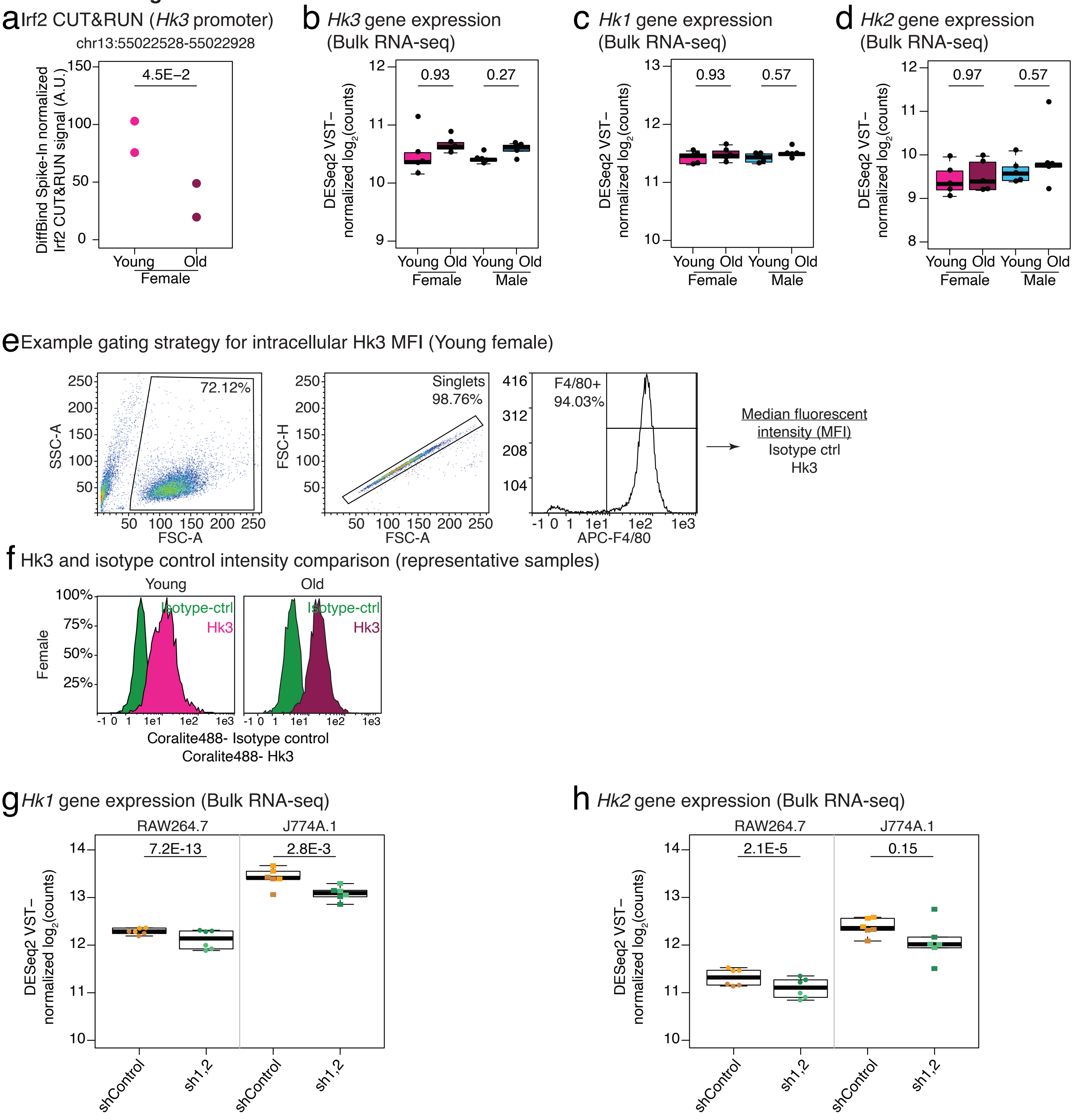
